# Spatial organization and mitigation of autofluorescence in multiplexed spatial proteomics of aged fresh-frozen human brain

**DOI:** 10.64898/2026.09.11.750056

**Authors:** Iva Sutevski, Luochen Guo, Niclas Branzell, Per E. Andrén, Per Svenningsson, Burcu Ayoglu

## Abstract

Multiplexed imaging technologies are transforming the study of human tissue biology, but their application to the aged brain is hindered by autofluorescence, particularly in fresh-frozen specimens. Here, we characterized autofluorescence across four brain regions from 21 donors and found broad spectral emission, regional and gray-white matter differences, and an association with donor age. Photobleaching conditions adopted from formalin-fixed paraffin-embedded tissue caused marked region- and compartment-dependent damage in fresh-frozen sections. We therefore developed a Tris–EDTA-supplemented photobleaching workflow that reduced autofluorescence by 58–70% while preserving tissue architecture and cellular content. We established a custom 28-plex DNA-barcoded antibody panel targeting neuronal, glial, immune, and vascular markers, providing a resource for fresh-frozen human brain. Integration of the optimized photobleaching workflow with this panel enabled spatial proteomic analysis across fresh-frozen brain regions. By co-registering pre-photobleaching autofluorescence with multiplexed protein maps, we further established a cellular-resolution framework for spatial characterization of autofluorescence. This revealed region-dependent protein marker relationships and preferential enrichment of autofluorescent particles near nuclei and within microglial and CD68-positive regions. Together, this work establishes a practical workflow for multiplexed spatial proteomics in fresh-frozen brain and characterizes autofluorescence as both a technical confound and a spatially structured feature of the aged human brain.

## INTRODUCTION

Multiplexed imaging technologies are rapidly transforming the study of tissue biology by enabling spatially resolved analysis of dozens of protein targets at single-cell resolution while preserving tissue architecture ^1,2^. Their application to human specimens is particularly valuable because it provides direct access to disease-relevant cellular states and interactions that cannot be fully recapitulated in model systems.

In the human brain, multiplexed imaging is enabling detailed spatial characterization of cellular composition and neuropathology across neurological and neurodegenerative conditions. However, high-plex antibody panels and workflows reported to date have predominantly been developed for formalin-fixed paraffin-embedded (FFPE) tissue ^3–7^. Fresh-frozen human brain tissue, despite its availability in biobanks spanning diverse neurological diseases, remains largely unexplored for high-plex spatial proteomic, limiting its use for spatial studies of brain biology, aging, and disease.

A major barrier to multiplexed imaging of aged fresh-frozen human brain is intrinsic autofluorescence arising from endogenous fluorophores, including lipofuscin and other age-associated lipopigments that accumulate over time ^8^. Their broad emission overlaps with reporter fluorophores used for immunostaining and may mimic marker-positive structures, impair cell segmentation, confound marker interpretation, and bias downstream single-cell quantification, particularly in highly multiplexed imaging workflows ^8,9^. Although autofluorescence is widely recognized as a source of imaging artefacts, little is known about its regional, cellular, and molecular organization in aged fresh-frozen human brain.

Conventional strategies for autofluorescence suppression primarily rely on chemical quenching applied after tissue staining. Although effective in low-plex settings ^10^, post-staining quenching approaches are incompatible with high-plex imaging workflows requiring repeated cycles of antibody staining or reporter hybridization, imaging and signal removal, particularly when these processes are fully automated. Computational approaches for *post hoc* autofluorescence correction have also been proposed ^11^, but may not fully recover signal fidelity in highly autofluorescent tissues.

Photobleaching offers an attractive alternative by irreversibly suppressing endogenous fluorophores prior to antibody staining without introducing residual quenching reagents. However, photobleaching protocols developed primarily for FFPE tissue ^7,12^ are not necessarily transferable to fresh-frozen human brain. The intrinsic fragility of fresh-frozen sections, together with elevated iron ^13^ and other redox-active species in aged brain may increase susceptibility to oxidative damage during photobleaching. Achieving effective autofluorescence suppression while preserving tissue integrity therefore represents a major challenge for multiplexed imaging of fresh-frozen brain. In parallel, the absence of openly specified high-plex antibody panels tailored to the fresh-frozen human brain further limits the broader adoption of multiplexed imaging in this tissue type.

Here, we systematically investigate autofluorescence in aged fresh-frozen human brain and address the technical barriers it poses to multiplexed imaging. We establish an optimized photobleaching strategy that suppresses autofluorescence while preserving tissue integrity and develop a fully defined 28-plex DNA-barcoded antibody panel tailored to fresh-frozen human brain, providing a high-plex resource for this tissue preparation. By integrating pre-photobleaching autofluorescence with multiplexed protein maps, we introduce a high-plex cellular-resolution framework for interrogating autofluorescence itself. This enables analysis of its spatial organization and protein-marker relationships across brain regions, cellular compartments, and cell types. Together, these advances provide both a practical workflow for multiplexed spatial proteomics in fresh-frozen human brain and a new analytical approach to study autofluorescence as a spatially organized tissue feature.

## RESULTS

### Autofluorescence in aged fresh-frozen human brain is widespread, spectrally broad, and varies by region, compartment, and donor age

To enable multiplexed spatial proteomics in the fresh-frozen human brain, we first sought to characterize the extent and impact of autofluorescence and evaluate strategies for its suppression. Autofluorescence was assessed across the 488, 550 and 647 nm fluorescence imaging channels in fresh-frozen human brain tissue from a total of 21 donors aged 59-97 years **(Supplementary Table 1)**, encompassing samples from the caudate-putamen, hippocampus, parietal cortex, and prefrontal cortex **(Supplementary Table 2)**. Autofluorescence was detected in all three channels and across all four brain regions, with the highest signal intensities observed in the 488 nm channel, followed by 550 nm and 647 nm channels **(Figure 1A)**.

**Figure 1.**
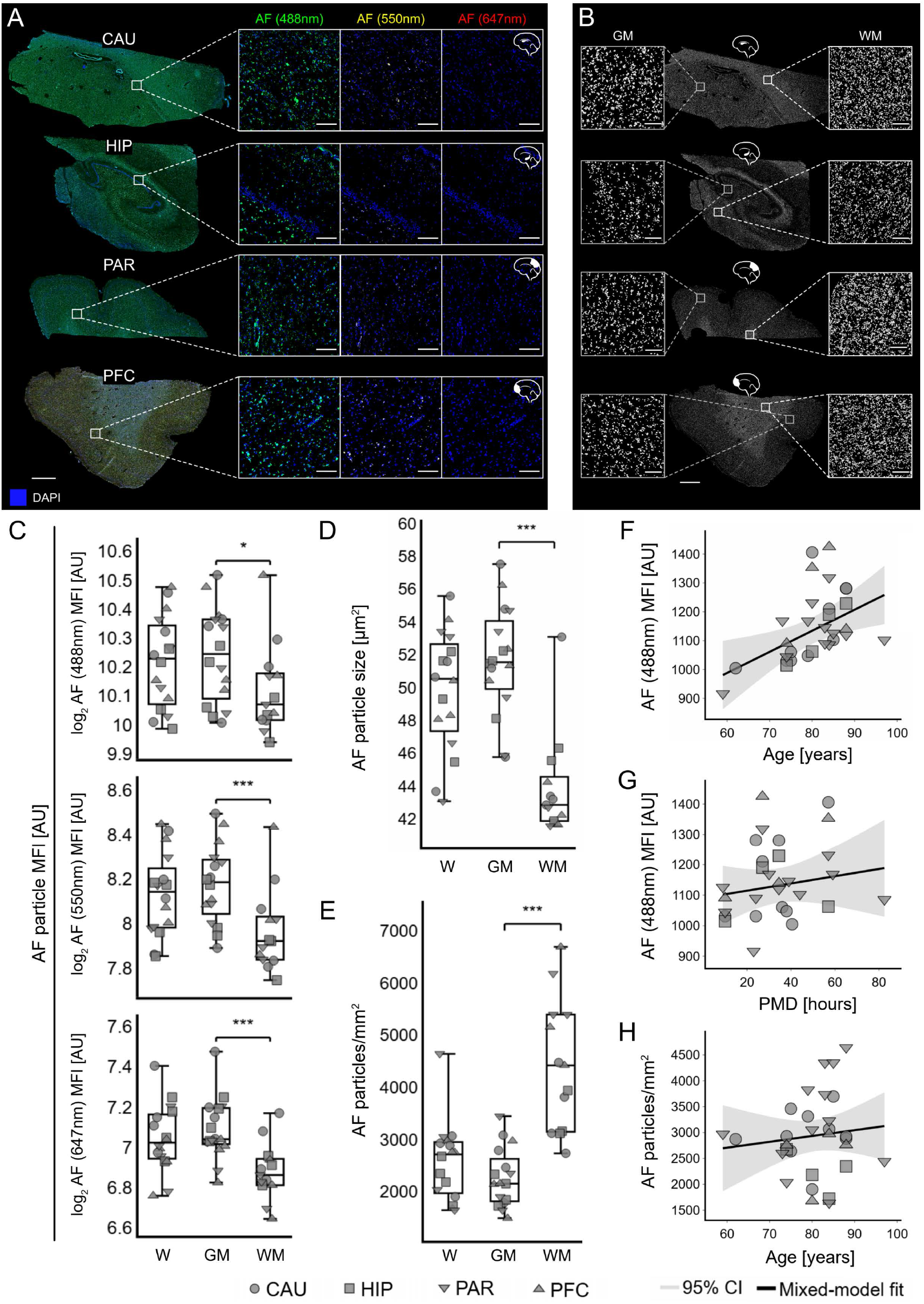
Spectral, regional, and compartmental differences in autofluorescence across aged fresh-frozen human brain tissue. **A.** Representative autofluorescence patterns acquired in the 488 nm, 550 nm, and 647 nm fluorescence microscopy imaging channels across four regions of aged human brain: caudate-putamen (CAU), hippocampus (HIP), parietal cortex (PAR), and prefrontal cortex (PFC). Scale bars: 2 mm (whole section images), 150 µm (magnified regions). **B.** Representative autofluorescence particle masks across the four brain regions, with magnified views illustrating distinct gray matter (GM) and white matter (WM) patterns. Scale bars: 2 mm (whole section images), 150 µm (magnified regions). **C.** Boxplots showing the mean autofluorescence intensity (MFI in arbitrary units, AU) in the 488 nm, 550 nm, and 647 nm channels across whole tissue sections (W), gray matter only (GM), and white matter only (WM). Each point represents an individual sample, and point shape indicates brain region. Mixed-effects models adjusted for brain region, with donor as a random intercept, showed higher MFI in GM than WM at 488 nm (GM/WM ratio = 1.08 [95% CI, 1.02–1.14], *p* = 0.013), 550 nm (1.14 [95% CI, 1.07–1.22], *p* = 1.18 × 10⁻⁴), and 647 nm (1.15 [95% CI, 1.09–1.22], *p* = 2.14 × 10⁻⁷). Sample size: *n* (CAU) = 4, *n* (HIP) = 4, *n* (PAR) = 4, *n* (PFC) = 4. **D-E.** Boxplots showing the mean autofluorescent particle size **(D)** and particle density **(E)** in the 488 nm channel across whole sections (W), gray matter (GM) only and white matter (WM) only. Each point represents an individual sample, and point shape indicates brain region. Mixed-effects models adjusted for brain region, with donor as a random intercept, showed larger particles in GM than WM (mean difference = 7.86 µm^2^ [95% CI, 5.96–9.76], *p* = 5.97 × 10⁻^16^) **(D)**, but lower particle density in GM (mean difference = -2189 particles/mm^2^, [95% CI, -2815.3 to -1562.9], *p* = 7.32 × 10⁻¹²) **(E)**. Sample size: *n* (CAU) = 4, *n* (HIP) = 4, *n* (PAR) = 4, *n* (PFC) = 4. **F-G.** Association between mean autofluorescent particle intensity in the 488 nm channel with donor age **(F)** and post-mortem delay (PMD) **(G)**. Mixed-effects models adjusted for brain region, with donor as a random intercept, showed increasing MFI with donor age (β = 7.31 AU/year [95% CI, 2.05–12.56], *p* = 0.0065) **(F)**, but no significant association with PMD (β = 1.18 AU/hour [95% CI, -1.89 to 4.26], *p* = 0.45) **(G)**. Sample size: *n*(CAU) = 10, *n* (HIP) = 4, *n* (PAR) = 12, *n* (PFC) = 4. **H.** Association between autofluorescent particle density and donor age. Mixed-effects model adjusted for brain region, with donor as a random intercept, showed no significant association (β = 11.25 particles/mm^2^/year [95% CI, −24.39 to 46.89], *p* = 0.54). Sample size: *n* (CAU) = 10, *n* (HIP) = 4, *n* (PAR) = 12, *n* (PFC) = 4.

To quantify the size, density and mean signal intensity of autofluorescent particles, segmentation masks of autofluorescent particles were generated based on the 488 nm channel **(Figure 1B)**. In total, approximately 2.7 million autofluorescent particles were analyzed, including ∼1.8 million particles from gray matter and ∼0.9 million particles from white matter across all four examined brain regions **(Supplementary Table 3)**. These masks recapitulated tissue architecture, distinguishing gray from white matter **(Figure 1B)**. Across all three channels, autofluorescence intensity was significantly higher in gray matter than in white matter **(Figure 1C)**, with the most pronounced gray-white matter differences observed in the parietal cortex and hippocampus **(Supplementary Figure 1)**. Among the regions, the caudate-putamen exhibited the highest autofluorescence signal intensity across all channels, while the prefrontal cortex also showed comparatively elevated levels in the two channels with higher autofluorescence **(Supplementary Figure 1)**. Analysis of particle-level features revealed that autofluorescent structures were significantly smaller **(Figure 1D)** but more densely distributed in white matter compared to gray matter **(Figure 1E)**, with particularly pronounced differences in the parietal and prefrontal cortex **(Supplementary Figure 2)**, suggesting gray and white matter might contain biologically distinct autofluorescence sources.

Autofluorescence intensity in the 488 nm channel showed a positive correlation with donor age across all regions and tissue compartments analyzed **(Figure 1F)**, with the strongest associations observed in the hippocampus **(Supplementary Figure 3A)**. No significant correlation was observed between post-mortem delay and autofluorescence intensity in the 488 nm channel **(Figure 1G; Supplementary Figure 3B)**. Donor age and post-mortem delay did not significantly associate with autofluorescent particle density **(Figure 1H; Supplementary Figure 5)** or particle size (**Supplementary Figure 4)**.

Together, these findings establish autofluorescence as a prominent, regionally and compartmentally variable feature of aged fresh-frozen human brain, and highlight the need for effective suppression for multiplexed imaging. Because photobleaching-based approaches are compatible with automated cyclic imaging and widely used for autofluorescence suppression in FFPE tissue, we next tested its suitability for fresh-frozen human brain.

### Photobleaching induces region- and compartment-dependent tissue damage that is mitigated by EDTA without compromising autofluorescence removal

We first tested a commonly used photobleaching strategy ^7,14^ based on combined chemical and light-mediated bleaching of endogenous fluorophores. Fresh-frozen caudate-putamen sections were photobleached and tissue integrity was assessed by hematoxylin and eosin (H&E) staining **(Figure 2A)**. Compared to non-photobleaching consecutive sections, photobleached samples exhibited substantial, readily visible tissue damage **(Figure 2A–B)**. Immunostaining of the photobleached caudate-putamen sections for the neuronal markers tubulin beta 3 class III (TUBB3) and synaptophysin (SYP) further revealed that tissue damage was compartment-dependent, with gray matter being markedly more affected than white matter **(Figure 2B)**. Because the initial experiments used pre-sectioned tissue and standard fixation, we repeated photobleaching on freshly sectioned samples using stronger fixation. However, neither modification prevented photobleaching-induced tissue damage **(Supplementary Figure 6)**.

**Figure 2.**
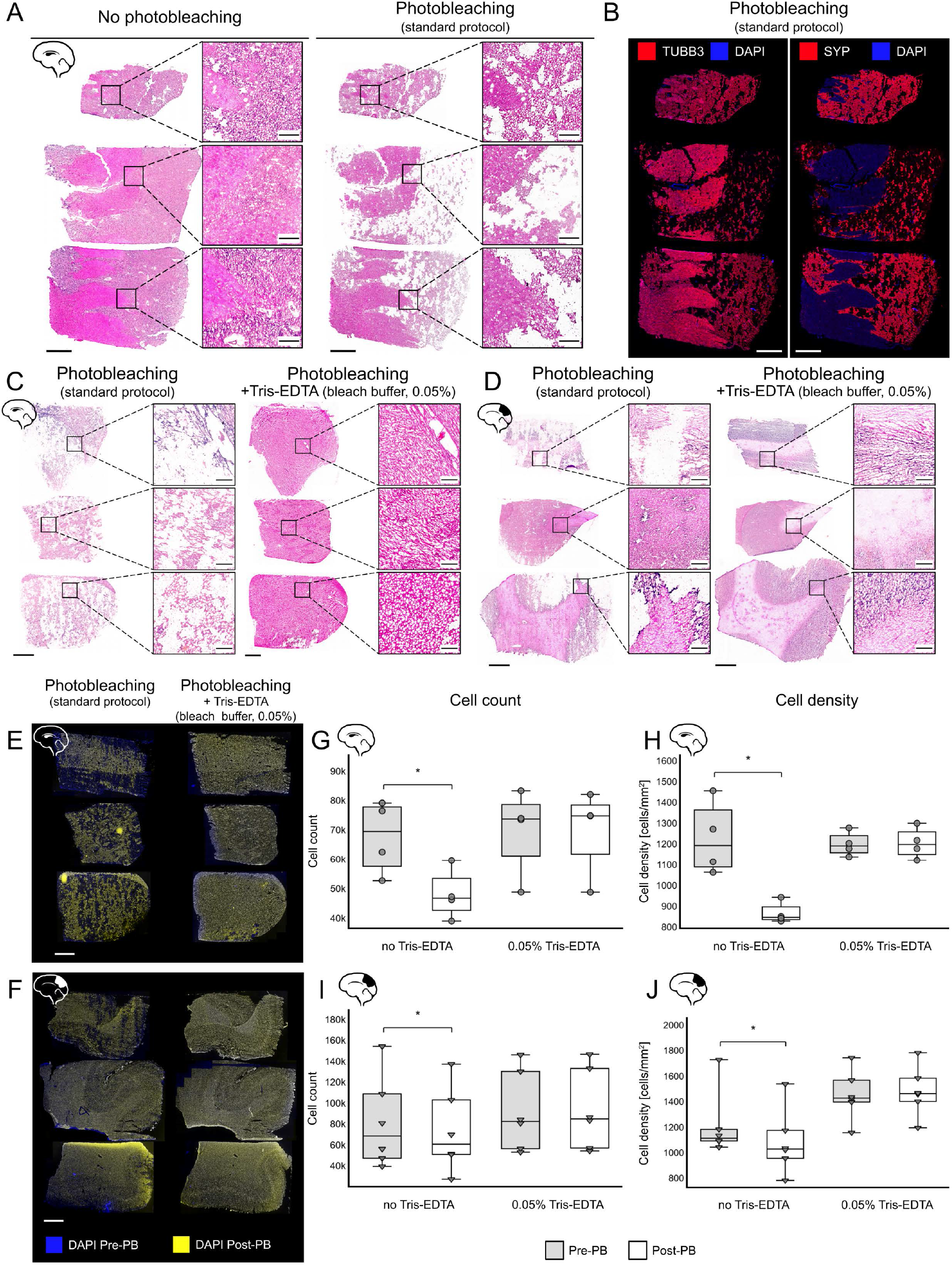
Standard photobleaching damages fresh-frozen human brain tissue, while Tris-EDTA supplementation preserves tissue integrity. **A.** Representative hematoxylin and eosin (H&E)-stained fresh-frozen caudate-putamen sections not exposed to photobleaching (left) and consecutive sections exposed to photobleaching (right) using the standard photobleaching protocol, illustrating substantial photobleaching-induced tissue damage. Whole-section images and corresponding magnified regions are shown. **B.** Representative immunofluorescence staining of photobleached caudate-putamen sections for tubulin III (TUBB3, red) and synaptophysin (SYP, red), with DAPI nuclear staining (blue) demonstrating that tissue damage predominantly affects gray matter. **C-D.** Representative H&E-stained caudate-putamen **(C)** and parietal cortex **(D)** sections following photobleaching without Tris-EDTA (left), and with 0.05% Tris-EDTA (1:100 dilution) (right) added during the first of two photobleaching incubations. Whole-section images and corresponding magnified regions are shown. **E-F.** Spatially aligned DAPI images acquired before (Pre-PB, blue) and after photobleaching (Post-PB, yellow) from caudate-putamen **(E)** and parietal cortex **(F)**, without (top) or with 0.05% Tris-EDTA (bottom) added to the photobleaching buffer. The without- and with-Tris-EDTA conditions were performed on consecutive sections. Pre- and post-photobleaching images were acquired from the same section. **G–H**. Cell count **(G)** and density **(H)** before (Pre-PB) and after photobleaching (Post-PB) in caudate-putamen. Without Tris-EDTA, both cell count and density were significantly reduced (mean change:−19,666 cells, *p* = 0.011; −360 cells/mm², *p* = 0.012), whereas no significant reductions were observed with 0.05% Tris-EDTA. Sample size: *n* (CAU)= 4. **I-J.** Cell count **(I)** and density **(J)** before (Pre-PB) and after photobleaching (Post-PB) in parietal cortex. Without Tris-EDTA, both cell count and density were significantly reduced (−7,819 cells, p = 0.022; −127 cells/mm², p = 0.043), whereas no reductions were observed with 0.05% Tris-EDTA. Sample size: *n* (PAR)= 6. One-sided paired *t*-tests were used to test for post-photobleaching reductions in cell count and density. **Scale bars:** 2 mm (whole section images), 300 µm (magnified regions).

We next tested whether metal ion chelation could mitigate the oxidative damage arising during photobleaching, thereby reducing tissue damage, by supplementing the wash or photobleaching buffers with the chelating agent ethylenediaminetetraacetic acid (EDTA) **(Supplementary Figure 7)**. Supplementation of the photobleaching buffer with an EDTA-based formulation (5% Tris, 5% tetrasodium EDTA, 1.5% 2-butoxyethanol), hereafter referred to as Tris-EDTA, markedly preserved tissue integrity in both caudate-putamen and parietal cortex samples **(Figure 2C-D; Supplementary Figure 7)**. Comparable protective effects were observed with both 0.025% and 0.05% Tris-EDTA supplementation **(Supplementary Figure 8)**.

We quantified this protective effect by spatially aligning DAPI images acquired before and after photobleaching in caudate-putamen and parietal-cortex sections **(Figure 2E-F; Supplementary Figure 9)**. Comparison of aligned images confirmed substantial photobleaching-induced cell loss in the absence of Tris-EDTA supplementation. On average, photobleaching without Tris-EDTA reduced cell count and density by 28% in caudate-putamen **(Figure 2G-H)** and 10% in parietal cortex **(Figure 2I-J)**. In contrast, both 0.025% and 0.05% Tris-EDTA effectively preserved cellular content, with no detectable loss in cell count or cell density following photobleaching (**Supplementary Figure 9)**. Because both concentrations provided comparable protection, 0.05% Tris-EDTA was selected for subsequent experiments to provide the greatest margin of protection across brain regions.

Having established that Tris-EDTA preserved tissue integrity during photobleaching, we next determined whether this protection came at the expense of autofluorescence removal. Importantly, autofluorescence reduction remained substantial across all four brain regions and three imaging channels **(Figure 3A-B).** On average, autofluorescence intensity decreased by approximately 58% in the highly autofluorescent 488 nm channel, 70% in the 550 nm channel, and 63% in the 647 nm channel **(Figure 3C)**, with highly consistent reductions observed across caudate-putamen, hippocampus, parietal cortex, and prefrontal cortex samples **(Figure 3C)**.

**Figure 3.**
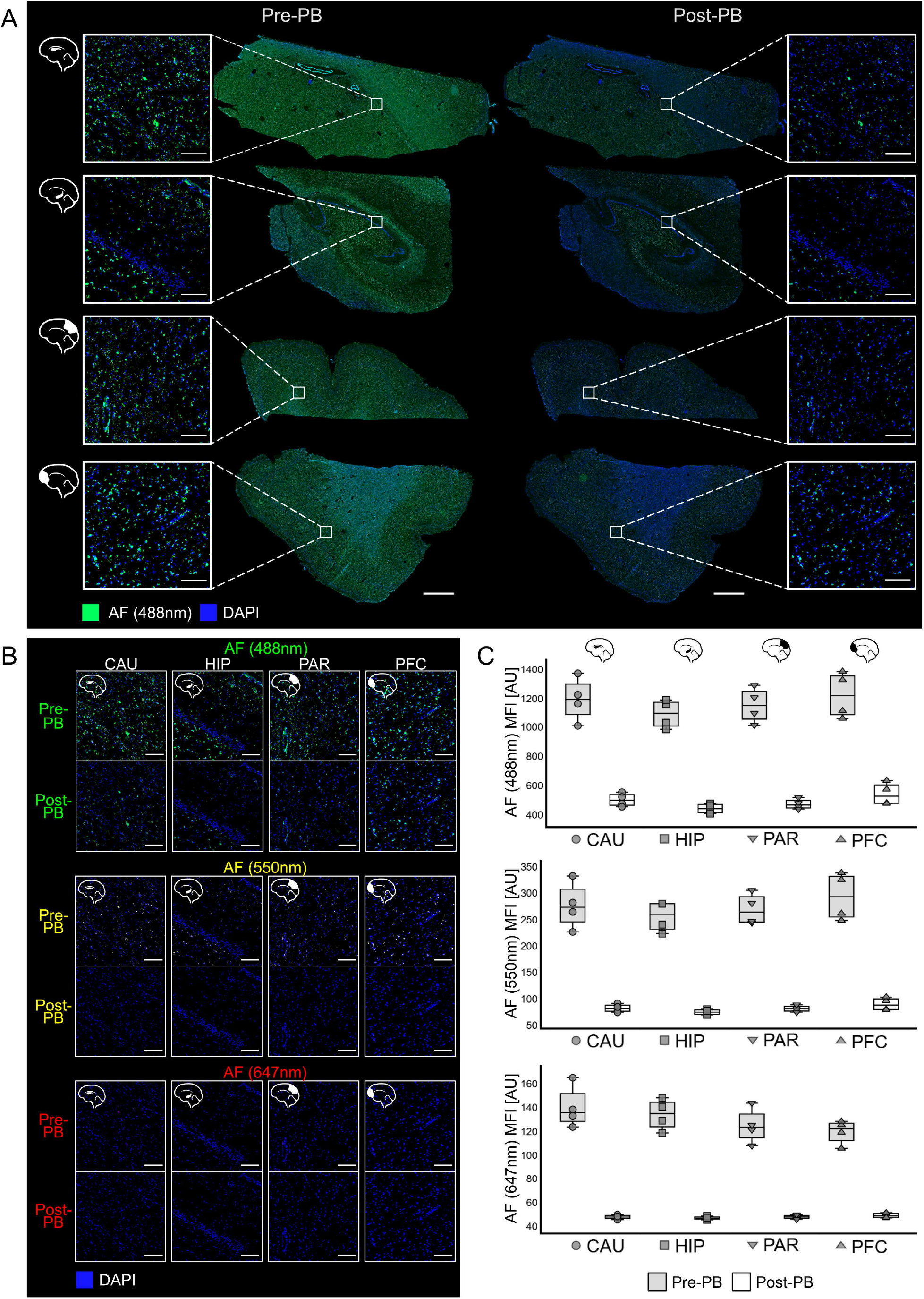
Optimized photobleaching protocol with 0.05% Tris-EDTA significantly reduces autofluorescence across brain regions and fluorescence channels. **A.** Overview of autofluorescence signal before (Pre-PB) and after photobleaching (Post-PB) in the 488 nm channel across four brain regions, demonstrating robust autofluorescence reduction at the whole-section level. Scale bars: 2 mm (whole-section images), 150 µm (magnified regions). **B.** Representative images showing autofluorescence signal before and after photobleaching with Tris-EDTA across the four analyzed brain regions and three fluorescence channels. Photobleaching with Tris-EDTA resulted in a marked reduction of autofluorescence signal in all tested channels and brain regions. Scale bars: 150 µm. **C.** Boxplots showing mean autofluorescence intensity (MFI in arbitrary units, AU) before and after photobleaching with Tris-EDTA across the four analyzed brain regions and three fluorescence channels. Optimized photobleaching significantly reduced autofluorescence intensity across all three channels and all four brain regions (one-sided paired *t*-tests, all *p* ≤ 0.001). Sample size: *n* (CAU) = 4, *n* (HIP) = 4, *n* (PAR) = 4, *n* (PFC) = 4.

Together, these results demonstrate that standard photobleaching causes substantial, region- and compartment-dependent damage in fresh-frozen human brain, with pronounced susceptibility in gray matter and deep subcortical regions such as the caudate-putamen. Tris-EDTA supplementation preserved tissue integrity during photobleaching without compromising autofluorescence removal, enabling an optimized photobleaching workflow for fresh-frozen brain **(Supplementary Figure 10)** compatible with downstream multiplexed immunofluorescence imaging **(Supplementary Figures 11–12)**.

### Development of a 28-plex antibody panel for fresh-frozen human brain

Having established an optimized autofluorescence-suppression workflow compatible with fresh-frozen brain, we next developed a DNA-barcoded antibody panel for high-dimensional, single-cell resolved spatial proteomic analysis. A total of 64 antibodies targeting neuronal, glial, vascular, immune and pathology-associated proteins were evaluated for inclusion in a DNA-barcoded multiplexed imaging panel **(Supplementary Table 5)**. Of these, 44 antibodies were incorporated into a preliminary custom panel **(Supplementary Tables 5 and 7)**. Following evaluation of staining quality, signal intensity range, and compatibility with the multiplexed workflow, 28 of these antibodies were selected to form the final optimized panel for fresh-frozen human brain tissue **(Figure 4; Supplementary Table 6)**.

**Figure 4.**
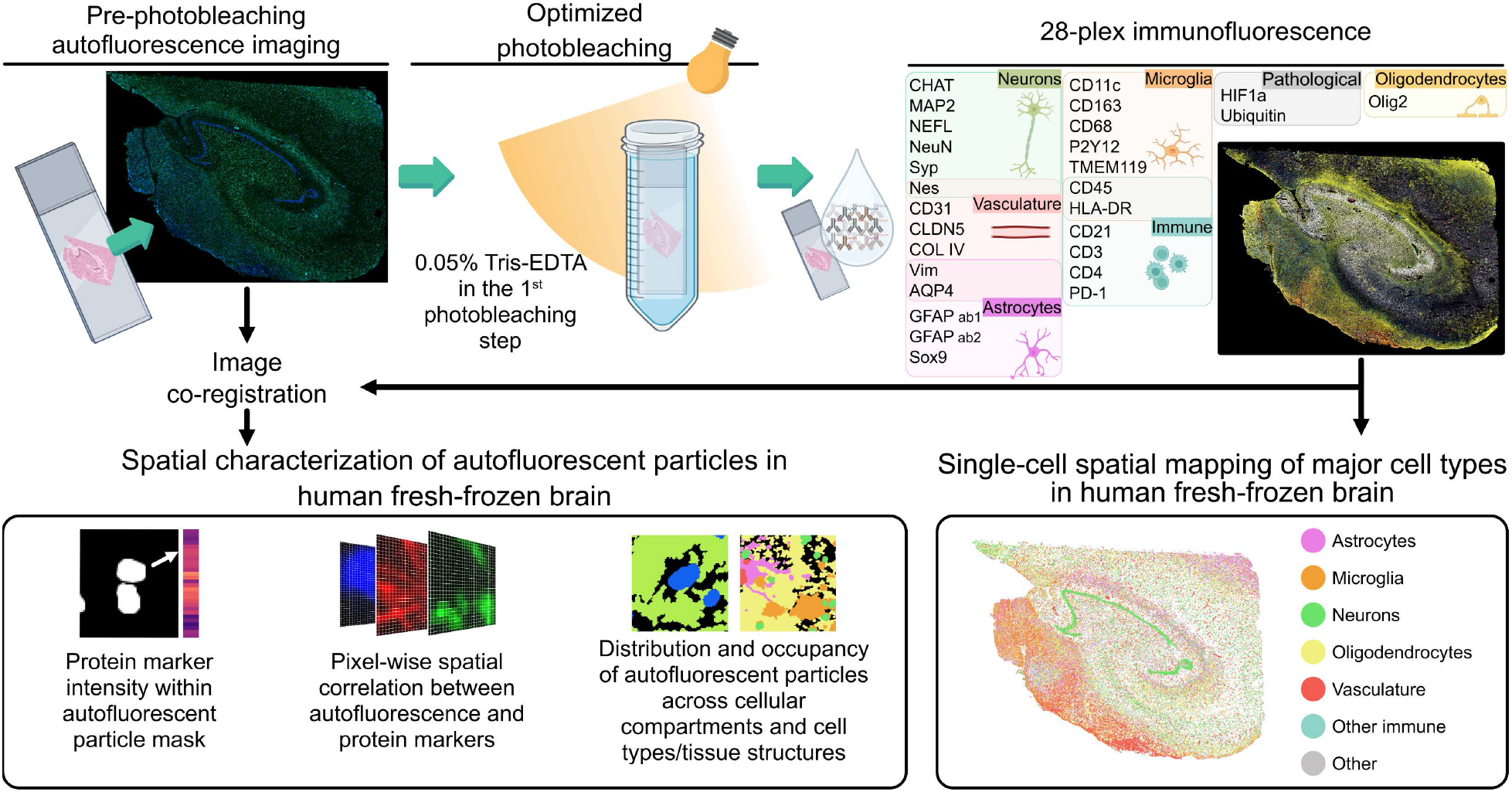
Optimized photobleaching workflow enabling multiplexed immunofluorescence imaging and cell-resolved characterization of autofluorescence in fresh-frozen human brain tissue. Fresh-frozen human brain sections were stained with DAPI and imaged to capture native autofluorescence before undergoing optimized photobleaching, in which the first bleaching solution was supplemented with 0.05% Tris–EDTA to preserve tissue integrity. Tissue sections were subsequently stained with a custom 28-plex DNA-barcoded antibody panel targeting major neuronal, glial, vascular, immune, and pathology-associated proteins and imaged using automated cyclic multiplexed immunofluorescence. The resulting datasets enabled single-cell resolved spatial proteomic analysis across four brain regions, while co-registration of pre-photobleaching autofluorescence images with multiplexed immunofluorescence data enabled molecular and spatial characterization of autofluorescent particles. The latter included quantification of mean fluorescence intensity of all 28 protein markers within autofluorescent particles, pixel-wise spatial correlation between autofluorescence and protein markers, and analyses of autofluorescent particle distribution and occupancy across cellular compartments and cell types/tissue structures. (Parts of the figure created in BioRender. Ayoglu, B. (2026) BioRender.com)

The final panel included markers for neurons (NeuN, ChAT, MAP2, SYP, NEFL), astrocytes (AQP4, GFAP-1, GFAP-2, SOX9, VIM), microglia (CD68, CD11c, CD163, P2Y12, CD45, HLA-DR, TMEM119), other immune cell populations (CD3, CD4, CD21, CD45, HLA-DR, PD1), oligodendrocytes (OLIG2), endothelial cells (CD31, CLDN5, COLIV, NES, VIM), and neuropathology-associated markers (HIF1a, Ubiquitin) **(Figure 4)**. Only ten antibodies, predominantly targeting immune cell markers, were available in pre-barcoded form; the remaining antibodies required in-house conjugation and validation **(Supplementary Table 5)**. Together, these efforts established an optimized 28-plex panel for multiplexed imaging of fresh-frozen human brain tissue.

### Optimized photobleaching enables single-cell resolved spatial proteomics in fresh-frozen human brain tissue

We next combined the optimized photobleaching workflow with the 28-plex panel in an automated cyclic imaging workflow across all four brain regions. Residual autofluorescence, most prominent in the 488 nm channel, was captured during an initial blank imaging cycle (cycle 0). This blank-cycle signal was used for background subtraction during initial image processing, a step commonly supported by software accompanying automated multiplexed imaging platforms. As shown for representative markers including CD45, MAP2, and GFAP, this effectively removed residual background while revealing marker-specific staining patterns in subsequent imaging cycles **(Figure 5)**.

**Figure 5.**
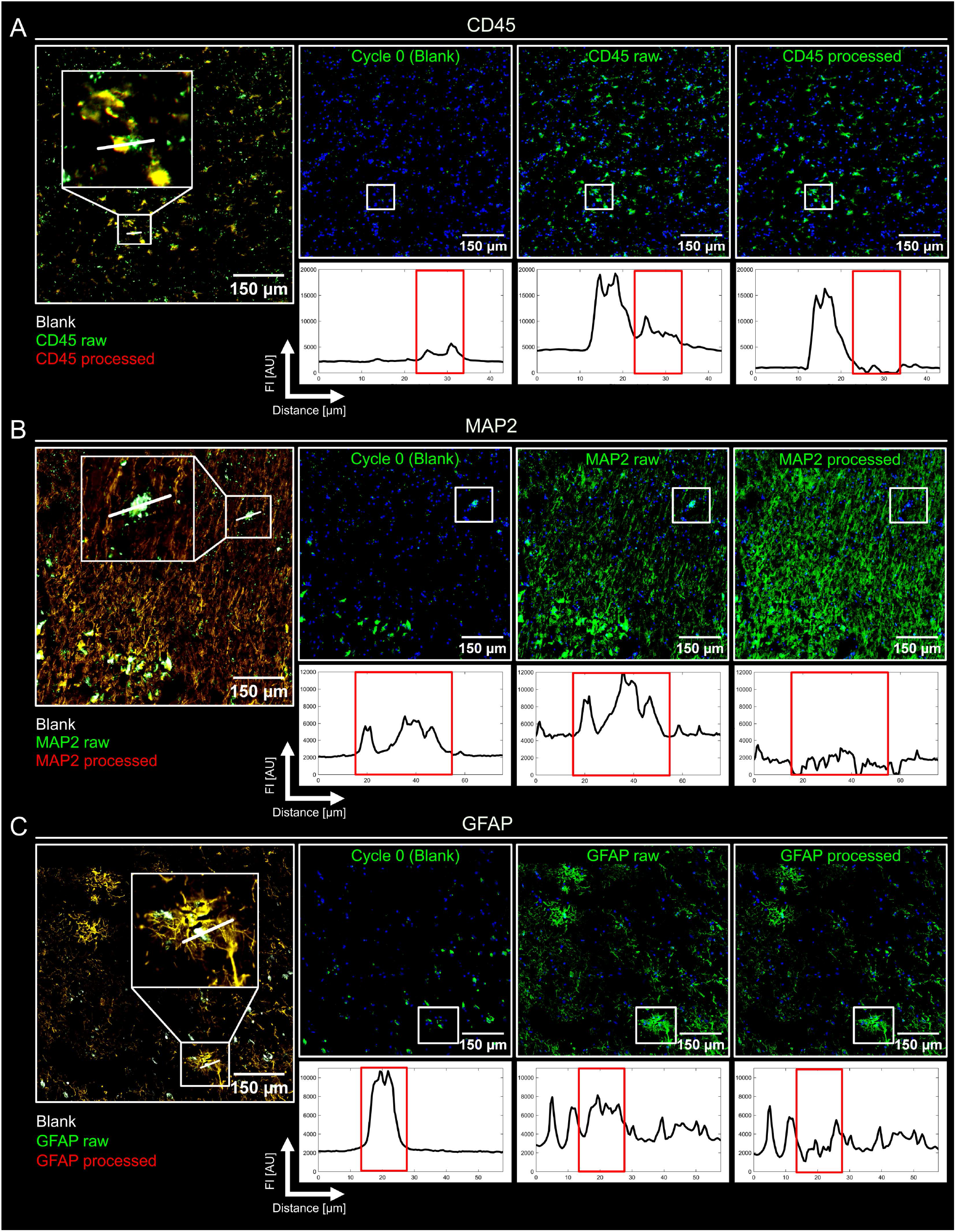
Residual autofluorescence after optimized photobleaching and its removal by blank-cycle background subtraction. Residual autofluorescence remaining after optimized photobleaching was evaluated in the 488 nm imaging channel for three representative markers: CD45 (immune cells including microglia) **(A)**, MAP2 (neurons) **(B)**; and GFAP (astrocytes) **(C)**. In each panel, the leftmost image shows an overlay of residual autofluorescence captured in the blank-cycle (white), raw marker signal (green), and processed marker signal after blank-cycle subtraction (red), illustrating the spatial overlap between residual autofluorescence and raw marker signal prior to correction. The three images to the right show, from left to right, the blank-cycle image (cycle 0) acquired before antibody staining to capture residual autofluorescence, the raw marker image, and the processed marker image following blank-cycle subtraction. Fluorescence intensity (FI) profiles measured along the indicated white line segments are shown below the corresponding images. Blank-cycle subtraction effectively removed residual autofluorescence while preserving marker-specific staining, demonstrating that residual autofluorescence remaining after optimized photobleaching can be effectively corrected during image processing.

Having established that residual autofluorescence could be effectively controlled within an automated multiplexed workflow, the resulting imaging datasets were subjected to single-cell segmentation, feature extraction and cell type annotation **(Figure 6)**. Cellular composition was analyzed across four representative samples, one from each brain region, corresponding to approximately 407,000 segmented cells in total, including ∼130,000 from caudate-putamen, ∼107,000 from hippocampus, ∼57,000 from parietal cortex, and ∼114,000 from prefrontal cortex **(Figure 6E–F)**. Using threshold-based annotation rules, neurons, oligodendrocytes, astrocytes, microglia and other immune cells, as well as endothelial cells, could be identified and spatially mapped **(Figure 6E)**, enabling annotation of approximately 89% of all segmented cells across the four brain regions **(Figure 6F)**. Distinct cell populations resolved by multiplexed marker profiles showed spatial distributions consistent with known neuroanatomical, regional, and compartmental organization of the human brain **(Figure 6E–F; Supplementary Figure 13)**. This analysis confirmed that the optimized workflow and 28-plex panel support single-cell-resolved spatial proteomic analysis of fresh-frozen human brain tissue.

**Figure 6.**
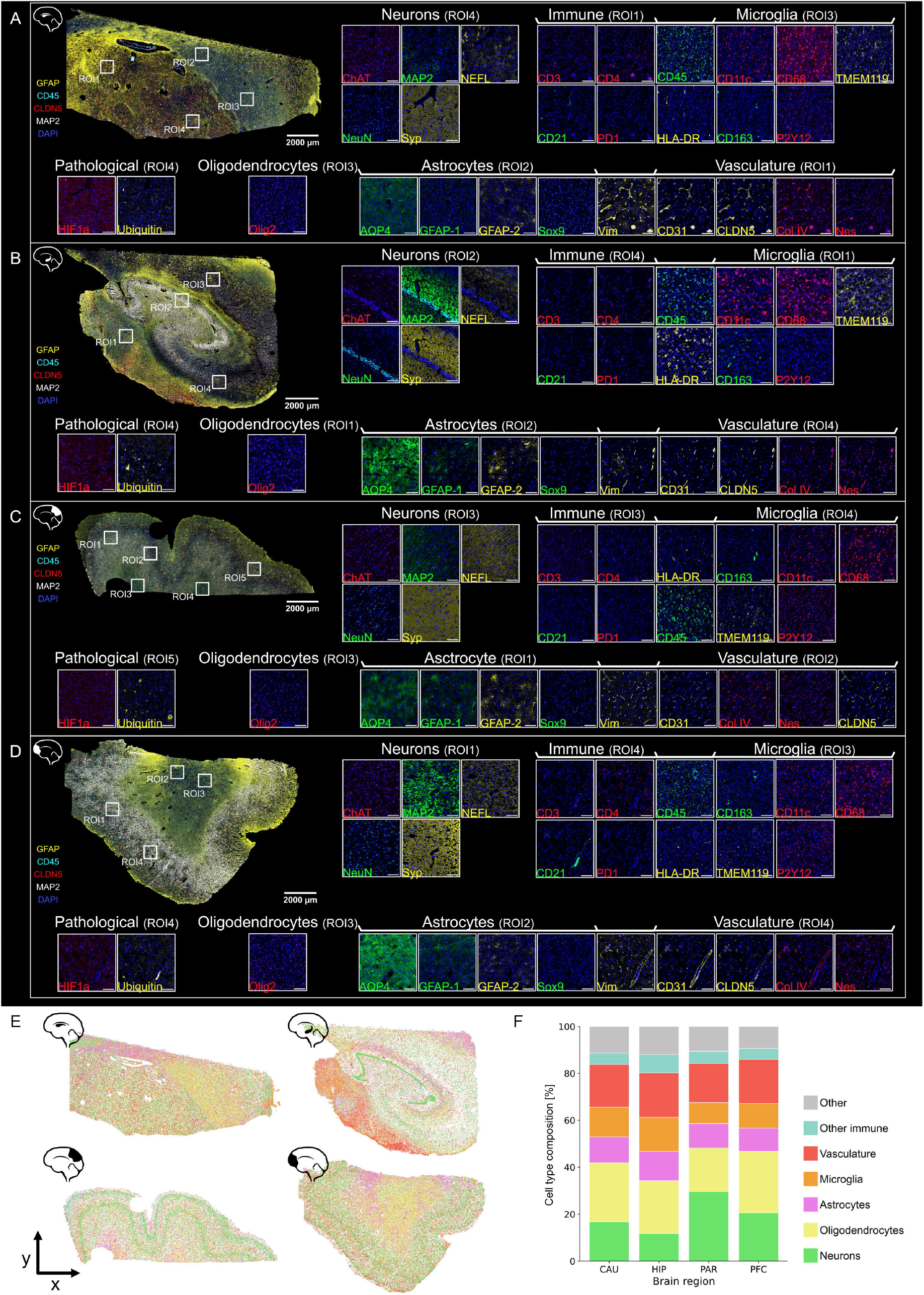
Application of the 28-plex DNA-barcoded antibody panel developed for multiplexed spatial proteomic imaging of fresh-frozen human brain tissue. **A-D.** Representative multiplexed images are shown for the caudate-putamen **(A)**, hippocampus **(B)**, parietal cortex **(C)**, and prefrontal cortex **(D)**. Whole-section images provide an overview of tissue architecture together with selected markers highlighting major cellular and structural features: GFAP (yellow), CD45 (cyan), CLDN5 (red), MAP2 (white), and DAPI (blue). Magnified regions display the complete 28-plex antibody panel, organized by major cellular or molecular targets, including neurons (NeuN, ChAT, MAP2, SYP, NEFL), astrocytes (AQP4, GFAP-1, GFAP-2, SOX9, VIM), microglia (CD68, CD11c, CD163, P2Y12, CD45, HLA-DR, TMEM119), other immune cells (CD3, CD4, CD21, CD45, HLA-DR, PD1), oligodendrocytes (OLIG2), vasculature (CD31, CLDN5, collagen IV, NES, VIM), and pathology-associated markers (HIF1α, ubiquitin). Individual antibody staining patterns are displayed using the corresponding fluorescence detection channel (green for 488 nm, yellow for 550 nm, and red for 647 nm), together with DAPI nuclear staining (blue). Detailed antibody information is provided in **Supplementary Tables 6–7**. Scale bars: 2 mm (whole section images), 150 µm (magnified regions). **E.** Spatial maps showing the localization of annotated cell types in representative samples from each analyzed brain region. Each dot represents an individual cell and is colored according to its assigned cell type: astrocytes in magenta, microglia in orange, neurons in green, oligodendrocytes in yellow, immune cells in cyan, vascular cells in red, and other/unidentified cells in gray. **F.** Stacked bar chart summarizing the relative cell-type composition of the representative samples from each brain region. Sample size: *n* (CAU) = 1, *n* (HIP) = 1, *n* (PAR) = 1, *n* (PFC) = 1.

### Autofluorescence overlaps with protein marker signals in a region-dependent manner

We next co-registered pre-photobleaching 488 nm autofluorescence with post-photobleaching 28-plex images to examine spatial relationships between autofluorescent particles and protein-marker signals (**Figure 4 and 7A**). Aligned autofluorescence and multiplexed protein marker images revealed substantial overlap between autofluorescence and several brain cell type and structural markers, particularly those with fine-granular staining patterns **(Figure 7B)**, indicating that these markers would be substantially confounded without adequate autofluorescence suppression.

**Figure 7.**
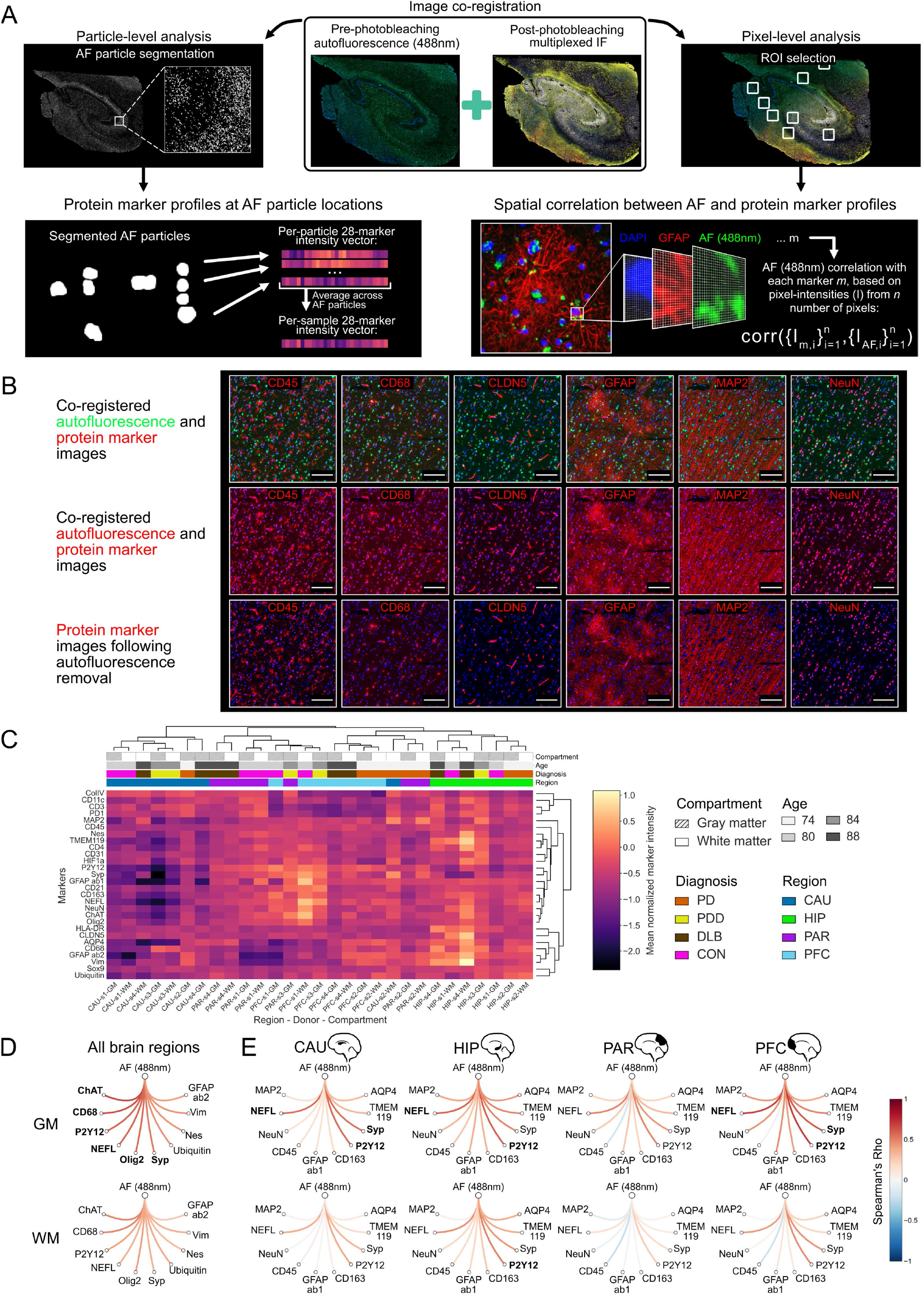
Molecular and spatial relationships between autofluorescence and protein marker expression in aged fresh-frozen human brain. **A.** Workflow for analysis of molecular and spatial relationships between autofluorescence and multiplexed protein expression. Autofluorescence images acquired in the 488 nm channel before antibody staining were computationally aligned (co-registered) with multiplexed immunofluorescence images acquired following optimized photobleaching and 28-plex immunostaining. Two complementary analyses were then performed. First, autofluorescent particle masks were generated from the 488 nm autofluorescence images, and mean protein marker intensities were quantified within the segmented autofluorescent particles across the 28-plex antibody panel. Second, representative regions of interest (ROIs) were selected from gray and white matter of each brain region (four ROIs from gray matter and up to four ROIs from white matter per sample), and pixel-wise Spearman correlations were calculated between 488 nm autofluorescence intensity and the intensity of each protein marker. **B.** Co-registered autofluorescence and multiplexed protein marker images illustrate the potential confounding effect of autofluorescence on marker interpretation. Representative co-registered images are shown for CD45 (immune cells), CD68 (microglia), CLDN5 (endothelial cells), GFAP (astrocytes), MAP2 and NeuN (neurons). The top row shows co-registered pre-photobleaching autofluorescence (green) and protein marker staining (red). The middle row shows the same co-registered images, with autofluorescence displayed in the protein marker color (red), illustrating the marker signal that would be observed if autofluorescence were not experimentally removed. The bottom row shows the corresponding protein marker images acquired following optimized photobleaching and multiplexed imaging, demonstrating recovery of marker-specific staining patterns after autofluorescence removal. Scale bars: 150 µm. **C.** Protein-marker profiles associated with autofluorescent particles across brain regions and tissue compartments. Mean marker intensities measured within segmented autofluorescent particle masks are shown as a hierarchically clustered heatmap. Columns represent individual samples and rows represent markers from the 28-plex antibody panel. Sample annotations indicate tissue compartment, donor age, diagnosis and brain region. Colors represent log₁₀-transformed, normalized mean marker intensities measured within autofluorescent particle masks. **D.** Protein markers showing the strongest overall pixel-wise spatial correlation with autofluorescence. The ten markers exhibiting the strongest Spearman correlation with 488 nm autofluorescence intensity are shown for gray (top) and white matter (bottom), with correlation coefficients averaged across the four analyzed brain regions. Correlation strength is indicated by the color scale. Markers with correlation coefficients greater than 0.5 are highlighted in bold. **E.** Region- and compartment-specific variation in pixel-wise spatial correlation between autofluorescence and protein marker expression. The ten markers showing the greatest variation in Spearman correlation with 488 nm autofluorescence intensity across brain regions and tissue compartments are shown for caudate-putamen, hippocampus, parietal cortex, and prefrontal cortex (from left to right), separately for gray (top) and white matter (bottom). Correlation strength is indicated by the color scale. Markers with correlation coefficients greater than 0.5 or lower than −0.5 are highlighted in bold.

To characterize protein marker signals at autofluorescent particle locations, mean intensities of all 28 protein markers were then quantified within autofluorescent particle masks generated from the pre-photobleaching 488 nm images. Marker intensities within autofluorescent particle masks varied substantially across brain regions and compartments and were generally higher in white than gray matter **(Figure 7C; Supplementary Figures 14–15; Supplementary Table 8)**, with GFAP, P2Y12, NEFL and COLIV showing the highest mean intensities within autofluorescent particle masks in both compartments **(Supplementary Figure 15)**. Region-specific differences were also prominent. Regardless of tissue compartment, caudate-putamen showed the lowest overall marker intensities within autofluorescent particle masks, whereas hippocampus showed the highest **(Figure 7C; Supplementary Figure 14; Supplementary Table 8)**. In hippocampus, CD68, CLDN5, GFAP, TMEM119 and VIM displayed the strongest signals within autofluorescent particle masks, whereas both cortical regions showed relatively high ChAT, NEFL, P2Y12 and SYP signals **(Supplementary Table 8)**. Hierarchical clustering based on these intensities segregated samples primarily by brain region, with comparatively little contribution from tissue compartment, donor age, or diagnosis **(Figure 7C; Supplementary Figure 16)**. Thus, this analysis suggests that anatomical context is the major determinant of the protein marker milieu at autofluorescent particle locations.

### Autofluorescence shows selective spatial co-variation with protein marker signals across brain regions

While the preceding analysis examined protein marker signals at autofluorescent particle locations, we next asked which protein marker signals spatially co-varied with autofluorescence intensity across the broader tissue environment. Multiple representative 1,500 pixel × 1,500 pixel regions of interest (ROIs) were selected from gray and white matter in each brain region **(Figure 7A; Supplementary Table 4)**, and pixel-wise correlations were calculated between autofluorescence intensity in the 488 nm channel and the intensity of all 28 protein markers within each ROI.

Pixel-wise spatial correlations between autofluorescence and protein marker signals were generally stronger in gray matter than in white matter **(Supplementary Figure 17)**. When correlations were averaged across brain regions, the strongest correlations with autofluorescence were observed for ChAT, CD68, P2Y12, NEFL, and OLIG2 **(Figure 7D)**. Region-specific patterns were further examined among markers showing the greatest variation in correlation across brain regions and tissue compartments. Parietal cortex showed the weakest overall pixel-wise correlations between autofluorescence and protein marker signals with strong correlations observed only for CD68 and ChAT **(Figure 7E; Supplementary Table 9)**, whereas hippocampus and prefrontal cortex showed stronger correlations for additional markers including NES and ubiquitin **(Supplementary Table 9)**. Overall, these findings indicate that autofluorescence does not co-vary uniformly across protein markers but shows selective spatial relationships with a subset of neuronal and glial markers, with distinct patterns across brain regions.

### Autofluorescent particles show distinct spatial organization across cellular compartments and tissue structures

Because these protein marker relationships could partly reflect regional differences in tissue composition, we next examined whether autofluorescent particles preferentially localized to specific cellular compartments, cell types, and tissue structures. Using the same ROIs, masks for cellular compartments (nuclei, cytoplasm, and extracellular space) and cell types/tissue structures (astrocytic, microglial, neuronal, neurofilament-associated, and vascular) were generated from the co-registered 28-plex imaging data using combinatorial thresholding and marker exclusion criteria **(Figure 8A; Supplementary Figure 18)**. Within these ROIs, autofluorescent particles occupied approximately 4–6% of tissue area (**Supplementary Figure 19)**. We assessed their spatial organization using complementary distribution, enrichment, and occupancy analyses **(Figure 8B, F, I, M)**. Distribution analysis measured the fraction of the autofluorescence overlapping each cellular compartment or cell type/tissue structure mask, enrichment analysis accounted for the relative abundance of that mask within the ROI **(Figure 8B, I)**, and occupancy analysis measured the fraction of each cellular compartment or cell type/tissue structure mask occupied by autofluorescence relative to areas outside it **(Figure 8F, M)**.

**Figure 8.**
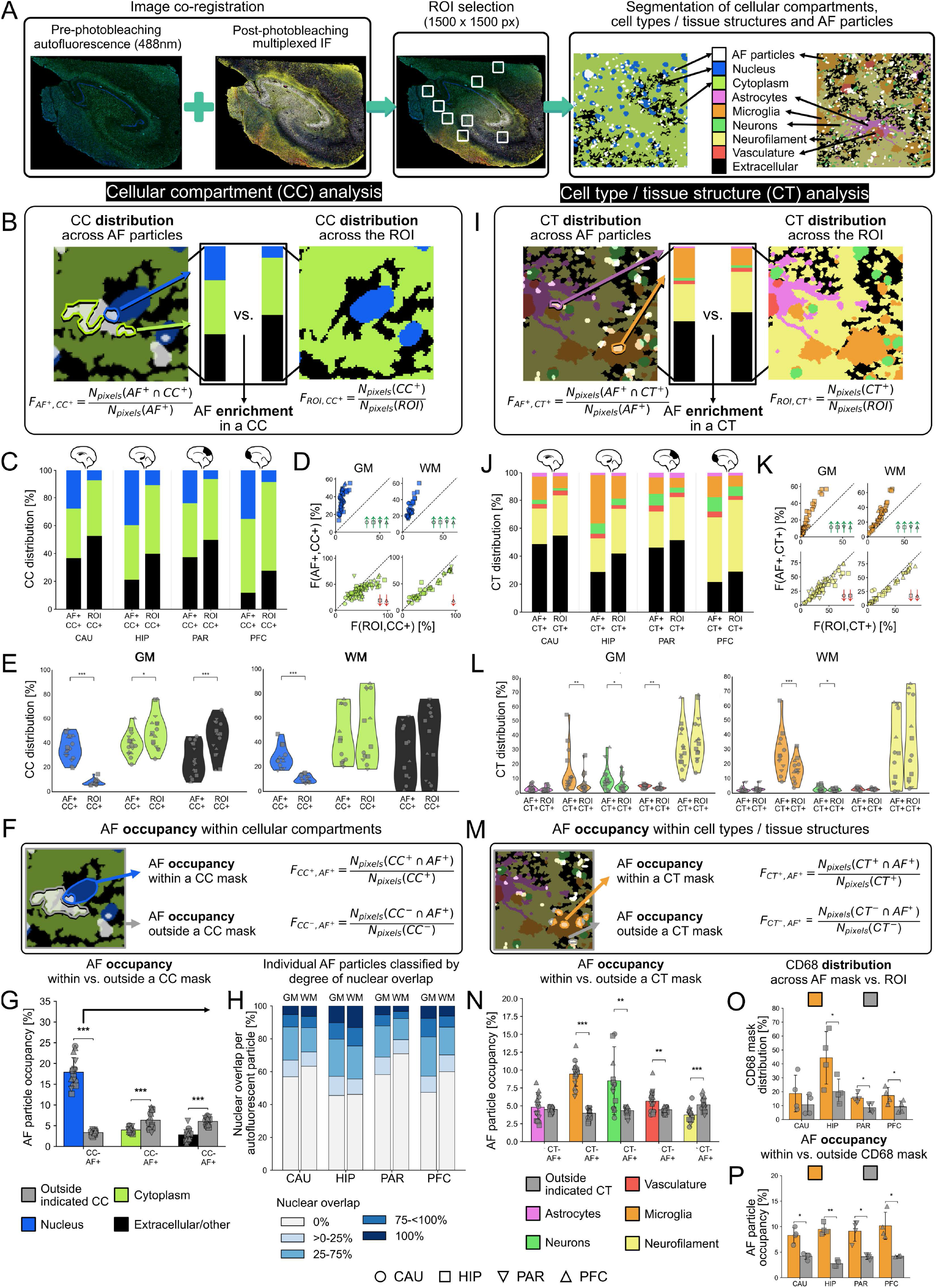
Spatial organization of autofluorescent particles across cellular compartments (CC) and cell types/tissue structures (CT). Two complementary metrics were used: CC / CT distribution, defined as the fraction of autofluorescent particle masks localized within a given CC or CT, and autofluorescent particle occupancy, defined as the fraction of each CC or CT occupied by autofluorescent particles. **A.** Image-processing workflow for spatial characterization of autofluorescent particles. Pre-photobleaching autofluorescence images acquired in the 488 nm channel were co-registered with multiplexed immunofluorescence images following optimized photobleaching. Four gray matter and up to four white matter regions of interest (ROIs; 1500 × 1500 pixels) were selected from each sample. Binary masks were generated for autofluorescent particles, three cellular compartments and five cell types/tissue structures. **B–E.** Distribution of cellular compartments across autofluorescent particles and ROIs. **B.** Schematic illustrating CC distribution and distribution-enrichment analyses. For each CC, the fraction of the autofluorescent particle mask localized within that CC (F_AF+,CC+_) was compared with the fraction of the ROI occupied by that CC (F_ROI,CC+_). **C–E.** Results are shown as stacked bar charts **(C)**, scatter plots used for log₂ enrichment analysis **(D)**, and violin plots **(E)**. Nucleus is shown in blue, cytoplasm in green, and extracellular space in black. Stacked bar charts summarize fractions by brain region **(C)**, whereas scatter plots **(D)** and violin plots **(E)** show gray and white matter separately. Each point represents individual ROI **(D)** or mean sample **(E)** measurements, with a symbol indicating brain region. In (**D)**, green upward arrows indicate significant enrichment, while red downward arrows indicate significant depletion, based on two-sided linear mixed-effect models (*p* ≤ 0.05). In **(E)**, stars indicate statistical significance from two-sided linear mixed-effects models: * for *p* ≤ 0.05, ** for *p* ≤ 0.005, and *** for *p* ≤ 0.0005. **F-G.** Occupancy of autofluorescent particles within cellular compartments. **F.** Schematic illustrating occupancy analysis. For each CC, the fraction of the CC occupied by the autofluorescent particle mask (F_CC+,AF+_) was compared with autofluorescent occupancy outside the same CC (F_CC-,AF+_). **G.** Results are shown as bar charts comparing autofluorescence occupancy within and outside each CC. Each point represents one sample, with symbol indicating brain region. Stars indicate statistical significance from two-sided linear mixed-effects models: * for *p* ≤ 0.05, ** for *p* ≤ 0.005, and *** for *p* ≤ 0.0005. **H.** Nuclear overlap analysis of individual autofluorescent particles. Autofluorescent particles were grouped according to the percentage of their area overlapping the nuclear mask. The stacked bar chart shows the proportion of autofluorescent particles within each nuclear-overlap category (0%, >0–25%, 25–75%, 75–<100%, and 100%), summarized by brain region and tissue compartment. **I–L.** Distribution of cell types/tissue structures across autofluorescent particles and ROIs. **I.** Schematic illustrating distribution and distribution-enrichment analyses. For each CT, the fraction of the autofluorescent particle mask localized within that CT (F_AF+,CT+_) was compared with the fraction of the ROI occupied by that CT (F_ROI,CT+_). **J-L.** Results are shown as stacked bar charts **(J)**, scatter plots used for log₂ enrichment analysis **(K)**, and violin plots **(L)**. Astrocytes are shown in magenta, microglia in orange, neurons in green, neurofilament-positive structures (NEFL) in yellow, vasculature in red and extracellular space in black. Stacked bar charts **(J)** summarize results by brain region, whereas scatter plots **(K)** and violin plots **(L)** show gray and white matter separately. Each point represents individual ROI **(K)** or mean sample **(L)** measurements, with a symbol indicating brain region. In **(K)**, green upward arrows indicate significant enrichment, while red downward arrows indicate significant depletion, based on two-sided linear mixed-effect models (*p* ≤ 0.05). In **(L)**, stars indicate statistical significance from two-sided linear mixed-effects models: * for *p* ≤ 0.05, ** for *p* ≤ 0.005, and *** for *p* ≤ 0.0005. **M-N.** Occupancy of autofluorescent particles within cell types/tissue structures. **M.** Schematic illustrating occupancy analysis. For each CT, the fraction of the CT occupied by the autofluorescent particle mask (F_CT+,AF+_) was compared with autofluorescence occupancy outside the same CT (F_CT-,AF+_). **N.** Results are shown as bar charts comparing autofluorescence occupancy within and outside each CT. Each point represents one sample, with a symbol indicating brain region. Stars indicate statistical significance from two-sided linear mixed-effects models: * for *p* ≤ 0.05, ** for *p* ≤ 0.005, and *** for *p* ≤ 0.0005. **O–P.** Distribution and occupancy of autofluorescent particles within CD68-positive regions. **O.** Distribution analysis comparing the fraction of the autofluorescent particle mask localized within CD68-positive regions (F_AF+,CD68+_) with the fraction of CD68-positive regions in the ROI (F_ROI,CD68+_). **P.** Occupancy analysis comparing the fraction of the CD68 mask occupied by the autofluorescent particle mask (F_CD68+,AF+_) with autofluorescence occupancy outside the CD68 mask (F_CD68-,AF+_). Each point represents one sample, with a symbol indicating brain region. Stars indicate statistical significance from paired t test: * for *p* ≤ 0.05, ** for *p* ≤ 0.005, and *** for *p* ≤ 0.0005. **Sample size:** *n* (CAU) = 4, *n* (HIP) = 4, *n* (PAR) = 4, *n* (PFC) = 4.

Across brain regions, the largest fraction of the autofluorescent particle mask was localized within the cytoplasmic compartment, averaging 39.2% in gray matter and 44.3% in white matter, followed by the nuclear compartment (35.4% and 27.4%, respectively) **(Figure 8C–E; Supplementary Figure 20)**. Because cytoplasmic regions occupy a larger fraction of the ROIs than nuclei, distribution alone does not indicate preferential localization. Comparison with cellular compartment distribution across the entire ROIs revealed that autofluorescence was enriched in nuclear regions and underrepresented in cytoplasmic and extracellular regions **(Figure 8C–E, Supplementary Figures 20–21)**.

This enrichment in nuclear regions was independently supported by occupancy analysis. Autofluorescence occupancy was highest within nuclear masks, and the nuclear compartment was the only compartment with significantly greater occupancy inside than outside the corresponding mask. On average, autofluorescent particles occupied 20.5% of the nuclear mask in gray matter and 14.1% in white matter, compared with only 2.9% and 4.1%, respectively, outside the nuclear mask. Cytoplasmic and extracellular compartments showed the opposite pattern **(Figure 8G; Supplementary Figure 22)**. However, enrichment and increased occupancy within nuclear regions did not necessarily imply intranuclear localization. To distinguish intranuclear from perinuclear localization, we classified individual autofluorescent particles by their degree of nuclear overlap **(Figure 8H; Supplementary Figure 23)**. Only 7.0% of autofluorescent particles in gray matter and 7.3% in white matter were completely contained within nuclei, whereas 52.0% and 60.1%, respectively, showed no nuclear overlap. Thus, despite their enrichment within nuclear regions, most autofluorescent particles were not intranuclear, indicating predominantly perinuclear localization **(Figure 8H; Supplementary Figure 23)**.

We next extended these analyses to cell types and tissue structures. Neurofilament and microglial regions accounted for the largest fractions of the autofluorescent particle mask **(Figure 8J)**, with greater microglial representation in white matter **(Figure 8L; Supplementary Figure 24)**. On average, neurofilament masks accounted for 31.4% and 28.8% of the autofluorescent particle mask in gray and white matter, respectively, whereas microglial masks accounted for 14.3% and 27.5%. However, accounting for the abundance of these structures across the ROIs revealed a markedly different pattern: autofluorescence was consistently enriched within microglial regions but underrepresented within neurofilament-positive regions across all analyzed brain regions **(Figure 8K; Supplementary Figures 24–25)**. Thus, extensive neurofilament overlap reflected its abundance in the tissue, whereas microglial overlap reflected preferential autofluorescence localization.

Occupancy analysis independently supported this preferential microglial localization **(Figure 8N)**. Autofluorescent particles occupied on average 10.1% of microglial masks in gray matter and 8.8% in white matter, compared with 3.7% and 4.4%, respectively, outside these masks. Neuronal and vascular masks showed a similar pattern, astrocytic masks showed no clear difference, and neurofilament masks showed the opposite pattern, with slightly lower occupancy inside than outside the mask, particularly in gray matter **(Supplementary Figure 26)**.

Given this preferential microglial localization, we next examined whether autofluorescence was also associated with CD68-positive regions, marked by CD68, a marker of lysosomal activity and phagocytic myeloid state. Autofluorescence was consistently enriched within CD68-positive regions, most prominently in hippocampus, where 43% of the autofluorescent particle mask overlapped the CD68 mask despite CD68-positive regions comprising only approximately 19.3% of the analyzed hippocampal ROIs **(Figure 8O; Supplementary Figure 27)**. Autofluorescence occupancy was likewise greater within than outside CD68-positive regions across all brain regions and tissue compartments, averaging 9.2% within CD68 masks compared with 3.9% outside **(Figure 8P; Supplementary Figure 27)**.

Together, these analyses demonstrate that autofluorescence is non-randomly organized in aged fresh-frozen human brain, showing preferential localization in the vicinity of nuclei and within microglial and CD68-positive regions. This structured localization increases the potential for residual autofluorescence to confound marker interpretation and single-cell quantification, reinforcing the importance of effective autofluorescence suppression in high-plex spatial proteomics.

## DISCUSSION

Autofluorescence is a major obstacle for fluorescence microscopy of aged human tissues, yet methods optimized for fresh-frozen human brain remain limited. Here, we developed a photobleaching workflow for effective autofluorescence suppression and a 28-plex DNA-barcoded antibody panel for fresh-frozen human brain, which we then combined to characterize the spatial organization of brain autofluorescence.

Formalin-fixed, paraffin-embedded (FFPE) brain tissue remains the standard for histopathology because of its excellent morphological preservation ^15,16^ and widespread availability in archival biobanks ^15,17^. However, formaldehyde fixation and paraffin processing introduce protein cross-links and other chemical modifications that can compromise fixation-sensitive epitopes ^18^, as well as protein extraction and mass-spectrometry-based protein analysis ^15,17^. As spatial biology increasingly moves toward integrating complementary modalities ^19^, combining multiplexed immunofluorescence with other spatial techniques may dictate the use of fresh-frozen over FFPE tissue when these modalities prefer or even require fresh-frozen specimens, such as matrix-assisted laser desorption/ionization mass spectrometry imaging ^20^. This highlights the need for robust multiplexed immunofluorescence workflows compatible with fresh-frozen human brain, particularly for multimodal studies using the same or consecutive sections.

Brain autofluorescence is largely attributed to the accumulation of naturally occurring lipofuscin pigments ^21,22^. These highly heterogeneous aggregates of oxidized and cross-linked macromolecules consist primarily of lipids and misfolded or damaged proteins, with smaller amounts of carbohydrates and metal cations ^8,21,23–25^. Because no universally accepted molecular marker exists, lipofuscins are commonly identified by their histological appearance and characteristic autofluorescence ^8^. We therefore refer to the endogenous fluorescent structures detected here as autofluorescent particles rather than assigning a definitive molecular identity. Nevertheless, their broad spectral emission, morphology, age association, and widespread abundance strongly suggest that they predominantly represent lipofuscin or closely related autofluorescent material ^8,21,23–25^.

Consistent with previous reports ^21,24,25^, autofluorescent particles exhibited broad fluorescence across all four brain regions and three imaging channels (488, 550, and 647 nm), with the strongest signal intensity in the 488 nm channel **(Figure 1C)**. This has important implications for multiplexed antibody panel design, favoring assignment of weaker antibody signals to longer-wavelength channels with lower autofluorescence, where compatible with other panel-design constraints.

Beyond representing a technical challenge, autofluorescence emerged as a biologically structured feature of the brain tissue. Autofluorescent particles were larger and brighter in gray matter, but more numerous in white matter, resulting in greater overall autofluorescence burden in white matter **(Figure 1D–E)**. Particle masks consequently recapitulated brain architecture, generating virtual histology-like maps distinguishing gray from white matter **(Figure 1B)**, suggesting that autofluorescence should not be viewed solely as unwanted background, but also as a source of biological information for multimodal spatial analyses of the human brain. Although previous studies reported greater lipofuscin accumulation in gray matter ^8,24^, our findings highlight that such conclusions depend on how autofluorescence burden is defined: While autofluorescent particle size and intensity were greater in gray matter, density and tissue coverage were greater in white matter **(Figure 1D-E; Supplementary Figure 19)**. Future studies should therefore consider multiple metrics of autofluorescence burden rather than relying on a single measure.

The gray-white matter differences may partly reflect their distinct cellular composition and metabolism, with gray matter enriched in neuronal cell bodies, dendrites, synapses, and astrocytes and white matter dominated by myelinated axons and oligodendrocytes ^26^. Because lipofuscin formation is closely linked to cellular metabolism and lysosomal function ^21^, these compositional differences may contribute to compartment-specific accumulation of autofluorescent material. Neurodegenerative pathology may further influence these patterns by altering lysosomal processing and promoting the formation and/or accumulation of lipofuscin-related material ^8^. Studies in larger, disease-stratified cohorts will be needed to disentangle the relative contributions of tissue composition, aging, and neurodegenerative pathology to regional autofluorescence patterns.

Autofluorescent particle intensity increased with donor age across examined brain regions **(Figure 1F; Supplementary Figure 3A)**, consistent with progressive accumulation of degradation-resistant lipofuscin during aging ^21^. Particle size and density were not associated with age **(Figure 1H; Supplementary Figures 4A and 5A)**, likely reflecting the predominantly elderly cohort. However, while lipofuscin has been reported in young mice ^24,27^, evidence from pediatric human brain remains limited to isolated reports ^28^, highlighting the need for human lifespan studies to determine when autofluorescent particles emerge and how they evolve with age.

The abundance, intensity, and broad spectral range of autofluorescence pose a particular challenge for multiplexed immunofluorescence microscopy. Available approaches to address autofluorescence include chemical quenching, fluorescence lifetime imaging ^11,29^, spectral unmixing ^30^, and non-chemical ^24,25^ or chemical-assisted photobleaching ^14^. Chemical quenchers such as Sudan Black or TrueBlack can effectively suppress autofluorescence in conventional immunofluorescence microscopy settings, but may also reduce specific antibody signals ^24,25^. In our hands, TrueBlack had such an effect, suppressing strong autofluorescence but also reducing specific staining intensity. Moreover, their post-staining application limits compatibility with iterative workflows ^31^. Fluorescence lifetime imaging ^11,29^ and spectral unmixing ^30^ can distinguish autofluorescence from fluorophore-derived signals but require specialized instrumentation. Photobleaching therefore offered the most compatible approach, particularly because chemical-assisted photobleaching substantially shortens high-intensity LED exposure times while improving quenching efficiency compared with LED exposure alone ^32^. However, existing protocols were developed primarily for mouse ^24^ or FFPE human tissue ^14,25,27^, rather than fresh-frozen human brain.

We therefore developed a chemical-assisted photobleaching workflow ^7,14^. However, direct transfer of FFPE photobleaching protocols to aged fresh-frozen human brain caused marked region- and compartment-dependent tissue damage **(Figures 2)**, with gray matter and deep subcortical regions particularly vulnerable. This suggests that tissue composition and local microenvironment influence tissue resilience during photobleaching. Importantly, preserving tissue adhesion and histological architecture is not merely a technical concern, but essential for reliable cyclic multiplexed imaging and quantitative analysis.

A key methodological advance was the identification of EDTA supplementation as a simple and effective strategy to preserve fresh-frozen tissue during photobleaching **(Figure 2C-J)**. Hydrogen peroxide, the principal active component of chemical photobleaching protocols ^14^, promotes autofluorescence bleaching through oxidation of the endogenous fluorophores ^33^ but under illumination, it can also generate highly reactive oxygen species (ROS) in the presence of transition-metal ions such as iron ^34^. Given the relatively high iron content of certain brain regions including the basal ganglia where caudate-putamen is located ^13,35^, we hypothesized that metal-catalyzed ROS generation may contribute to the pronounced photobleaching-induced tissue damage. The protective effect of EDTA, a metal-ion chelator ^36^, is consistent with a role for metal-dependent chemistry, although the underlying mechanism requires further investigation.

Regardless of the underlying mechanism, the optimized EDTA-supplemented workflow substantially reduced autofluorescence while preserving tissue morphology and cellular content **(Figure 3)**. Autofluorescence suppression (58–70% across analyzed channels; **Figure 3B–C**) was comparable to that reported for FFPE protocols (45.9–75.5%)^25^, despite the greater fragility of fresh-frozen tissue. This simple and accessible strategy may therefore have broader applicability to fluorescence-based analyses of fresh-frozen tissues requiring autofluorescence suppression.

Following effective autofluorescence suppression, we established a 28-plex, neuroscience-oriented antibody panel for fresh-frozen human brain **(Figure 4; Supplementary Table 6)**. Antibody panel development remains a major bottleneck in multiplexed immunofluorescence because antibody performance varies substantially across tissue types and preservation methods, a challenge compounded on platforms requiring antibodies to be conjugated further to DNA barcodes ^31^. Reflecting this extensive optimization, 64 antibodies were evaluated, of which 37 underwent in-house DNA-barcode conjugation, ultimately yielding the final 28-plex panel **(Supplementary Table 5)**. Although multiplexed neuro-panels have recently been reported for mouse brain ^37^ and human FFPE brain ^7^, to our knowledge, this is the first DNA-barcoded multiplexed antibody panel specifically developed for fresh-frozen human brain. Together with the optimized photobleaching workflow, it provides a practical resource for high-dimensional spatial profiling of fresh-frozen human brain.

Co-registration of pre-photobleaching autofluorescence with post-photobleaching multiplexed images enabled direct spatial comparison of autofluorescent particles with protein marker signals in the same tissue **(Figure 7**, **Figure 8)**. Protein marker signals at autofluorescent particle locations varied predominantly by anatomical region rather than tissue compartment, donor age, or diagnosis **(Figure 7C; Supplementary Figure 16)**, with cortical particles coinciding with higher neuronal marker signals and hippocampal particles with higher glial marker signals **(Supplementary Table 8)**. This suggested that anatomical context shapes the local protein environment surrounding these particles. However, because these patterns could also reflect regional differences in cellular and protein composition, we next used binary masks to examine autofluorescent particle localization more directly across cellular compartments, cell types, and tissue structures **(Figure 8)**.

Cellular compartment analysis revealed enrichment of autofluorescent particles in nuclear regions **(Figure 8C, E, G; Supplementary Figures 20–22)**, initially appearing inconsistent with the described localization of lipofuscin within lysosomal or lysosome-related structures in the cytoplasm ^8,24^. However, most particles showed no or only partial nuclear overlap, indicating a predominantly perinuclear rather than intranuclear localization **(Figure 8H; Supplementary Figure 23)**. Although higher-resolution three-dimensional imaging is needed to define their precise intracellular localization, these findings consistently place autofluorescent particles in close proximity to the nucleus, supporting the nuclear vicinity as a characteristic feature of their spatial organization.

Among cell types and tissue structures, autofluorescence showed the strongest enrichment within microglial and CD68-positive regions **(Figure 8K-P; Supplementary Figures 24–27)**, consistent with preferential lipofuscin accumulation in microglia reported both in mouse ^24^ and human brain ^8^. Studies in mice ^24,38^, and non-human primates ^38^ further suggest that microglia are the earliest brain cell type to accumulate lipofuscin-associated autofluorescence and that autofluorescent microglia even exhibit distinct ultrastructural and proteomic characteristics ^24,38^. Our independent observations of enriched autofluorescence in CD68-positive regions across brain regions and compartments **(Figure 8O, P; Supplementary Figure 27)**, and strong pixel-wise spatial correlations between CD68 signal and autofluorescence **(Supplementary Table 9)** support such preferential accumulation within lysosomal compartments, potentially resulting from incomplete degradation or impaired clearance ^8,39^.

These findings have broader implications for understanding aging and neurodegeneration. Lipofuscin accumulation has been reported to be particularly prominent in regions vulnerable to Alzheimer’s disease, including the hippocampus, and has been spatially associated with both amyloid-β and tau pathology ^8^. It has also been proposed to impair lysosomal function and thereby promote protein aggregation and ROS production ^8^, although whether it contributes directly to neurodegeneration or accumulates secondary to lysosomal dysfunction remains unclear ^8^. Applying the spatial framework developed here to larger disease-stratified cohorts could help resolve relationships among lipofuscin accumulation, lysosomal dysfunction, pathological protein deposition, and neurodegeneration.

These findings also have methodological implications for immunofluorescence imaging of the aged human brain. Autofluorescence can generate false-positive signals, impair cell segmentation, and distort antibody staining patterns, particularly in studies of microglia, where lipofuscin-associated autofluorescence may be mistaken for engulfed material and confound conclusions regarding microglial state ^24^. Some antibodies against amyloid-β have also been reported to recognize neuronal lipofuscin, either because of shared epitopes or the close spatial association of both within lysosomal compartments ^8^. Effective autofluorescence suppression, appropriate negative controls, and careful antibody validation are therefore essential for reliable imaging of the aged brain.

Overall, this work establishes a practical experimental and analytical framework for multiplexed spatial proteomics of fresh-frozen human brain while addressing autofluorescence as a major technical barrier. Beyond improving imaging reliability, integration of pre-photobleaching autofluorescence with multiplexed protein maps provides an approach for studying autofluorescence itself as a spatially organized tissue feature. This framework may enable future studies to both control and exploit autofluorescence in spatial analyses of brain aging and neurodegeneration.

## METHODS

### Ethics statement, human *post-mortem* brain cohort and tissue preparation

The use of human *post-mortem* brain tissue was approved by the Stockholm Regional Ethics Review Board (Dnr 2014/1366-31). All tissue samples were anonymized before analysis.

Fresh frozen post-mortem human brain tissue was obtained from the King’s College London Brain Bank. The study included 34 tissue samples from 21 unique donors, comprising 10 female and 11 male donors, with a mean age of 79.7 ± 8.7 years and a mean post-mortem delay (PMD) of 33.6 ± 8.3 h **(Supplementary Tables 1 and 2)**. Donors included individuals diagnosed with Parkinson’s disease (PD), Parkinson’s disease dementia (PDD) or dementia with Lewy bodies (DLB), as well as age-matched cognitively healthy controls (CON) **(Supplementary Table 1)**. Neuropathological assessment was performed at King’s College London. Four brain regions were analyzed: caudate–putamen (CAU; *n* = 14), hippocampus (HIP; *n* = 4), parietal cortex (PAR; *n* = 12) and prefrontal cortex (PFC; *n* = 4). In total, 34 tissue samples from 21 unique donors were included.

Tissue was cryosectioned at 10 µm thickness using a CM1860 UV cryostat (Leica Microsystems) maintained at −15 °C. Sections were thaw-mounted onto SuperFrost Plus Gold Adhesion microscope slides (K5800AMNZ72, Epredia) and stored at −80 °C until use.

### Optimization and evaluation of autofluorescence photobleaching in human fresh frozen brain tissue

To optimize photobleaching conditions, tissue autofluorescence was measured before and after photobleaching treatment. Fresh-frozen tissue sections were removed from storage at −80 °C and allowed to equilibrate to room temperature (RT) for 15–20 min. Sections were fixed in 4% paraformaldehyde (PFA; 043368.9M, Thermo Fisher Scientific) for 15 min at RT and subsequently washed three times in phosphate-buffered saline (PBS; 18912.014, Gibco) for 15 min per wash. Sections were mounted using a DAPI-containing mounting medium (P36931, Invitrogen) and coverslips (631-0147, VWR). Imaging was performed using a PhenoCycler-Fusion 2.0 system (Akoya Biosciences, now Quanterix) equipped with a 10× objective, corresponding to a total magnification of 20×, and 0.5 µm/pixel resolution. Autofluorescence was recorded in the 488, 550, and 647 nm channels with a fixed exposure time of 300 ms per channel. DAPI images were acquired using an exposure time of 5 ms.

Following baseline imaging, coverslips were removed by immersing the slides in PBS until they detached. Tissue sections were then washed three times in PBS for 5 min before photobleaching.

Photobleaching was performed by exposing the sections to high-intensity LED illumination (32,000 lux; SunnyLight, OneSunrise) during two consecutive 45 min incubations in a photobleaching solution. The starting solution was adapted from a previously published protocol for formalin-fixed, paraffin-embedded tissue (PMID: 31534232) and contained 4.5% H₂O₂ (31642, Sigma-Aldrich) and 26.4 mM NaOH (1.09137.1000, Merck) in PBS. During optimization, the addition of Tris–EDTA (AB93684, Abcam) at final concentrations of 0.025% and 0.05% (v/v) was evaluated. In the optimized protocol, 0.05% (v/v) Tris-EDTA was added to the first photobleaching solution, together with 4.5% H₂O₂ and 26.4 mM NaOH. The second 45 min incubation step was performed in PBS solution containing only 4.5% H₂O₂ and 26.4 mM NaOH.

Following treatment, sections were washed, remounted, and reimaged using the same acquisition settings as before photobleaching. All fluorescence images were acquired and stored as 16-bit multichannel QPTIFF files.

### Design of the multiplexed antibody panel for spatial protein mapping in fresh-frozen human brain tissue

To develop the oligonucleotide-conjugated antibody panel, a selection of biologically relevant primary antibodies was first evaluated using conventional immunofluorescence staining. Staining specificity was assessed by comparison of cellular or anatomical signal distribution with publicly available reference data. Antibodies showing specific and reproducible staining patterns were subsequently conjugated to oligonucleotide barcodes and re-evaluated after conjugation. Only antibodies that retained specific and reliable staining were included in the final multiplexed imaging panel. All antibodies assessed during panel development are listed in **Supplementary Table 5.**

#### Immunofluorescent staining protocol

Conventional immunofluorescence staining was performed on fresh-frozen human brain tissue sections to evaluate antibody specificity and staining patterns. The protocol was modified to include an optimized photobleaching step to reduce tissue autofluorescence.

Briefly, tissue sections were removed from storage at −80 °C and allowed to equilibrate to RT for 15–20 min. Sections were fixed in 4% PFA for 15 min at RT and washed in PBS. The optimized photobleaching protocol was then applied, followed by PBS washes. A hydrophobic barrier was carefully drawn around each tissue section using a PAP pen (ab2601, Abcam). Primary antibodies were diluted in PBS containing 0.3% Triton X-100 (T8787-100ML, Sigma-Aldrich). Sections were incubated with primary antibodies for 3 h at RT or overnight at 4 °C. All tested antibodies and their corresponding working dilutions are listed in **Supplementary Table 5**. Following primary antibody incubation, sections were washed in Tris-buffered saline containing 0.1% Tween 20 (TBS-T; TBS, 09-7500-100, Medicago; Tween 20, P9416-100ML, Sigma-Aldrich) and non-specific binding was blocked by incubating the sections for 30 min at RT in TNB buffer, prepared by dissolving TSA Blocking Reagent (FP1020, Akoya Biosciences) in TBS. Secondary antibodies were diluted 1:800 in TNB buffer containing 0.02% Hoechst 33342 (H3570, Invitrogen) and incubated with the tissue sections for 90 min at RT. Sections were then washed in TBS-T and mounted using Fluoromount-G (00-4958-02, Invitrogen) and coverslips (631-0147, VWR). All incubation steps were performed in a humidified chamber to prevent tissue drying.

Whole-section images were acquired using a PhenoCycler-Fusion 2.0 system (Akoya Biosciences) equipped with a 10× objective, corresponding to a total magnification of 20×, and 0.5 µm/pixel resolution.

#### Antibody conjugation with oligonucleotide barcodes

Antibodies that demonstrated specific staining but were not commercially available in DNA-barcoded format were conjugated in-house using the Antibody Conjugation Kit (232195, Akoya Biosciences), as previously described ^40^. Briefly, conjugation was performed in 50 kDa MWCO filter tubes (UFC5050, Millipore) that were blocked for non-specific binding with Filter Blocking Solution (200033, Akoya Biosciences). After each step, the solution present in the filter was removed by centrifuging the filter tube at 12,000 × g for 8 min and discarding the flow-through. To break the disulfide bonds (needed for DNA barcode ligation), 50 µg of antibody of interest was added to the filter and incubated in the Antibody Reduction Master Mix, containing Reduction solution 1 (200032, Akoya Biosciences) and Reduction solution 2 (200028, Akoya Biosciences) in a 1:41.67 ratio, for 30 min at RT. Lyophilized DNA barcodes were resuspended in 10 µL of Ambion Nuclease free water (AM9937, Invitrogen) and 210 µL of Conjugation Solution (200029, Akoya Biosciences). Reduced antibodies were washed with Conjugation Solution and incubated with the selected barcode for 2 h at RT. The conjugated antibody was then purified by three successive washes with the Purification Solution (200030, Akoya Biosciences) and resuspended in the Antibody storage solution (200031, Akoya Biosciences).

Conjugated antibodies were used no earlier than 48 h after conjugation, as recommended by the manufacturer, to minimize non-specific nuclear staining by residual oligonucleotides. Successful conjugation was verified by comparing staining patterns obtained with conjugated antibodies in the multiplexed workflow to those generated using the corresponding unconjugated antibodies by conventional immunofluorescence (see previous section)

### Integration of optimized autofluorescence photobleaching with multiplexed spatial proteomics

High-plex immunostaining and imaging were performed using the PhenoCycler-Fusion Sample Kit (232193, Akoya Biosciences) according to the manufacturer’s protocol for fresh-frozen tissue (PhenoCycler-Fusion User Guide v1.0.7, PD-000011 Rev J), based on the method described by Black et al. (2021) ^31^.

Fresh-frozen tissue sections were retrieved from −80 °C storage and dried at RT for 5 min on Drierite beads (43056, Alfa Aesar). Sections were incubated in acetone (650501, Sigma-Aldrich) for 10 min at RT, air-dried for 2 min and rehydrated in Hydration Buffer (240196, Akoya Biosciences). Sections were then fixed in 1.6% PFA for 10 min at RT and washed again in Hydration Buffer.

Baseline autofluorescence images were acquired as described above. Sections were then subjected to the optimized photobleaching protocol consisting of two consecutive 45 min incubations under high-intensity LED illumination (32,000 lux; SunnyLight, OneSunrise) for. The first photobleaching solution contained 4.5% H₂O₂, 26.4 mM NaOH and 0.05% (v/v) Tris–EDTA in PBS, whereas the second solution contained 4.5% H₂O₂ and 26.4 mM NaOH in PBS. Following photobleaching, sections were remounted and reimaged using identical acquisition settings to assess the reduction in autofluorescence. Images acquired before and after photobleaching were stored as 16-bit multichannel QPTIFF files.

After imaging, coverslips were removed, and the multiplexed immunofluorescent staining protocol was resumed. Sections were equilibrated in Staining Buffer (240198, Akoya Biosciences) for 30 min at RT and incubated overnight at 4 °C with the oligonucleotide-conjugated antibody cocktail. Antibodies were diluted in Blocking Buffer comprising Staining Buffer with 2.4% (v/v) of each blocker; Blocker N (232108), Blocker G (232109), Blocker J (232110) and Blocker S (232111; all Akoya Biosciences). All antibody specifications and working dilutions are provided in **Supplementary Table 6**.

Following antibody incubation, sections were washed in Staining Buffer and fixed in 1.6% PFA for 10 min at RT. Sections were then washed in PBS, incubated in ice-cold methanol (322415, Sigma-Aldrich) for 5 min, washed again in PBS and incubated with Fixative Reagent (232112, Akoya Biosciences) for 20 min at RT. After a final PBS wash, sections were either stored in Storage Buffer (232107, Akoya Biosciences) at 4 °C for up to 48 h or immediately assembled into flow cells (240205, Akoya Biosciences) using the Flow Cell Assembly Device and loaded onto the PhenoCycler-Fusion instrument.

For each experiment, a 96-well reporter plate (7000006, Akoya Biosciences) was prepared, with each well corresponding to a single imaging cycle per slide. Each well contained up to three fluorescent reporters, each diluted 1:50 in 250 µl of Reporter Stock Solution, comprising PhenoCycler-Fusion Buffer, prepared from 10% (v/v) 10× Buffer (7000001, Akoya Biosciences) and 10% (v/v) Buffer Additive (240257, Akoya Biosciences), supplemented with 8.3% (v/v) Assay Reagent (7000002, Akoya Biosciences) and 3.3% (v/v) nuclear stain (H3570, Invitrogen).

The cyclic staining and imaging workflow was designed using PhenoCycler Experiment Designer v2.1.0 (Akoya Biosciences; **Supplementary Table 7**). The DAPI exposure time was fixed at 2 ms throughout all imaging cycles. Automated reporter hybridization, imaging and reporter removal were performed using Fusion v2.3.1 software (Akoya Biosciences). Images were acquired using a 10× objective, corresponding to a total magnification of 20× and 0.5 µm/pixel resolution. Image alignment and background subtraction, using the first and final blank cycles (containing nuclear stain only), were performed automatically within the instrument software. Raw images were stored as 16-bit QPTIFF files, and the final registered multichannel images were exported as 8-bit QPTIFF files.

### Hematoxylin and eosin (H&E) staining

Hematoxylin and eosin (H&E) staining was performed on fresh-frozen tissue sections following photobleaching optimization experiments or completion of the PhenoCycler-Fusion workflow. Before staining, coverslips were removed by spontaneous detachment in PBS, as described above, whereas PhenoCycler-Fusion flow cells were removed using a custom procedure designed to minimize tissue damage.

Following completion of PhenoCycler-Fusion imaging, slides were kept fully immersed in PBS at 4 °C. Before flow-cell removal, both flow-cell openings were sealed with Fixogum to prevent liquid exchange. Slides were wrapped in Kimwipes and incubated in acetone for 3 h, after which the flow cells were removed manually by applying gentle pressure. Residual adhesive was carefully removed from the edges of the slide using 100% ethanol, avoiding dehydration of the tissue sections. Slides were subsequently stored in PBS at 4 °C until H&E staining. For sections that had not undergone prior staining or imaging, slides were removed from storage at −80 °C and allowed to equilibrate to RT for 15–20 min. Sections were fixed in 4% PFA for 15 min at RT and washed three times in PBS for 15 min per wash.

Sections were incubated in hematoxylin solution (S3309, Dako) for 15 min and rinsed under running tap water for at least 2 min. Differentiation was performed by immersion in 0.1% (v/v) HCl (1.00316.1000, Merck) prepared in 70% ethanol for 3 s, followed by sequential rinsing in 70% and 90% ethanol. Sections were subsequently incubated in freshly acidified eosin solution (HT110216, Sigma-Aldrich) for 11 min. After eosin staining, slides were rinsed under running tap water and dehydrated by sequential immersion in 90% ethanol for 3 s and 100% ethanol for 3 s. Sections were cleared in HistoChoice clearing agent (H2779, Sigma-Aldrich) for 4 min and mounted in DPX mounting medium (6522, Sigma-Aldrich). Slides were allowed to dry overnight and were cleaned with 100% ethanol where required.

Whole-section bright-field images were acquired using the PhenoCycler-Fusion imaging system equipped with a 10x objective, yielding a total magnification of 20× and a resolution of 0.5 µm/pixel. Images were stored as 8-bit QPTIFF files.

Where hematoxylin staining appeared visually dominant, red color balance was adjusted in ImageJ (v 1.54g) ^41^ during image post-processing to improve image representation. Identical adjustment parameters were applied across all samples within each experiment. White-balance correction was applied to all H&E images using the White Balance Correction_1.0 plugin in ImageJ/Fiji (https://github.com/pmascalchi/ImageJ_Auto-white-balance-correction/tree/master). Minor cosmetic adjustments were performed in Affinity Designer (v3.2.1).

### Processing of high-parametric imaging data

QPTIFF files generated from autofluorescence imaging before and after photobleaching, together with multiplexed immunofluorescence images, were processed using QuPath and custom Python workflows.

#### Image co-registration

To compare autofluorescence signals before and after photobleaching and to assess colocalization between autofluorescence and protein markers, images were aligned using the Warpy interactive alignment extension for QuPath (Warpy v0.4.2; QuPath v0.6.0) ^42^.

#### Anatomical tissue annotations

Following image alignment, tissue regions for analysis were manually annotated in QuPath using the Brush and Magic Wand tools. Imaging and tissue artefacts, including air bubbles, tissue folds and fabric fibers, were excluded from subsequent analyses. All annotations were exported as GeoJSON files. Furthermore, gray and white matter regions were annotated separately using synaptophysin to delineate gray matter. In caudate–putamen sections, the caudate and putamen gray-matter regions could not be reliably distinguished and were therefore analyzed collectively as CAU. These sections also contained white matter corresponding anatomically to the internal capsule. For consistency with the nomenclature used for other brain regions, these measurements are reported as CAU-WM, although they anatomically represent internal-capsule white matter.

#### Cell and autofluorescent particle segmentation

Separate segmentation workflows were performed to characterize all cells and all autofluorescent particles within the analyzed tissue regions.

Cells inside the annotated regions were segmented on aligned, unprocessed images using the Cell Detection tool in QuPath (v0.6.0), with the DAPI channel used for detection. The following parameters were applied: background radius = 10 µm, sigma = 2 µm, minimum area = 4 µm², maximum area = 400 µm², threshold = 20, and cell expansion = 2 µm.

Autofluorescent particles were segmented on the same aligned, unprocessed images using 488 nm channel as the autofluorescence detection channel. The following parameters were applied: background radius = 8 µm, sigma = 1.5 µm, minimum area = 0.05 µm², maximum area = 1000 µm², threshold = 300, and cell expansion = 2 µm.

Measurements describing cell or autofluorescent particle size, morphology and fluorescence intensity were exported separately for each sample as CSV files. Intensity measurements included the mean, standard deviation, minimum and maximum signal across all protein-marker and autofluorescence channels. Spatial x–y coordinates were also retained. Subsequent analyses were performed with custom Python scripts, using pandas and NumPy for data handling, scikit-image for image processing, SciPy and scikit-learn for analysis and Matplotlib, Seaborn and Plotly for visualizations.

Cell-level measurements were used for a preliminary assessment of cell type composition across the analyzed brain regions. Autofluorescent particle measurements were used to: (1) characterize particle size, density and autofluorescence intensity across brain regions; (2) assess associations between autofluorescence features and donor characteristics; (3) compare autofluorescence intensity before and after photobleaching; and (4) quantify mean protein marker intensities within autofluorescent particle masks.

#### Processing of segmented cell and autofluorescent particle data

CSV files containing cell or autofluorescent particle measurements were filtered to retain variables relevant for downstream analysis. Data from individual samples were subsequently merged into a single dataset.

Mean cell-level marker intensities were log^10^-transformed using a pseudo count of 1 × 10⁻⁹ to avoid undefined values at zero and subsequently z-score normalized independently for each marker across all cells.

Raw autofluorescent particle measurements were used to assess Spearman correlations between donor characteristics (age and post-mortem delay; PMD), and autofluorescent particle size, density, and fluorescence intensity, as well as to compare mean autofluorescent particle intensity before and after photobleaching. For comparisons across brain regions, autofluorescence intensities were log_2_-transformed.

For joint analyses of autofluorescence and protein-marker intensities within the autofluorescent particles, measurements were first rescaled to a common 0–1 range to account for differing image bit depths. Autofluorescence intensities derived from 16-bit images were divided by 65,535, whereas protein-marker intensities derived from 8-bit images were divided by 255. Rescaled intensities were log_10_-transformed using a pseudo count of 1 × 10⁻⁹ to avoid undefined values at zero, and subsequently z-score-normalized independently for each marker.

### Computational workflow for the analysis of cellular composition using multiplexed immunofluorescence data

Log_10_-transformed, z-score-normalized protein marker intensities derived from cell segmentation masks, together with cell centroid x–y coordinates, were organized in an AnnData object for downstream characterization of cellular composition across the analyzed brain regions.

Each cell was assigned as positive or negative for each protein marker independently, using two-component Gaussian mixture models implemented with the GaussianMixture class in scikit-learn. The initial threshold was defined as the intensity at which the weighted probability density functions of the two Gaussian components intersected. This intersection was determined numerically using the fsolve function in SciPy. To obtain a more stringent definition of marker positivity, positive threshold values were multiplied by 2.0, whereas negative threshold values were divided by 2.0, thereby shifting the adjusted threshold towards higher marker intensities. Several threshold multipliers were empirically evaluated beforehand, and a multiplier of 2.0 was selected based on optimal separation of marker-positive and marker-negative cell populations and subsequent cell type assignment. Cells with intensities above the adjusted threshold were classified as marker-positive, whereas cells with intensities equal to or below the threshold were classified as marker-negative. Markers for which threshold estimation was unsuccessful were assigned no marker-positive cells. For quality control, marker-intensity distributions were visualized alongside the fitted low- and high-intensity Gaussian components and their corresponding thresholds.

Cells were then assigned to broad cellular classes using predefined marker-combination rules. Neurons were defined as NeuN-positive cells, oligodendrocytes as OLIG2-positive cells, astrocytes as cells positive for both SOX9 and GFAP, microglia as cells positive for CD45, CD11c and TMEM119, other immune cells as CD45-positive cells, and vascular cells as CLDN5-positive cells. Cells meeting criteria for multiple classes were assigned according to the marker combination showing the highest normalized marker intensity. Cells that did not express any marker above the positivity threshold or did not satisfy any of the predefined marker combinations were classified as “Other”. Cell type assignments were validated by visual inspection of spatial cell maps **(Figure 6E)** and marker-intensity heatmaps **(Supplementary Figure 13)**.

### Computational workflow for spatial characterization of autofluorescence

To characterize the spatial organization of autofluorescent particles in the human brain tissue, three complementary analyses were performed: (1) analysis of mean protein marker intensities within autofluorescent particle masks **(Figure 7A, C)**, (2) pixel-wise spatial correlation analysis between autofluorescence signal in the 488 nm channel and protein marker signals **(Figure 7A, D, E)**, and (3) analysis of the spatial association between autofluorescent particle masks and cellular compartment or cell type masks **(Figure 8)**.

#### Analysis of mean protein marker intensities within autofluorescent particle masks

Mean protein marker intensities were extracted from autofluorescent particle segmentation masks across gray and white matter regions **(Figure 7A)**. Log_10_-transformed and z-score-normalized mean marker intensities were visualized as a heatmap. Unsupervised hierarchical clustering of both markers and samples was performed using correlation distance and average linkage, as implemented in Seaborn. Principal component analysis (PCA) was performed on the standardized marker intensity matrix. PCA scores were visualized according to brain region, diagnosis, and age, and marker loadings were used to identify the protein markers contributing most strongly to each principal component **(Supplementary Figure 16)**.

#### Pixel-wise spatial correlation analysis

To further characterize the spatial co-distribution of autofluorescence and protein marker expression, pixel-wise correlation analysis was performed on up to eight representative regions of interest (ROIs) (1500 × 1500 pixels) per sample, including four from gray matter and up to four from white matter **(Figure 7A)**. Samples containing insufficient white matter were analyzed using fewer white matter ROIs. The number of ROIs analyzed for each sample is summarized in **Supplementary Table 4**.

For each ROI, individual TIFF channels were loaded and converted to floating-point arrays. Intensities were rescaled according to image bit depth by dividing 16-bit images by 65,535 and 8-bit images by 255. Intensities for each channel were then independently z-score normalized. Only spatially corresponding pixels with finite values in both the 488 nm autofluorescence reference image and the respective protein marker image were included in analysis.

Pixel-wise correlation between the 488 nm autofluorescence signal and each protein marker was quantified using Spearman’s rank correlation coefficient. Correlation coefficients were calculated for each ROI and subsequently averaged at the sample and brain-region levels. Chord diagrams were used to visualize (i) the strongest pixel-wise spatial correlations between autofluorescence and protein markers **(Figure 7D)** and (ii) the protein markers exhibiting the greatest regional variation in correlation across the four analyzed brain regions, separately for gray and white matter **(Figure 7E)**.

#### Analysis of spatial organization of autofluorescent particles relative to cellular compartments and cell types/tissue structures

To determine the spatial organization of autofluorescent particles relative to cellular compartments and major brain cell types/tissue structures, binary masks were generated for autofluorescent particles, cellular compartments, and major brain cell types within selected ROIs **(Figure 8A)**.

Image processing was performed in Python using scikit-image ^43^. Briefly, single-channel images of relevant markers, including CD31, CD45, CLDN5, GFAP, HLA-DR, MAP2, NEFL, NeuN, DAPI and autofluorescence in the 488 nm channel, were extracted from selected ROIs **(Supplementary Table 4)**. Images were converted to floating-point format and contrast-normalized to a 0–1 intensity range using the 1^st^ and 99.9^th^ percentiles of non-zero pixel intensities. Binary marker masks were generated by automated, ROI-specific intensity thresholding. Thresholding methods were selected empirically based on visual agreement with manually inspected marker distributions. Otsu’s method was used for NEFL, whereas Yen’s method was used for all other markers and autofluorescent particles, as implemented in the threshold_otsu and threshold_yen functions of the skimage.filters module. Thresholds were calculated using non-zero normalized pixel values. Binary masks were post-processed by removing objects smaller than 5 pixels, followed by one round of binary dilation using a disk-shaped structuring element with a radius of 1 pixel, implemented with functions from the skimage.morphology module. For each marker and ROI, both raw and processed binary masks were saved as TIFF files.

Processed individual marker masks were combined to generate biologically defined cell type/tissue structure masks using predefined marker combinations Individual marker masks occupying more than 70% of the ROI were excluded from the combination step, except for NEFL masks, for which this filter was not applied. Neuronal masks were generated as the union of NeuN and MAP2 binary masks. Vascular masks were generated from CD31, CLDN5, and Collagen IV masks. Microglial masks were generated from CD68, CD45 and HLA-DR masks. Astrocyte masks were generated from GFAP (antibody 2) mask, and neurofilament masks were generated from NEFL masks. To generate mutually exclusive cell-type masks, a hierarchical subtraction workflow was applied. Vascular masks were generated after subtraction of the neuronal mask. Microglial masks were generated after subtraction of neuronal and vascular masks. Astrocyte masks were generated after subtraction of neuronal, vascular and microglial masks. Neurofilament masks were generated after subtraction of neuronal, vascular, microglial and astrocyte masks. This hierarchical assignment ensured that each pixel was assigned to only one cell type category. Final cell type masks were saved separately for each ROI (**Supplementary Figure 18**).

Cellular compartment masks were generated separately to distinguish nuclear, cytoplasmic and extracellular regions. DAPI-positive pixels were classified as the nuclear compartment. Cytoplasmic pixels were defined as pixels assigned to the combined cellular masks (neuronal, vascular, microglia, astrocyte and NEFL) after removal of DAPI-positive pixels. Pixels not assigned to either the nuclear or cytoplasmic compartments were classified as extracellular or other. This procedure generated mutually exclusive nuclear, cytoplasmic and extracellular masks for each ROI.

Two complementary spatial metrics were calculated. “Autofluorescence distribution” quantified the fraction of the 488 nm autofluorescent particle mask overlapping each cellular compartment or cell type mask **(Figure 8 B, I)**. Then, to account for differences in the overall abundance of each cellular compartment or cell type mask within the ROI, the observed fraction of autofluorescent particle-positive pixels assigned to a given cellular compartment or cell type was compared with the fraction of the cellular compartment or cell type in that ROI.

Autofluorescence distribution for each cellular compartment (CC) was defined as:

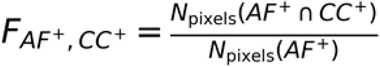

And autofluorescence distribution for each cell type/tissue structure (CT) was defined as:

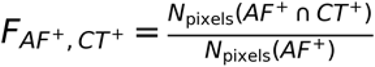

Abundance of each cellular compartment or cell type/tissue structure within the ROIs was defined as:

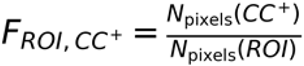

or:

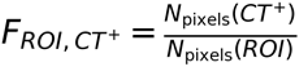

Where **F** denotes fraction, **AF** autofluorescence, **CT** cell type, **CC** cellular compartment, **ROI** region of interest, and **N_pixels_** the number of pixels.

The complementary metric, “autofluorescence occupancy”, quantified the fraction of each cellular compartment or cell type/tissue structure mask occupied by autofluorescent particles **(Figure 8 F, M)**. Then, autofluorescence occupancy within each cellular compartment or cell type/tissue structure was compared with autofluorescence occupancy outside.

Autofluorescence occupancy in each cellular compartment (CC) was defined as:

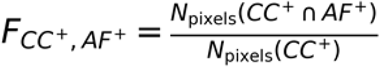

And autofluorescence occupancy in each cell type/tissue structure (CT) was defined as:

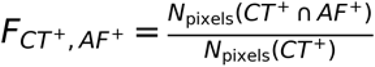

Autofluorescence occupancy outside each cellular compartment or cell type/tissue structure was defined as:

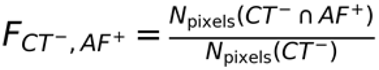

Or

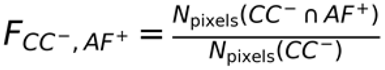

Where **F** denotes fraction, **AF** autofluorescence, **CT** cell type, **CC** cellular compartment, **ROI** region of interest and **N_pixels_** the number of pixels.

Autofluorescence distribution and occupancy analysis were also performed for CD68 masks.

For autofluorescent particle-level analysis, connected components in the binary autofluorescent particle mask were labelled as individual autofluorescent particles using eight-pixel connectivity. For each particle, the number and percentage of pixels overlapping the nuclei mask were calculated. Autofluorescent particles were then assigned to one of five nuclear-overlap categories: 0%, >0–25%, 25–75%, 75–<100% and 100%.

### Statistics

Statistical analyses were performed in Python (version 3.14.2) using SciPy (version 1.16.3) and statsmodels (version 0.14.6).

For analyses involving repeated measurements from the same donor, either across multiple brain regions **(Figures 1C–H and 8E, G, L, N)** or across multiple regions of interest (ROIs; **Figure 8D, K**), linear mixed-effects models were used. Donor was included as a random intercept, while variables relevant to each analysis were included as fixed effects. Outcome variables and fixed effects for each model are detailed in **Supplementary Table 10**. Models were fitted using restricted maximum likelihood (REML). The L-BFGS, Powell, and conjugate-gradient optimizers were attempted sequentially, and the first model that converged and produced valid, finite parameter estimates was retained. Models were fitted only when the prespecified minimum numbers of donors and observations were available **(Supplementary Table 10)**. These criteria were met for all analyses except the extracellular-compartment analysis of prefrontal-cortex white matter **(Supplementary Figures 21 and 25)**. Statistical significance was assessed using raw, two-sided mixed-model *p*-values, with *p* ≤ 0.05 considered statistically significant.

For comparisons with one paired measurement per donor under each experimental condition **(Figures 2G–J, 3C, and 8O–P)**, paired *t*-tests were used. Tests were two-sided unless a directional hypothesis had been specified a priori, in which case a one-sided paired *t*-test was applied **(Figures 2H–J and 3C)**. Statistical significance was defined as *p* ≤ 0.05.

Throughout the figures, statistical significance is indicated as follows: \**p* ≤ 0.05, \*\**p* ≤ 0.005, and \*\*\**p* ≤ 0.0005.

## DATA AVAILABILITY

Raw imaging datasets from (i) assay-optimization experiments (∼400 GB) and (ii) the final experiments, characterizing of autofluorescence in the caudate-putamen (CAU), hippocampus (HIP), parietal cortex (PAR), and prefrontal cortex (PFC) (∼235 GB), are available from the corresponding author upon request due to the large file sizes. CSV files generated following autofluorescent particle and cell segmentation, containing mean fluorescence-intensity measurements for autofluorescence and all analyzed protein markers, together with the binarized autofluorescence, cellular-compartment, and cell-type masks used to analyze the spatial organization of autofluorescence, will be deposited in Zenodo upon publication.

## CODE AVAILABILITY

The code used for data analysis, figure generation, and statistical analyses in this study is available in the following GitHub repository: https://github.com/Ayoglu-Lab.

## AUTHOR CONTRIBUTIONS

BA and IS conceived the study; IS and BA developed the methodology for the study; PS acquired the human tissue; IS and NB selected the human tissue specimen; IS and NB prepared the human tissue sections; IS and LG carried out the immunofluorescent experimental work; IS analyzed the data; IS, LG, and BA created the figures; BA and IS interpreted the data; BA and IS wrote the original manuscript; BA supervised and administered the project; BA, PS, and PEA acquired funding. All authors read and commented on the manuscript.

