## Supplementary Information for "Spatial organization and mitigation of autofluorescence in multiplexed spatial proteomics of aged fresh-frozen human brain"

### SUPPLEMENTARY TABLES

**Supplementary Table 1. Human brain tissue samples analyzed in the study.** Associated donor information including brain region, diagnosis, sex, age, and postmortem delay (PMD). (CAU - caudate-putamen, HIP - hippocampus, PAR - parietal cortex, PFC - prefrontal cortex), (CON - age-matched control, PD - Parkinson's disease, PDD - Parkinson's disease with dementia, DLB – Dementia with Lewy bodies)

| Sample number | Sample ID | Analyzed brain region(s) | Diagnosis | Sex | Age | PMD [h] |
|---|---|---|---|---|---|---|
| sample1 | A540/18 | CAU, HIP, PAR, PFC | CON | F | 80 | 57 |
| sample2 | A323/94 | CAU, HIP, PAR, PFC | PD | M | 74 | 10 |
| sample3 | NA170/03 | CAU, HIP, PAR, PFC | PDD | F | 84 | 27 |
| sample4 | A225/08 | CAU, HIP, PAR, PFC | DLB | M | 88 | 34.5 |
| sample5 | NA168/03 | CAU | PDD | F | 75 | 24 |
| sample6 | NA097/04 | CAU | PDD | M | 88 | 24 |
| sample7 | A188/03 | CAU | PD | F | 62 | 40.5 |
| sample8 | NA086/02 | CAU | PDD | F | 85 | n.a. |
| sample9 | NA085/02 | CAU | PDD | M | 75 | 36 |
| sample10 | A239/95 | CAU | CON | F | 79 | 38 |
| sample11 | ST18/02 | PAR | PDD | M | 79 | 30 |
| sample12 | A309/14 | PAR | DLB | M | 73 | 59 |
| sample13 | A065/11 | PAR | CON | M | 97 | 44 |
| sample14 | A121/21 | PAR | DLB | M | 84 | 82.5 |
| sample15 | ST01/01 | PAR | PDD | F | 83 | 24 |
| sample16 | A237/14 | PAR | CON | F | 83 | 39 |
| sample17 | A350/96 | PAR, CAU | PD | F | 85 | 9 |
| sample18 | A391/94 | PAR | PD | M | 59 | 23 |
| sample19 | A242/15 | CAU | CON | M | 82 | 26 |

| | | | | | | |
|---|---|---|---|---|---|---|
| sample20 | ST16/02 (NA094/02) | CAU | PDD | M | 80 | 26 |
| sample21 | A106/91 | CAU | DLB | F | 78 | 10 |

**Supplementary Table 2. Overview of the sample cohort analyzed across the four brain regions**, including number of donors, and donor age and post-mortem delay (PMD) ranges.

| Brain region | Number of donors | Age range [years] | PMD range [hours] |
|---|---|---|---|
| **Caudate-putamen** | 14 | 62-88 | 9-57 |
| **Hippocampus** | 4 | 74-88 | 10-57 |
| **Parietal cortex** | 12 | 59-97 | 9-82.5 |
| **Prefrontal cortex** | 4 | 74-88 | 10-57 |

**Supplementary Table 3. Total number of segmented autofluorescent particles analyzed across brain regions and tissue compartments.** Autofluorescent particles were quantified separately in gray matter (GM) and white matter (WM) across the four examined brain regions. The final column indicates the total number of analyzed particles per region.

| Brain region | GM | WM | GM+WM |
|---|---|---|---|
| **Caudate-putamen** | 392,330 | 258,100 | 650,430 |
| **Hippocampus** | 570,574 | 141,719 | 712,293 |
| **Parietal cortex** | 430,044 | 298,059 | 728,103 |
| **Prefrontal cortex** | 406,544 | 168,430 | 574,974 |
| **TOTAL** | 1,799,492 | 866,308 | 2,665,800 |

**Supplementary Table 4. Number of ROIs (1,500 pixel × 1,500 pixel) analyzed for pixel correlation and spatial cell type mask overlap analysis.**

| Brain region | Sample | Nr. of ROIs in gray matter | Nr. of ROIs in white matter |
|---|---|---|---|
| **Caudate-putamen** | sample1 | 4 | 3 |
| | sample2 | 4 | 4 |
| | sample3 | 4 | 4 |
| | sample4 | 4 | 4 |
| **Hippocampus** | sample1 | 4 | 4 |
| | sample2 | 4 | 4 |
| | sample3 | 4 | 0 |
| | sample4 | 4 | 3 |
| **Parietal cortex** | sample1 | 4 | 4 |
| | sample2 | 4 | 3 |
| | sample3 | 4 | 0 |
| | sample4 | 4 | 4 |
| **Prefrontal cortex** | sample1 | 4 | 2 |
| | sample2 | 4 | 4 |
| | sample3 | 4 | 0 |
| | sample4 | 4 | 4 |

**Supplementary Table 5.** List of 64 antibodies targeting 56 different proteins, which were evaluated for staining patterns on fresh-frozen post-mortem human brain tissue using either conventional immunofluorescence (IF) or PhenoCyclerFusion (PCF) multiplexed IF (for pre-conjugated antibodies). Of the 64 antibodies evaluated, 37 underwent in-house DNA-barcode conjugation. The table includes antibody information (catalog number, vendor, species and clone ID), DNA-barcode conjugation status, dilution rate and tested tissue. The final column indicates the panel outcome.

| | Marker | | Catalog nr. | Vendor | Species (Clone) | Barcode conjugation | Dilution | Tissue tested | Panel outcome |
|---|---|---|---|---|---|---|---|---|---|
| 1 | AQP4 | Aquaporin 4 | AB282586 | Abcam | rabbit (EPR24281-65) | In-house | 200x | CAU, HIP, PAR, PFC | Included in panel, used for analysis |
| 2 | CD11c | Integrin subunit alpha X (ITGAX) | 4550114 | Akoya Biosciences | n.a. | Pre-conjugated | 50x | CAU, HIP, PAR, PFC | Included in panel, used for analysis |
| 3 | CD163 | Cluster of Differentiation 163 | HPA046404 | HPA | rabbit polyclonal | In-house | 100x | CAU, HIP, PAR, PFC | Included in panel, used for analysis |
| 4 | CD31 | Platelet and endothelial cell adhesion molecule 1 (PECAM-1) | 4250009 | Akoya Biosciences | n.a. | Pre-conjugated | 100x | CAU, HIP, PAR, PFC | Included in panel, used for analysis |
| 5 | CD45 | Protein tyrosine phosphatase receptor type C | 4150003 | Akoya Biosciences | n.a. | Pre-conjugated | 100x | CAU, HIP, PAR, PFC | Included in panel, used for analysis |
| 6 | CD68 | CD68 molecule | 14-0688-82 | Invitrogen | mouse (KP1) | In-house | 200x | CAU, HIP, PAR, PFC | Included in panel, used for analysis |
| 7 | ChAT | Choline O-Acetyltransferase | AB15468 | Merck | chicken polyclonal | In-house | 50x | CAU, HIP, PAR, PFC | Included in panel, used for analysis |
| 8 | CLDN5 | Claudin 5 | 352500 | Fisher Scientific | mouse (4C3C2) | In-house | 100x | CAU, HIP, PAR, PFC | Included in panel, used for analysis |
| 9 | Col IV | Collagen IV | 4550122 | Akoya Biosciences | n.a. | Pre-conjugated | 200x | CAU, HIP, PAR, PFC | Included in panel, used for analysis |
| 10 | GFAP-ab1 | Glial fibrillary acidic protein | AMAb91033 | Atlas Antibodies | mouse (CL2713) | In-house | 200x | CAU, HIP, PAR, PFC | Included in panel, used for analysis |
| 11 | GFAP-ab2 | Glial fibrillary acidic protein | ab218309 | Abcam | rabbit (EPR1034Y) | In-house | 200x | CAU, HIP, PAR, PFC | Included in panel, used for analysis |

| | | | | | | | | | |
|---|---|---|---|---|---|---|---|---|---|
| 12 | HLA-DR | Major histocompatibility complex, class II, DR alpha | 4250006 | Akoya Biosciences | n.a. | Pre-conjugated | 50x | CAU, HIP, PAR, PFC | Included in panel, used for analysis |
| 13 | MAP2 | Microtubule associated protein 2 | AMAb91375 | Atlas Antibodies | mouse (CL5420) | In-house | 100x | CAU, HIP, PAR, PFC | Included in panel, used for analysis |
| 14 | NEFL | Neurofilament light chain | AMAb91314 | Atlas Antibodies | mouse (CL4729) | In-house | 400x | CAU, HIP, PAR, PFC | Included in panel, used for analysis |
| 15 | Nes | Nestin | 33475 | Cell Signaling Technology | mouse (10C2) | In-house | 50x | CAU, HIP, PAR, PFC | Included in panel, used for analysis |
| 16 | NeuN | RNA binding Fox-1 homolog 3 (RBFOX3) | ABN78 | Merck | rabbit polyclonal | In-house | 100x | CAU, HIP, PAR, PFC | Included in panel, used for analysis |
| 17 | Olig2 | Oligodendrocyte transcription factor 2 | MABN50 | Merck | mouse (211F1.1) | In-house | 100x | CAU, HIP, PAR, PFC | Included in panel, used for analysis |
| 18 | P2Y12 | Purinergic receptor P2Y12 | APR-012 | AlomoneLabs | rabbit polyclonal | In-house | 50x | CAU, HIP, PAR, PFC | Included in panel, used for analysis |
| 19 | SOX9 | SRY-Box transcription factor 9 | ab76997-1002 | Abcam | mouse (3C10) | In-house | 100x | CAU, HIP, PAR, PFC | Included in panel, used for analysis |
| 20 | Syp | Synaptophysin | MA1-213 | Invitrogen | mouse (SY38) | In-house | 100x | CAU, HIP, PAR, PFC | Included in panel, used for analysis |
| 21 | TMEM119 | Transmembrane protein 119 | 853302 | BioLegend | mouse (A16075D) | In-house | 50x | CAU, HIP, PAR, PFC | Included in panel, used for analysis |
| 22 | Ubiquitin | Ubiquitin | sc-8017 | Santa Cruz Biotechnology | mouse (P4D1) | In-house | 50x | CAU, HIP, PAR, PFC | Included in panel, used for analysis |
| 23 | Vim | Vimentin | 677802 | BioLegend | mouse (O91D3) | In-house | 200x | CAU, HIP, PAR, PFC | Included in panel, used for analysis |
| 24 | CD21 | Complement C3d receptor 2 | 4150009 | Akoya Biosciences | n.a. | Pre-conjugated | 100x | CAU, HIP, PAR, PFC | Included in panel, used for analysis |
| 25 | CD3 | CD3 molecule | 4550103 | Akoya Biosciences | n.a. | Pre-conjugated | 100x | CAU, HIP, PAR, PFC | Included in panel, used for analysis |
| 26 | CD4 | CD4 molecule | 4550112 | Akoya Biosciences | n.a. | Pre-conjugated | 100x | CAU, HIP, PAR, PFC | Included in panel, used for analysis |
| 27 | HIF1a | Hypoxia-Inducible Factor 1-alpha | 4550069 | Akoya Biosciences | n.a. | Pre-conjugated | 100x | CAU, HIP, PAR, PFC | Included in panel, used for analysis |

| | | | | | | | | | |
|---|---|---|---|---|---|---|---|---|---|
| 28 | PD1 | Programmed cell death protein 1 | 4550038 | Akoya Biosciences | n.a. | Pre-conjugated | 100x | CAU, HIP, PAR, PFC | Included in panel, used for analysis |
| 29 | ALDH1L1+2 | Aldehyde Dehydrogenase 1 Family Member L | ab300510 | Abcam | rabbit (EPR25443-54) | In-house | 100x | CAU, HIP, PAR, PFC | Included in panel but excluded from analysis (too few positive cells) |
| 30 | CD141 | Thrombomodulin (THBD) | 240192 | Akoya Biosciences | n.a. | Pre-conjugated | 200x | CAU, HIP, PAR, PFC | Included in panel but excluded from analysis (unreliable staining) |
| 31 | CD38 | CD38 molecule | 232143 | Akoya Biosciences | n.a. | Pre-conjugated | 100x | CAU, HIP, PAR, PFC | Included in panel but excluded from analysis (too few positive cells) |
| 32 | EAAT1 | Solute carrier family 1 member 3 (SLC1A3 ) | 250-113 | SYSY Antibodies | rabbit polyclonal | In-house | 50x | CAU, HIP, PAR, PFC | Included in panel but excluded from analysis (unreliable staining) |
| 33 | FoxA2 | Forkhead box A2 | H00003170-M10 | Abnova / Life Technologies Europe | mouse (1C7) | In-house | 100x | CAU, HIP, PAR, PFC | Included in panel but excluded from analysis (unreliable staining) |
| 34 | Iba1 | Allograft Inflammatory Factor | ab220815 | Abcam | rabbit (EPR16588) | In-house | 100x | CAU, HIP, PAR, PFC | Included in panel but excluded from analysis (unreliable staining) |
| 35 | Ki67 | Marker of proliferation Ki-67 | 4250019 | Akoya Biosciences | n.a. | Pre-conjugated | 100x | CAU, HIP, PAR, PFC | Included in panel but excluded from analysis (too few positive cells) |
| 36 | MBP | Myelin basic protein | AMAb91064 | Atlas Antibodies / Merck | mouse monoclonal (CL2829) | In-house | 200x | CAU, HIP, PAR, PFC | Included in panel but excluded from analysis (unreliable staining) |
| 37 | PLP | Proteolipid protein | ab275751 | Abcam | rabbit (EPR23504-106) | In-house | 200x | CAU, HIP, PAR, PFC | Included in panel but excluded from analysis (unreliable staining) |
| 38 | PRKN | Parkin | sc-133167 | Santa Cruz Biotechnology | mouse (D1) | In-house | 50x | CAU, HIP, PAR, PFC | Included in panel but excluded from analysis (unreliable staining) |
| 39 | pTau | phosphorilated Microtubule associated protein tau | MN1020B | invitrogen | mouse (AT8) | In-house | 50x | CAU, HIP, PAR, PFC | Included in panel but excluded from analysis (unreliable staining) |

| | | | | | | | | | |
|---|---|---|---|---|---|---|---|---|---|
| 40 | SNCA | alpha-Synuclein | 69973SF | CellSignalin g Technology | rabbit (D37A6) | In-house | 50x | CAU, HIP, PAR, PFC | Included in panel but excluded from analysis (unreliable staining) |
| 41 | Sox10 | SRY-box transcription factor 10 | ab220078 | Abcam | rabbit (EPR4007-104) | In-house | 100x | CAU, HIP, PAR, PFC | Included in panel but excluded from analysis (unreliable staining) |
| 42 | TCF1 | T-cell factor 1 | 4550068 | Akoya Biosciences | n.a. | Pre-conjugated | 100x | CAU, HIP, PAR, PFC | Included in panel but excluded from analysis (unreliable staining) |
| 43 | TOMM20 | Translocase of outer mitochondrial membrane 20 | ab56783 | Abcam | mouse (4F3) | In-house | 50x | CAU, HIP, PAR, PFC | Included in panel but excluded from analysis (unreliable staining) |
| 44 | TREM2 | Triggering receptor expressed on myeloid cells 2 | ab318263 | Abcam | rabbit (EPR26209-22) | In-house | 50x | CAU, HIP, PAR, PFC | Included in panel but excluded from analysis (unreliable staining) |
| 45 | CD11b | Cyclin dependent kinase 1B | 101202 | BioLegend | rat (M1/70) | In-house | 100x | CAU | Failed conjugation |
| 46 | CNP | 2',3'-cyclic nucleotide 3' phosphodiesterase | AMAb91069 | Atlas Antibodies | mouse (CL2872) | In-house | 100x | PAR | Failed conjugation |
| 47 | Iba1 | Allograft Inflammatory Factor | PA5-27436 | Thermo Fisher | rabbit polyclonal | In-house | 100x | PAR | Failed conjugation |
| 48 | NEFH | neurofilament chain heavy | AMAb91025 | Atlas Antibodies | mouse (CL2671) | In-house | 100x | CAU | Failed conjugation |
| 49 | PSD-95 | Discs Large MAGUK Scaffold Protein 4 (DLG-4) | MA1-045 | Thermo Fisher | mouse (6G6-1C9) | In-house | 100x | PAR | Failed conjugation |
| 50 | S100B | S100 Calcium Binding Protein B | AMAb91038 | Atlas Antibodies | mouse (CL2720) | In-house | 50x | PAR | Failed conjugation |
| 51 | TH | Tyrosine hydrohylase | AB152 | Merck | rabbit polyclonal | In-house | 100x | PAR | Failed conjugation |
| 52 | ALDH1L1 | aldehyde dehydrogenase 1 family, member L1 | HPA050139 | Atlas Antibodies | rabbit polyclonal | Not conjugated | 100x | PAR | Not conjugated due to unreliable staining |
| 53 | ChAT | Choline O-Acetyltransferase | AMAb91129 | Atlas Antibodies | mouse (CL3169) | Not conjugated | 100x | PAR | Not conjugated due to unreliable staining |
| 54 | DRD2 | Dopamine D2 receptor (DD2R) | MABN53 | Sigma Aldrich | art (2B9) | Not conjugated | 50x | PAR | Not conjugated due to unreliable staining |

| | | | | | | | | | |
|---|---|---|---|---|---|---|---|---|---|
| 55 | GLUL | glutamate-ammonia ligase | AMAb91103 | Atlas Antibodies | mouse (CL3013) | Not conjugated | 100x | PAR | Not conjugated due to unreliable staining |
| 56 | Iba1 | Allograft Inflammatory Factor | ab221790 | Abcam | rabbit (EPR16589) | Not conjugated | 50x | PAR | Not conjugated due to unreliable staining |
| 57 | ITGAM | Integrin subunit alpha M | AMab90911 | Atlas Antibodies | mouse (CL1719) | Not conjugated | 100x | PAR | Not conjugated due to unreliable staining |
| 58 | MOG | Myelin Oligodendrocyte Glycoprotein | AMAb91067 | Atlas Antibodies | mouse (CL2858) | Not conjugated | 100x | PAR | Not conjugated due to unreliable staining |
| 59 | MRC1 / CD206 | Mannose Receptor C - Type 1 (other name: CD206) | ab64693 | Abcam | rabbit polyclonal | Not conjugated | 50x | PAR | Not conjugated due to unreliable staining |
| 60 | MRC1 / CD206 | Mannose Receptor C - Type 1 (other name: CD206) | sc-376108 | Santa Cruz Biotechnology | mouse (D-1) | Not conjugated | 100x | PAR | Not conjugated due to unreliable staining |
| 61 | NEFM | Neurofilament Medium Chain | AMAb91027 | Atlas Antibodies | mouse (CL2678) | Not conjugated | 100x | PAR | Not conjugated due to unreliable staining |
| 62 | NEFM | Neurofilament medium chain | AMAb91030 | Atlas Antibodies | mouse (CL2705) | Not conjugated | 100x | PAR | Not conjugated due to unreliable staining |
| 63 | PDGFRA | Platelet derived growth factor receptor alpha | sc-398206 | Santa Cruz Biotechnology | mouse (C-9) | Not conjugated | 100x | PAR | Not conjugated due to unreliable staining |
| 64 | PDGFRB | Platelet derived growth factor receptor beta | MA5-35288 | Thermo Fisher | rabbit (6E8C10) | Not conjugated | 50x | PAR | Not conjugated due to unreliable staining |

**Supplementary Table 6. A 28-plex DNA-barcoded antibody panel developed and used for multiplexed spatial proteomics analysis**, including target antigen and cell type, antibody vendor, catalog number, clone, working dilution, DNA barcode, fluorescence microscopy channel and excitation time used for image acquisition.

| | Protein target (Synonym) | | Target cell type | Ref. no. and vendor | | Species & clone | Dilution | In house conjugation (barcode) | Fluorescent channel [nm] | Cycle nr. | Excitation time [ms] |
|---|---|---|---|---|---|---|---|---|---|---|---|
| 1 | **AQP4** | Aquaporin 4 | Astrocytes (endfeet) | AB282586 | Abcam | rabbit (EPR24281-65) | 200x | Yes (#10) | 488 | 14 | 200 |
| 2 | **CD11c** | Integrin subunit alpha X (ITGAX) | Activated microglia, dendritic/myeloid cells | 232166 | Akoya Biosciences | n.a. | 50x | No (#24) | 647 | 1 | 300 |
| 3 | **CD163** | CD163 | Perivascular macrophages | HPA046404 | HPA | rabbit polyclonal | 100x | Yes (#25) | 488 | 1 | 300 |
| 4 | **CD21** | Complement C3d receptor 2 (CR2) | B cells | 4150009 | Akoya Biosciences | n.a. | 100x | No (#13) | 488 | 13 | 300 |
| 5 | **CD3** | CD3 | T cells | 232149 | Akoya Biosciences | n.a. | 100x | No (#15) | 647 | 8 | 300 |
| 6 | **CD31** | Platelet and endothelial cell adhesion molecule 1 (PECAM-1) | Endothelial cells | 232153 | Akoya Biosciences | n.a. | 100x | No (#32) | 550 | 10 | 250 |
| 7 | **CD4** | CD4 | helper T cells | 4550112 | Akoya Biosciences | n.a. | 100x | No (#3) | 647 | 11 | 250 |

| | | | | | | | | | | | |
|---|---|---|---|---|---|---|---|---|---|---|---|
| 8 | **CD45** | Protein tyrosine phosphatase receptor type C (PTPRC) | Blood derived immune cells | 4150003 | Akoya Biosciences | n.a. | 100x | No (#1) | 488 | 4 | 300 |
| 9 | **CD68** | CD68 | Macrophages, microglia | 14-0688-82 | Invitrogen | mouse (KP1) | 200x | Yes (#50) | 647 | 7 | 200 |
| 10 | **ChAT** | Choline O-Acetyltransferase | Cholinergic neurons | AB15468 | Merck | chicken polyclonal | 50x | Yes (#16) | 647 | 13 | 300 |
| 11 | **CLDN5** | Claudin 5 | Endothelial cells | 352500 | Fisher Scientific | mouse (4C3C2) | 100x | Yes (#41) | 550 | 7 | 300 |
| 12 | **COLIV** | Collagen IV | Basement membrane (vasculature) | 240070 | Akoya Biosciences | n.a. | 200x | No (#42) | 647 | 15 | 200 |
| 13 | **GFAP ab1** | Glial fibrillary acidic protein | Astrocytes | AMAb91033 | Atlas Antibodies | mouse (CL2713) | 200x | Yes (#28 ) | 488 | 8 | 120 |
| 14 | **GFAP ab2** | Glial fibrillary acidic protein | Astrocytes | ab218309 | Abcam | rabbit (EPR1034Y) | 200x | Yes (#20) | 550 | 5 | 75 |
| 15 | **HIF-1α** | Hypoxia-inducible factor 1-alpha | Hypoxia responses | 240128 | Akoya Biosciences | n.a. | 100x | No (#62) | 647 | 12 | 250 |
| 16 | **HLA-DR** | Major histocompatibility complex, class II, DR alpha | Activated antigen presenting cells | 232140 | Akoya Biosciences | n.a. | 50x | No (#26) | 550 | 6 | 350 |

| | | | | | | | | | | | |
|---|---|---|---|---|---|---|---|---|---|---|---|
| 17 | **MAP2** | Microtubule associated protein 2 | Neurons (somatodendritic) | AMAb91375 | Atlas Antibodies | mouse (CL5420) | 100x | Yes (#31) | 488 | 6 | 300 |
| 18 | **NEFL** | Neurofilament light chain | Neurons (axons) | AMAb91314 | Atlas Antibodies | mouse (CL4729) | 400x | Yes (#23) | 550 | 11 | 50 |
| 19 | **NES** | Nestin | Neuronal stem / progenitor cells, reactive astrocytes, endothelial cells | 33475 | Cell Signaling Tech. | mouse (10C2) | 50x | Yes (#43) | 647 | 9 | 200 |
| 20 | **NeuN** | RNA binding Fox-1 homolog 3 (RBFOX3) | Neurons | ABN78 | Merck | rabbit polyclonal | 100x | Yes (#19) | 488 | 12 | 200 |
| 21 | **OLIG2** | Oligodendrocyte transcription factor 2 | Oligodendrocyte lineage | MABN50 | Merck | mouse (211F1.1) | 100x | Yes (#27) | 647 | 14 | 300 |
| 22 | **PD-1** | Programmed cell death protein 1 (PDCD1) | Activation/exhaustion marker of lymphocytes | 4550038 | Akoya Biosciences | n.a. | 100x | No (#46) | 647 | 6 | 300 |
| 23 | **P2Y12** | Purinergic receptor $P2Y_{12}$ | Microglia (homeostatic) | APR-012 | AlomoneLabs | rabbit polyclonal | 50x | Yes (#21) | 647 | 10 | 500 |
| 24 | **SOX9** | SRY-box transcription factor 9 | Astrocytes | ab76997-1002 | Abcam | mouse (3C10) | 100x | Yes (#37) | 488 | 10 | 100 |
| 25 | **SYP** | Synaptophysin | Neurons (synapses) | MA1-213 | Invitrogen | mouse (SY38) | 100x | Yes (#54) | 550 | 13 | 50 |
| 26 | **TMEM119** | Transmembrane protein 119 | Microglia | 853302 | BioLegend | mouse (A16075D) | 50x | Yes (#5) | 550 | 4 | 350 |

| | | | | | | | | | | | |
|---|---|---|---|---|---|---|---|---|---|---|---|
| 27 | **Ubiquitin** | Ubiquitin | Protein turnover | sc-8017 | Santa Cruz Biotech. | mouse (P4D1) | 50x | Yes (#17) | 550 | 14 | 250 |
| 28 | **VIM** | Vimentin | Endothelial cells, Perivascular cells, Reactive astrocytes | 677802 | BioLegend | mouse (O91D3) | 200x | Yes (#29) | 550 | 12 | 100 |

**Supplementary Table 7. Experimental plan** used for the multiplexed immunofluorescence experiment performed on fresh frozen human brain tissue after optimized photobleaching experiment, showing order, fluorescent channel and exposure time for each antibody.

| Cycle | Atto550 channel | | Cy 5 / AF647 channel | | AF488 channel | |
|---|---|---|---|---|---|---|
| | Marker (barcode) | Exposure time [ms] | Marker (barcode) | Exposure time [ms] | Marker (barcode) | Exposure time [ms] |
| 1 | ALDH1L1-2 (BX002) | 250 | CD11c (BX024) | 300 | CD163 (BX025) | 300 |
| 2 | Iba1 (BX014) | 250 | pTau (BX119) | 250 | Sox10 (BX040) | 300 |
| 3 | Ki67 (BX047) | 350 | TOMM20 (BX072) | 250 | FoxA2 (BX022) | 200 |
| 4 | TMEM119 (BX005) | 350 | TCF1 (BX061) | 250 | CD45 (BX001) | 300 |
| 5 | GFAP ab2 (BX020) | 75 | TREM2 (BX033) | 300 | CD38 (BX007) | 250 |
| 6 | HLA-DR (BX026) | 350 | PD1 (BX046) | 250 | MAP2 (BX031) | 300 |
| 7 | Claudin5 (BX041) | 300 | CD68 (BX050) | 200 | SNCA (BX004) | 350 |
| 8 | PRKN (BX055) | 350 | CD3 (BX015) | 300 | GFAP ab1 (BX028) | 120 |
| 9 | empty | - | Nes (BX043) | 200 | PLP (BX049) | 100 |
| 10 | CD31 (BX032) | 250 | P2Y12 (BX021) | 500 | Sox9 (BX037) | 100 |
| 11 | NEFL (BX023) | 50 | CD4 (BX003) | 250 | MBP (BX034) | 100 |
| 12 | Vim (BX029) | 100 | HIF1a (BX062) | 250 | NeuN (BX019) | 200 |
| 13 | Syp (BX054) | 50 | ChAT (BX016) | 300 | CD21 (BX013) | 300 |
| 14 | Ubiquitin (BX017) | 250 | Olig2 (BX027) | 300 | AQP4 (BX010) | 200 |
| 15 | EAAT1 (BX070) | 200 | COL IV (BX042) | 200 | | |
| 16 | CD141/THBD (BX087) | 300 | | | | |

**Supplementary Table 8.** Mean marker intensity inside autofluorescence particle mask (log10 transformed and z-score normalized), for four analyzed brain regions; caudate-putamen (CAU), hippocampus (HIP), parietal cortex (PAR) and prefrontal cortex (PFC) for both gray matter (GM) and white matter (WM). For each brain region and compartment top four markers with highest mean intensity are marked in bold.

| | CAU | | HIP | | PAR | | PFC | |
|---|---|---|---|---|---|---|---|---|
| **Antibody** | **GM** | **WM** | **GM** | **WM** | **GM** | **WM** | **GM** | **WM** |
| **AQP4** | -0.413 | -0.836 | 0.307 | 0.752 | -0.298 | 0.151 | -0.304 | 0.287 |
| **CD11c** | -0.141 | -0.086 | 0.204 | 0.237 | 0.161 | 0.128 | -0.413 | -0.156 |
| **CD163** | -0.553 | -0.52 | 0.221 | 0.557 | -0.077 | 0.146 | **0.372** | 0.352 |
| **CD21** | -0.421 | -0.068 | 0.16 | 0.203 | 0.167 | -0.034 | 0.259 | 0.265 |
| **CD3** | **0.181** | **0.137** | 0.409 | 0.05 | -0.137 | 0.126 | -0.784 | -0.488 |
| **CD31** | -0.326 | -0.008 | 0.118 | 0.107 | 0.054 | 0.028 | 0.006 | -0.034 |
| **CD4** | -0.454 | -0.281 | 0.282 | 0.572 | -0.042 | 0.17 | -0.006 | 0.203 |
| **CD45** | -0.27 | -0.215 | 0.044 | 0.094 | 0.052 | 0.1 | 0.119 | 0.276 |
| **CD68** | **-0.013** | -0.101 | **0.605** | 0.702 | -0.497 | -0.264 | -0.34 | 0.084 |
| **ChAT** | -0.533 | -0.43 | 0.139 | 0.465 | **0.222** | 0.184 | 0.151 | **0.632** |
| **CLDN5** | -0.08 | -0.062 | 0.016 | **0.896** | -0.03 | -0.163 | 0.087 | -0.176 |
| **CollV** | **0.302** | **0.485** | -0.106 | -0.242 | **0.31** | **0.28** | -0.51 | -0.766 |
| **GFAP ab1** | -1.116 | -0.93 | 0.37 | 0.366 | 0.153 | **0.432** | 0.134 | **0.776** |
| **GFAP ab2** | -0.259 | -0.68 | **0.526** | **0.942** | -0.431 | -0.206 | -0.306 | 0.454 |
| **HIF1a** | -0.349 | 0.053 | 0.215 | 0.444 | 0.056 | -0.004 | -0.106 | -0.094 |
| **HLA-DR** | -0.168 | -0.152 | **0.451** | 0.599 | -0.232 | -0.147 | -0.2 | -0.102 |
| **MAP2** | -0.868 | **0.204** | 0.291 | -0.11 | 0.025 | 0.234 | 0.167 | 0.066 |
| **NEFL** | -0.766 | -0.801 | 0.157 | 0.518 | 0.151 | 0.249 | **0.459** | **0.845** |
| **Nes** | -0.381 | -0.252 | 0.067 | 0.518 | 0.053 | 0.072 | 0.035 | 0.008 |
| **NeuN** | -0.572 | -0.238 | 0.013 | 0.679 | 0.178 | -0.087 | 0.333 | 0.387 |
| **Olig2** | -0.373 | -0.24 | 0.105 | 0.242 | 0.13 | 0.069 | 0.019 | 0.369 |
| **P2Y12** | -0.987 | -0.725 | -0.253 | -0.158 | **0.403** | **0.54** | **0.693** | **0.645** |
| **PD1** | **-0.01** | **0.149** | 0.278 | 0.2 | 0.014 | 0.257 | -0.646 | -0.463 |
| **Sox9** | -0.173 | -0.004 | 0.289 | 0.283 | -0.115 | -0.043 | -0.12 | -0.089 |
| **Syp** | -0.93 | -0.296 | 0.077 | -0.47 | **0.29** | **0.384** | **0.42** | 0.5 |

| | | | | | | | | |
|---|---|---|---|---|---|---|---|---|
| **TMEM119** | -0.673 | -0.494 | **0.449** | **1** | -0.17 | -0.002 | -0.037 | -0.027 |
| **Ubiquitin** | -0.41 | -0.349 | 0.379 | 0.092 | -0.016 | 0.078 | -0.06 | 0.236 |
| **Vim** | -0.041 | -0.336 | 0.376 | **1.222** | -0.414 | -0.217 | -0.301 | 0.128 |
| **Average over all markers** | -0.386 | -0.253 | 0.221 | 0.384 | -0.001 | 0.0879 | -0.0314 | 0.147 |

**Supplementary Table 9.** Spearman correlation index for pixel correlation between autofluorescence in 488nm channel and all 28 protein markers for each brain region; CAU (caudate-putamen), hippocampus (HIP), parietal cortex (PAR) and prefrontal cortex (PFC), for gray matter (GM) and white matter (WM). Top four correlation indexes are highlighted.

| | CAU | | HIP | | PAR | | PFC | |
|---|---|---|---|---|---|---|---|---|
| | GM | WM | GM | WM | GM | WM | GM | WM |
| AQP4 | 0.327 | 0.106 | 0.432 | 0.367 | 0.226 | 0.037 | 0.390 | 0.220 |
| CD11c | 0.203 | 0.132 | 0.327 | 0.273 | 0.102 | -0.018 | 0.188 | 0.068 |
| CD163 | 0.189 | 0.072 | 0.351 | 0.273 | 0.224 | -0.017 | 0.429 | 0.115 |
| CD31 | 0.117 | 0.062 | 0.344 | 0.171 | 0.206 | 0.040 | 0.327 | 0.062 |
| CD45 | 0.149 | 0.069 | 0.275 | 0.197 | -0.184 | -0.228 | -0.043 | -0.212 |
| CD68 | 0.674 | 0.490 | **0.683** | 0.575 | 0.516 | 0.363 | **0.696** | 0.543 |
| CLDN5 | 0.143 | 0.045 | 0.162 | 0.266 | 0.102 | 0.001 | 0.215 | -0.030 |
| ChAT | 0.615 | 0.416 | 0.681 | 0.551 | 0.623 | 0.430 | **0.763** | 0.581 |
| ColIV | 0.062 | 0.037 | 0.209 | 0.121 | 0.090 | 0.010 | 0.175 | 0.036 |
| GFAP ab2 | 0.447 | 0.253 | 0.505 | 0.374 | 0.246 | 0.117 | 0.484 | 0.343 |
| GFAP ab1 | 0.160 | -0.011 | 0.362 | 0.210 | 0.268 | 0.027 | 0.475 | 0.126 |
| HLA-DR | 0.034 | 0.046 | 0.197 | 0.216 | 0.014 | 0.025 | 0.069 | 0.068 |
| MAP2 | 0.157 | 0.035 | 0.391 | 0.053 | 0.219 | -0.115 | 0.438 | 0.005 |
| NEFL | 0.560 | 0.369 | 0.582 | 0.468 | 0.491 | 0.165 | 0.681 | 0.423 |
| Nes | 0.380 | 0.274 | 0.527 | 0.454 | 0.372 | 0.210 | 0.529 | 0.348 |
| NeuN | 0.141 | 0.023 | 0.332 | 0.207 | 0.258 | -0.057 | 0.451 | 0.039 |
| Olig2 | 0.521 | 0.387 | 0.588 | 0.509 | 0.490 | 0.303 | 0.656 | 0.483 |
| P2Y12 | 0.576 | 0.390 | 0.601 | 0.506 | 0.485 | 0.257 | **0.701** | 0.476 |
| Sox9 | 0.029 | 0.021 | 0.095 | 0.037 | -0.022 | -0.048 | 0.029 | -0.033 |
| Syp | 0.552 | 0.341 | 0.486 | 0.444 | 0.434 | 0.197 | 0.609 | 0.446 |
| TMEM119 | 0.332 | 0.188 | 0.486 | 0.368 | 0.317 | 0.088 | 0.495 | 0.209 |

| | | | | | | | | |
|---|---|---|---|---|---|---|---|---|
| Ubiquitin | 0.413 | 0.373 | 0.513 | 0.468 | 0.382 | 0.243 | 0.534 | 0.422 |
| Vim | 0.478 | 0.330 | 0.528 | 0.493 | 0.312 | 0.183 | 0.481 | 0.419 |

**Supplementary Table 10.** Information on mixed-effects models used to assess statistical significance when multiple measurements were obtained from the same donor.

| Figure reference | Outcome variable | Fixed effects | Formula | Minimum data rule |
|---|---|---|---|---|
| Figure 1C | Log2-transformed AF particle MFI for AF488, AF550, or AF647 | Tissue (GM vs WM) and brain region | log2_MFI ~ C(tissue) + C(brain_region) + (1 \| donor) | No explicit minimum |
| Figures 1D–E | Raw AF particle size (D) or raw AF particle density (E) | Tissue (GM vs WM) and brain region | value ~ C(tissue) + C(brain_region) + (1 \| donor) | No explicit minimum |
| Figures 1F–H | Raw AF488 MFI (F,G), AF particle density (H) | Donor age (F,H) or PMD and brain region (G) | model_value ~ predictor + C(brain_region) + (1 \| donor) | No explicit minimum |
| Figure 8D, K | $\log 2(F_{AF+,CC+}/F_{ROI,CC+})$ (D), $\log 2(F_{AF+,CT+}/F_{ROI,CT+})$ (K) | Intercept only | log2_enrichment ~ 1 + (1 \| donor) | ≥3 donors and ≥4 observations |
| Figure 8E, L | $F_{AF+,CC+}$, $F_{ROI,CC+}$ (E), $F_{AF+,CT+}$, $F_{ROI, CT+}$ (L) | Analysis type (AF versus ROI) and brain region | distribution_percent ~ C(analysis_type, reference="ROI") + C(brain_region) + (1 \| donor) | ≥2 donors and both AF and ROI values |
| Figure 8G, N | $F_{CC+,AF+}$, $F_{CC-,AF+}$ (G), $F_{CT+,AF+}$, $F_{CT-,AF+}$ (N) | Location (inside versus outside), brain region | AF_occupancy_percent ~ C(location, reference="Outside") + C(brain_region) + (1 \| donor) | ≥2 donors and both AF and ROI values |

### SUPPLEMENTARY FIGURES

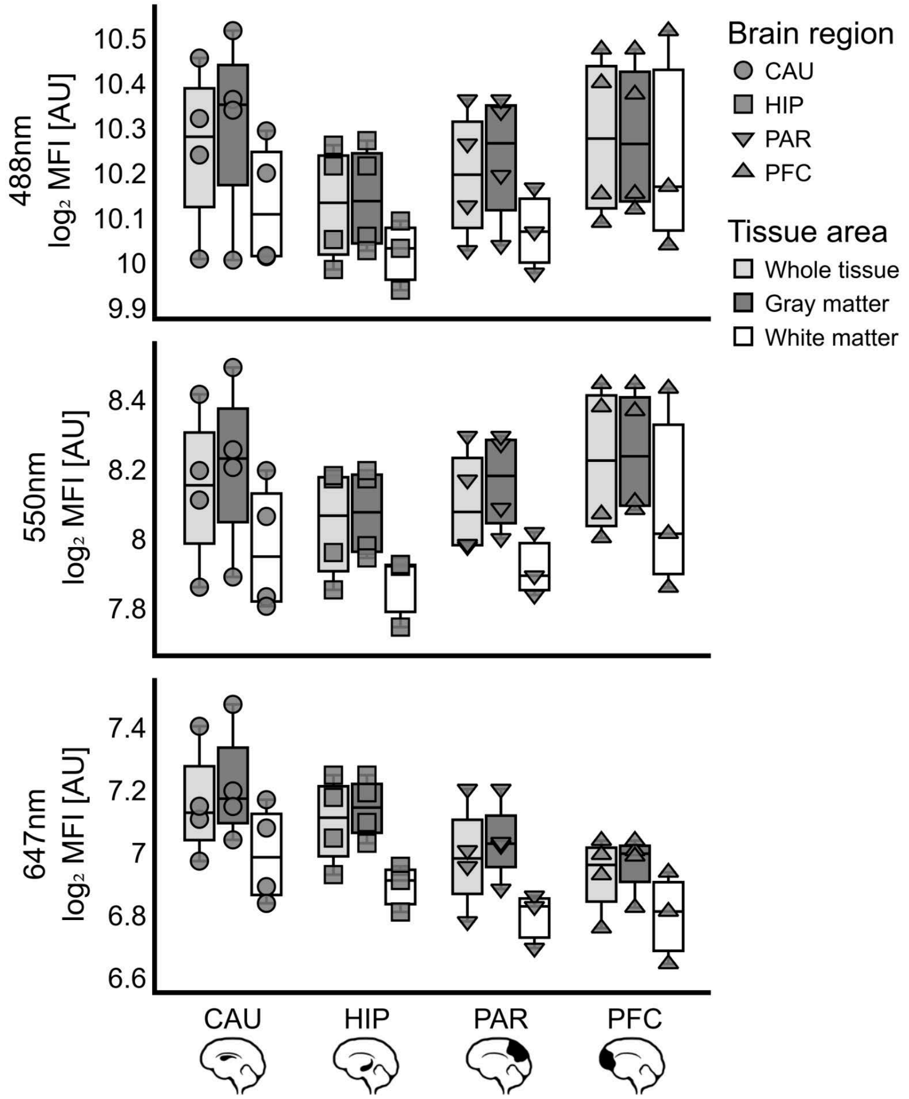


**Supplementary Figure 1. Mean autofluorescence intensity across brain regions and tissue compartments.** Boxplots summarize the mean autofluorescence (AF) intensity in the 488 nm, 550 nm, and 647 nm channels, quantified using AF particle segmentation masks in whole tissue sections (light gray) and separately in gray matter (GM, dark gray) and white matter (WM, white) across the four analyzed brain regions. Each point represents one donor sample, and point shape indicates the brain region. Sample size: *n* (CAU) = 4, *n* (HIP) = 4, *n* (PAR) = 4, *n* (PFC) = 4.

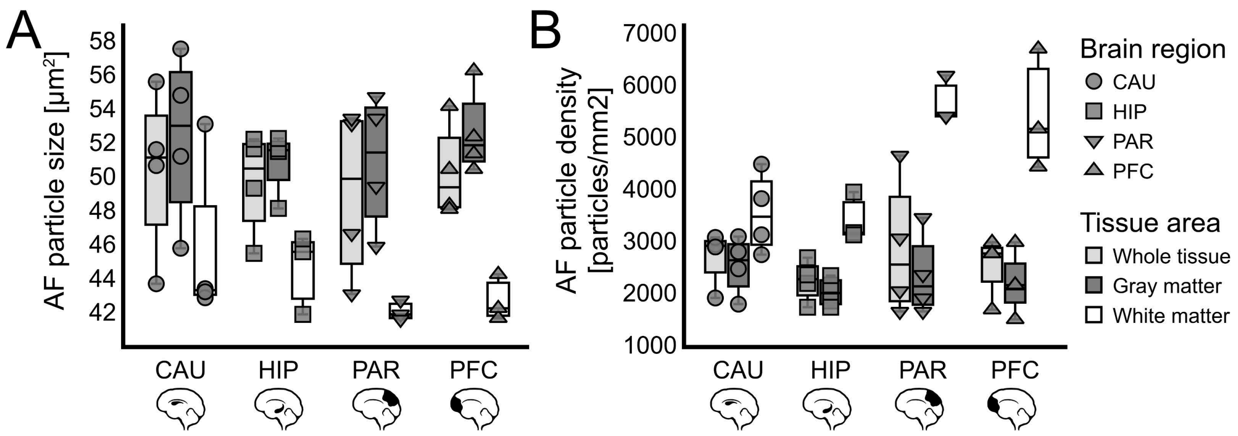


**Supplementary Figure 2. Autofluorescent (AF) particle size and density across brain regions and tissue compartments.** Boxplots summarize the mean AF particle size **(A)** and AF particle density **(B)**, quantified from autofluorescent particle segmentation masks in the 488 nm channel in whole tissue sections (light gray) and separately in gray matter (GM, dark gray) and white matter (WM, white) across the four analyzed brain regions. Each point represents one donor sample, and point shape indicates the brain region. Sample size: *n* (CAU) = 4, *n* (HIP) = 4, *n* (PAR) = 4, *n* (PFC) = 4.

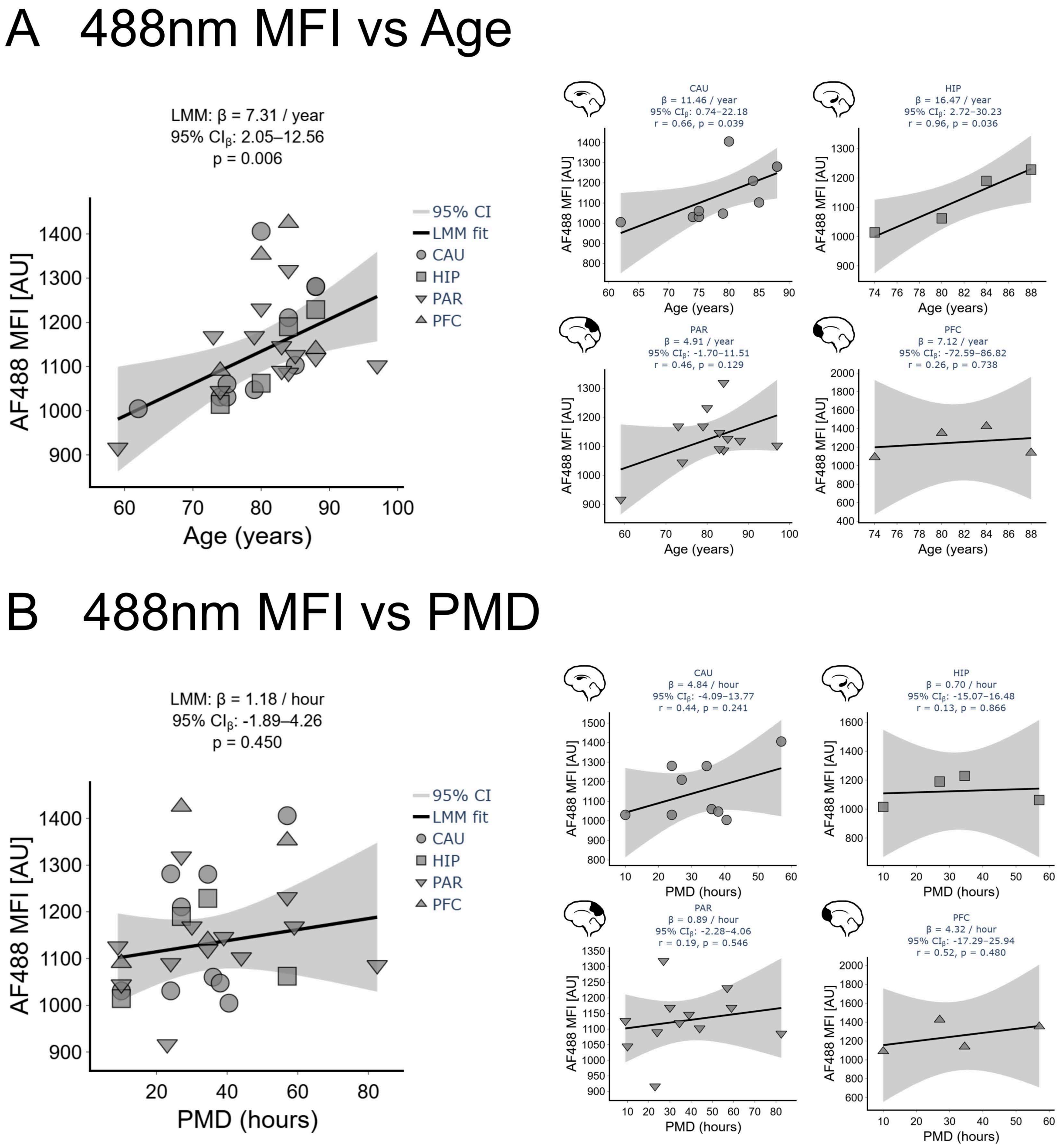


**Supplementary Figure 3. Association between autofluorescent (AF) particle intensity and donor characteristics.** Scatter plots showing the correlation between the mean AF particle intensity (MFI) in the 488 nm channel and donor age (**A**) or post-mortem delay (PMD; **B**). Large scatter plots show correlations across all analyzed brain regions combined, whereas smaller plots show region-specific correlations, indicated by brain region icons. Each point represents one donor sample, and point shape indicates the brain region. Sample size: *n* (CAU) = 10, *n* (HIP) = 4, *n* (PAR) = 12, *n* (PFC) = 4.

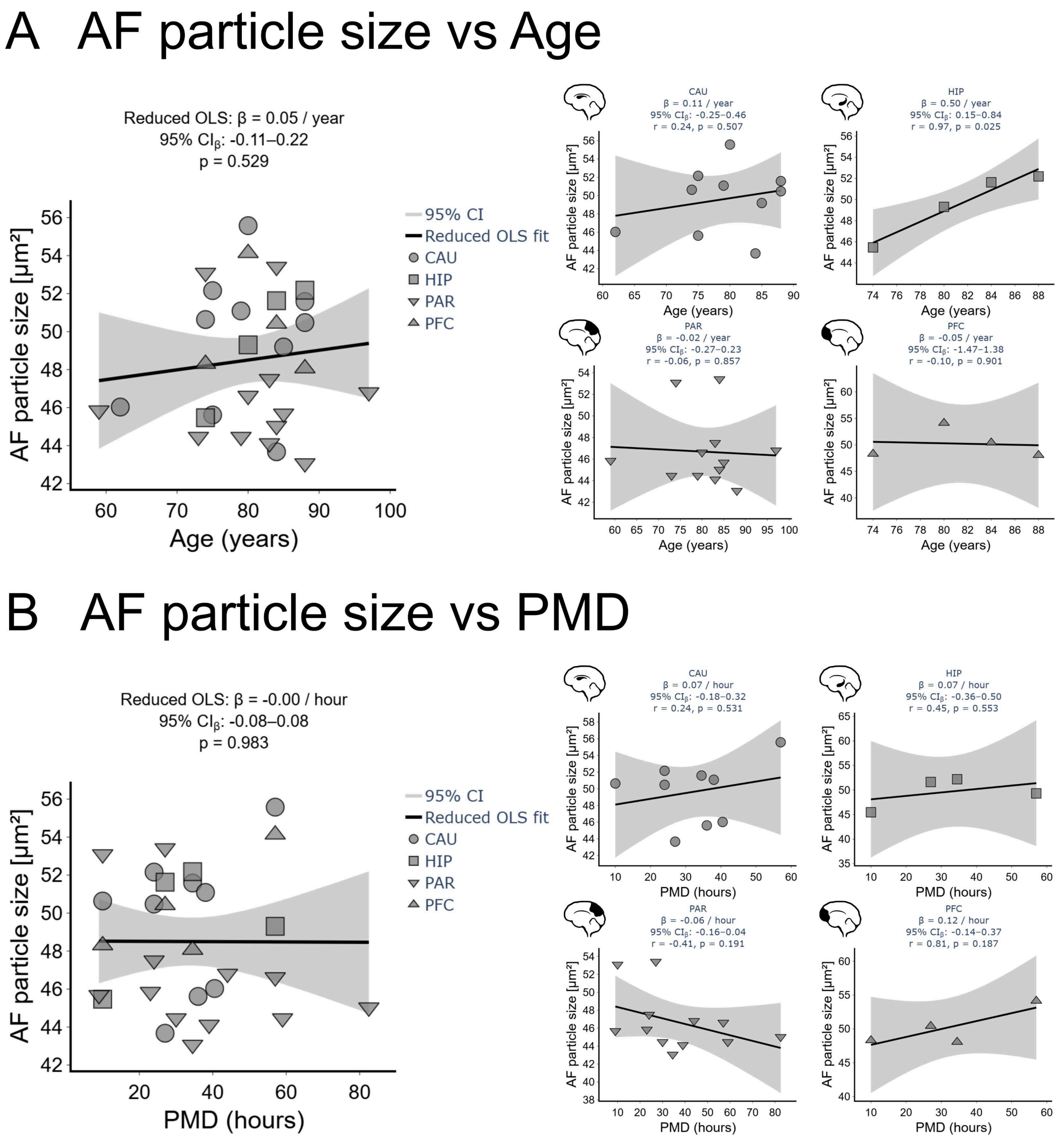


**Supplementary Figure 4. Association between autofluorescent (AF) particle size and donor characteristics.** Scatter plots showing the correlation between mean AF particle size and donor age (**A**) or post-mortem delay (PMD; **B**). Large scatter plots show correlations across all analyzed brain regions combined, whereas smaller plots show region-specific correlations, indicated by brain region icons. Each point represents one donor sample, and point shape indicates the brain region. Sample size: *n* (CAU) = 10, *n* (HIP) = 4, *n*  (PAR) = 12, *n* (PFC) = 4.

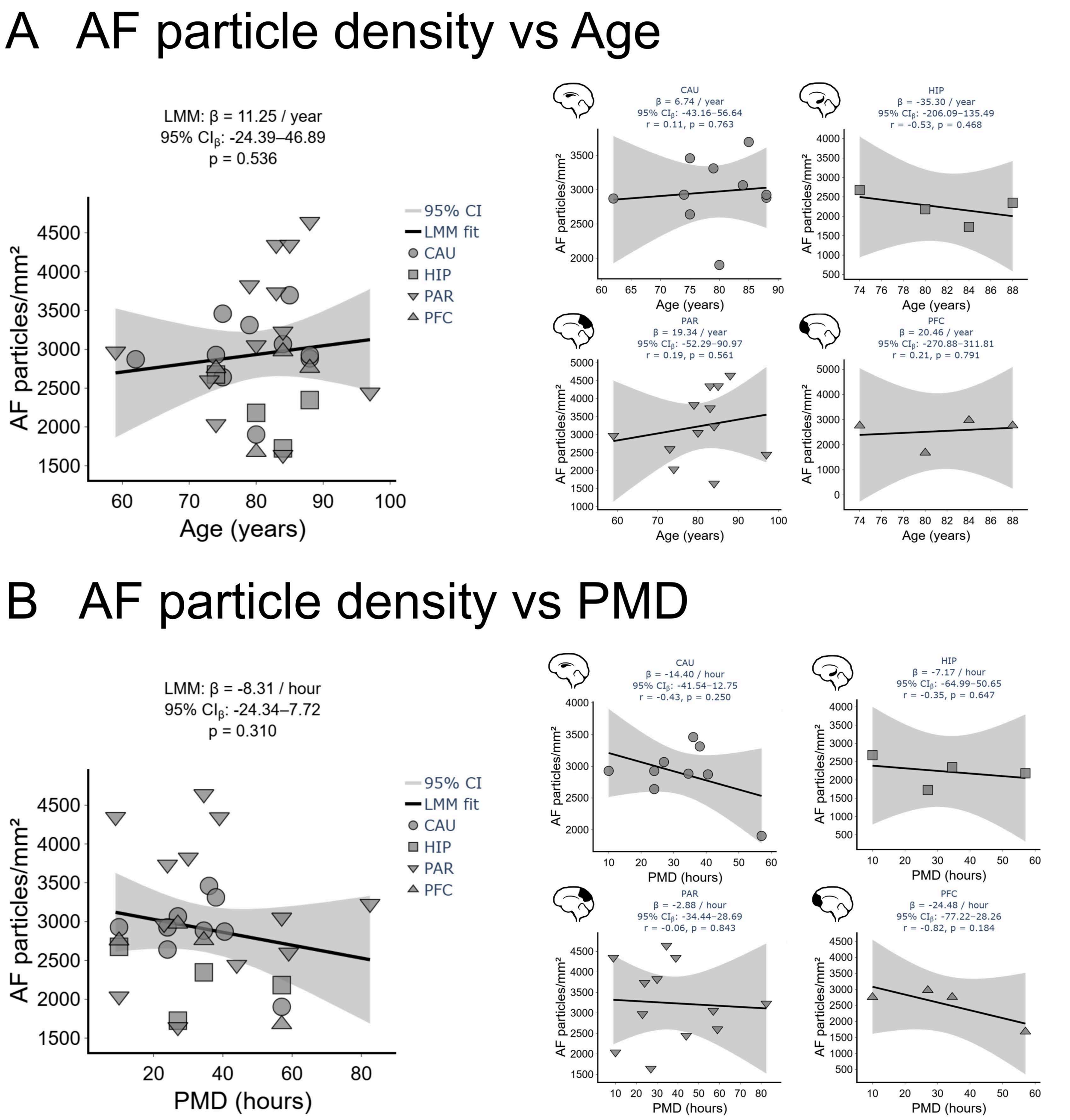


**Supplementary Figure 5. Association between autofluorescent (AF) particle density and donor characteristics.** Scatter plots showing the correlation between mean AF particle density and donor age (**A**) or post-mortem delay (PMD; **B**). Large scatter plots show correlations across all analyzed brain regions combined, whereas smaller plots show region-specific correlations, indicated by brain region icons. Each point represents one donor sample, and point shape indicates the brain region. Sample size: *n* (CAU) = 10, *n* (HIP) = 4, *n* (PAR) = 12, *n* (PFC) = 4.

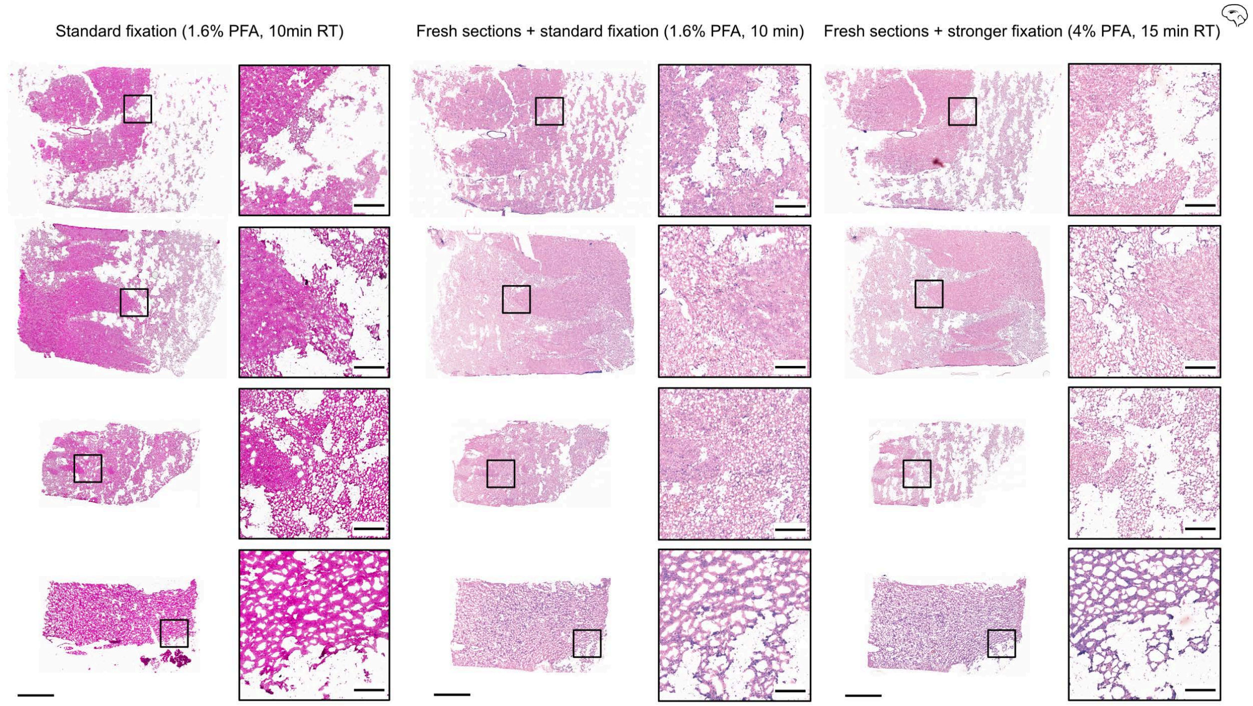


**Supplementary Figure 6. Fresh sectioning or stronger fixation does not prevent tissue damage induced by standard photobleaching in fresh-frozen human brain.** Representative hematoxylin and eosin (H&E)-stained fresh-frozen caudate-putamen sections following the standard photobleaching protocol. The left column shows the pronounced tissue damage induced by photobleaching using standard fixation protocol (1.6% PFA for 10 min). The middle and right columns show consecutive freshly sectioned tissue sections processed using either the standard fixation protocol or stronger fixation protocol (4% PFA for 15 min). Neither fresh sectioning nor stronger fixation prevented photobleaching-induced tissue damage. Whole-section images and corresponding magnified regions are shown. Scale bars: 2 mm (whole section images), 300 µm (magnified regions).

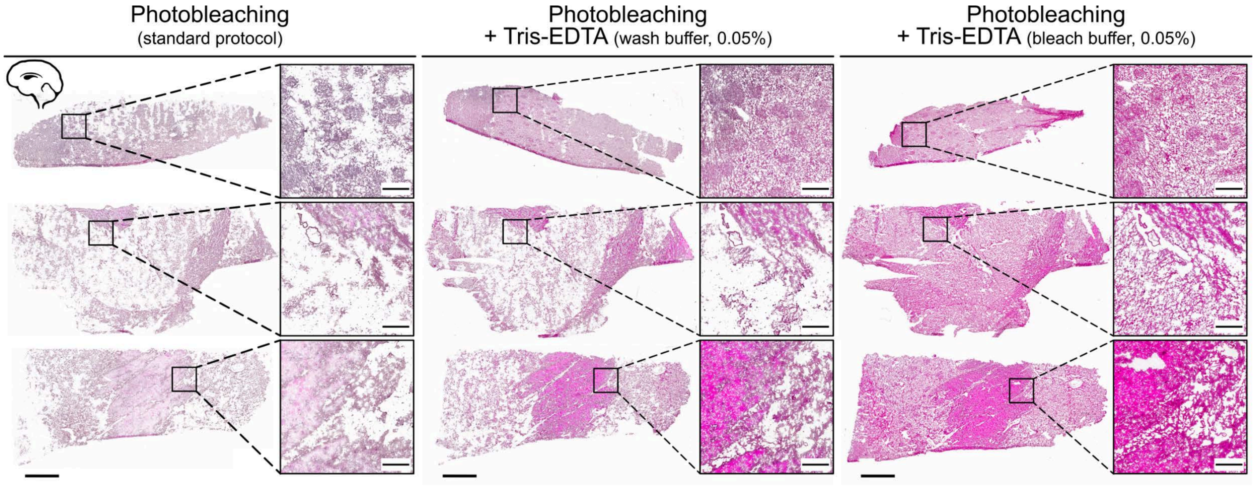


**Supplementary Figure 7. Tris-EDTA supplementation of the photobleaching buffer preserves tissue integrity during photobleaching of fresh-frozen human brain.** Representative H&E-stained caudate-putamen sections following photobleaching using the standard protocol alone (left), supplementation of the wash buffer with 0.05% Tris-EDTA prior to photobleaching (middle), or supplementation of the photobleaching buffer with 0.05% Tris-EDTA (right). Addition of Tris-EDTA directly to the photobleaching buffer preserved tissue integrity and reduced photobleaching-induced damage compared with the standard protocol or wash-buffer supplementation. Whole-section images and corresponding magnified regions are shown. Scale bars: 2 mm (whole section images), 300 µm (magnified regions).

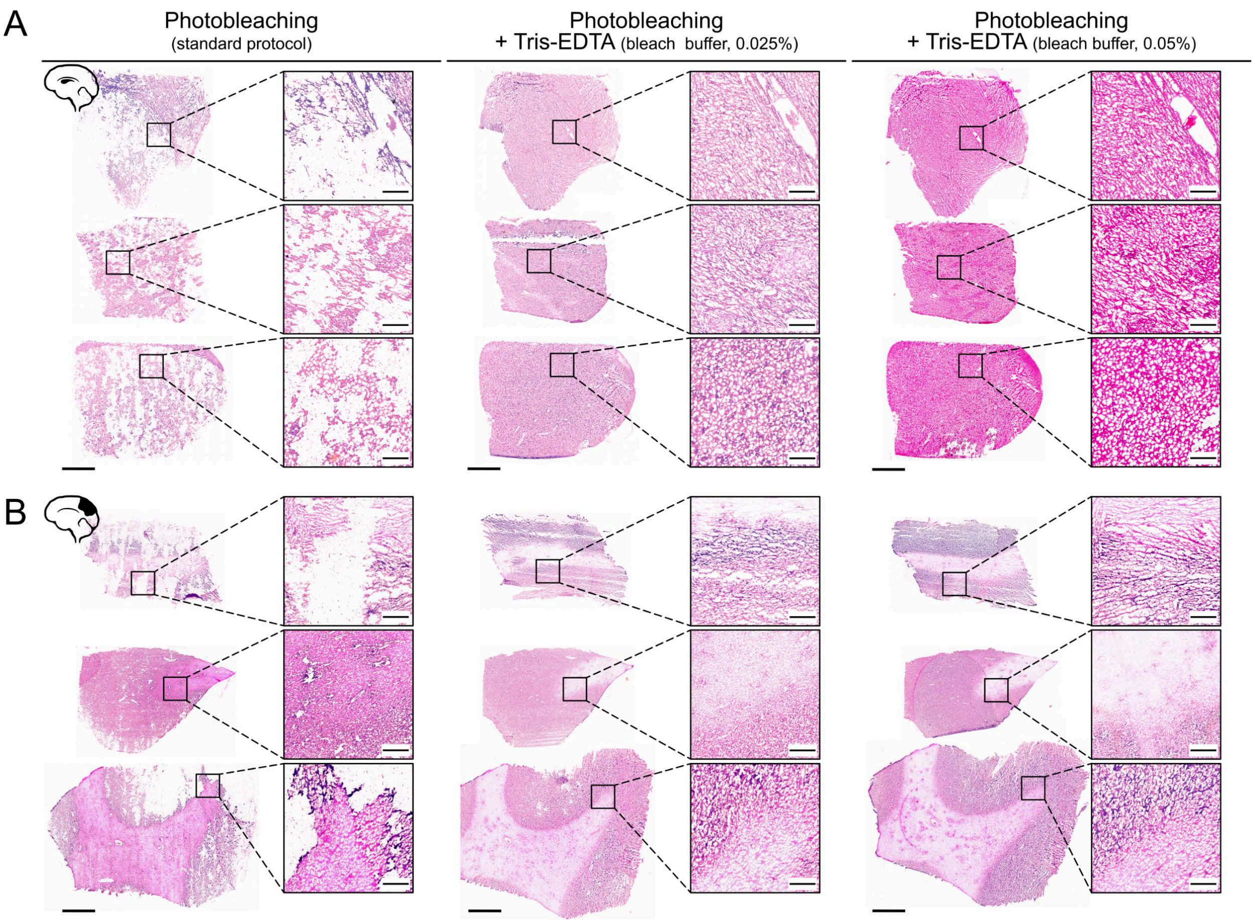


**Supplementary Figure 8. Tissue preservation during photobleaching with different concentrations of Tris-EDTA in fresh-frozen human brain.**

**A.** Representative H&E-stained caudate-putamen sections following photobleaching without Tris-EDTA (left), with 0.025% Tris-EDTA (1:200 dilution) (middle), or with 0.05% Tris-EDTA (1:100 dilution) (right) added to the photobleaching buffer.

**B.** Representative H&E-stained parietal cortex sections following the same photobleaching conditions.

For both brain regions, Tris-EDTA was added only during the first of two photobleaching incubations. Both tested concentrations improved tissue preservation compared with photobleaching without Tris-EDTA. Whole-section images and corresponding magnified regions are shown. Scale bars: 2 mm (whole section images), 300 μm (magnified regions).

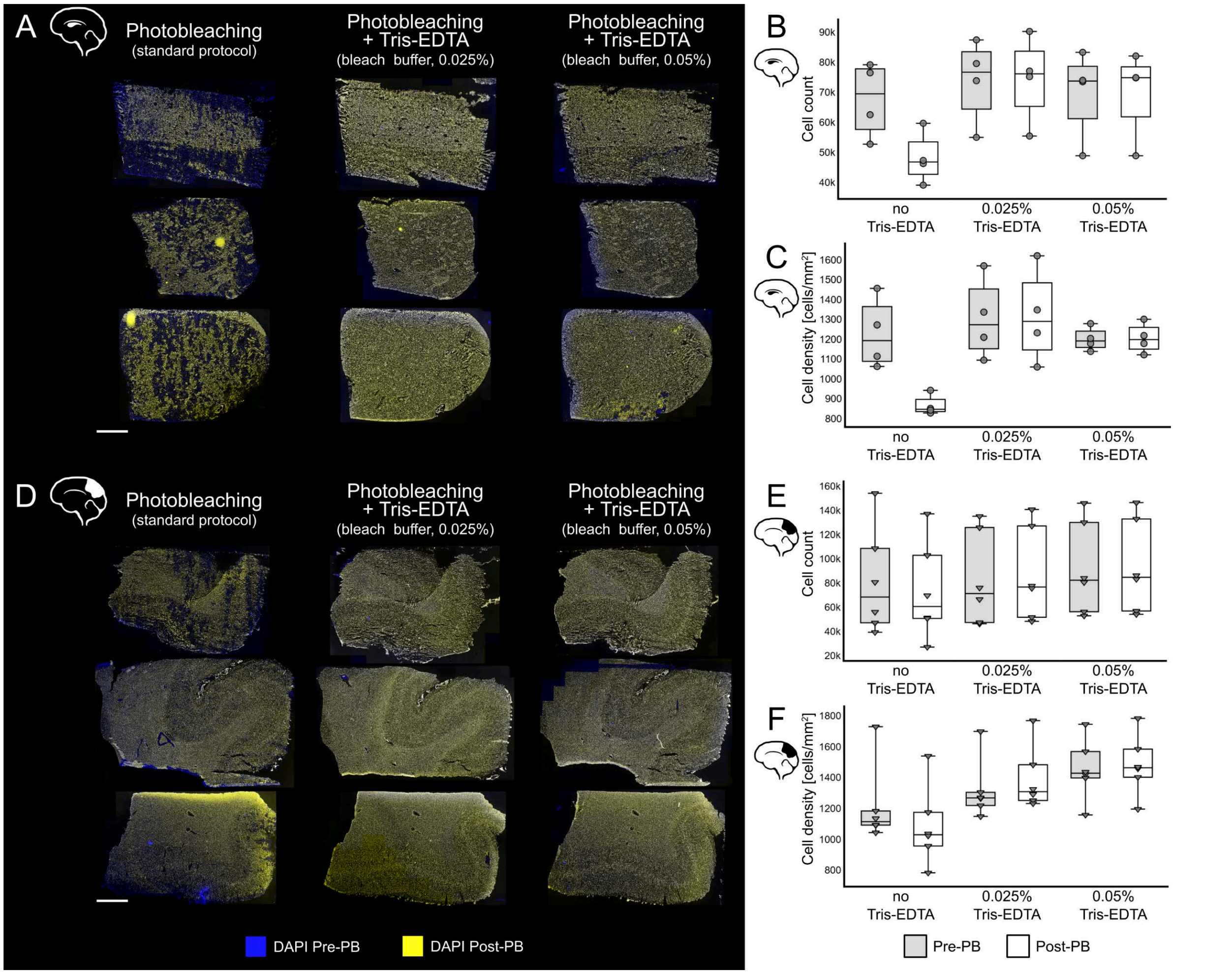


**Supplementary Figure 9. Nuclear staining-based assessment and quantification of photobleaching-induced cell loss demonstrates preservation of tissue integrity by Tris-EDTA supplementation.**

**A.** Spatially aligned DAPI staining images of caudate-putamen acquired before photobleaching (Pre-PB, blue) and after photobleaching (Post-PB, yellow) from three consecutive caudate-putamen sections, each subjected to one of the three photobleaching conditions: without Tris-EDTA (left), 0.025% Tris-EDTA (middle), or 0.05% Tris-EDTA (right) added to the photobleaching buffer. For each condition, pre- and post-photobleaching images were acquired from the same tissue section and spatially aligned. Overlay images visualize changes in cellular content and tissue architecture following treatment. Scale bar: 2 mm.

**B–C.** Boxplots showing the cell count **(B)** and cell density **(C)** before and after photobleaching in caudate-putamen sections across three photobleaching conditions: no Tris-EDTA, 0.025% Tris-EDTA, and 0.05% Tris-EDTA. Sample size: $n$ (CAU)= 4.

**D.** Spatially aligned DAPI staining images of parietal cortex sections before photobleaching (Pre-PB, blue) and after photobleaching (Post-PB, yellow) following photobleaching without Tris-EDTA (left), with 0.025% Tris-EDTA (middle), or with 0.05% Tris-EDTA (right) added to the photobleaching buffer. Scale bar: 2 mm.

**E–F.** Boxplots showing the cell count **(E)** and cell density **(F)** before and after photobleaching in parietal cortex sections across the three photobleaching conditions. Sample size: $n$ (PAR)= 6.

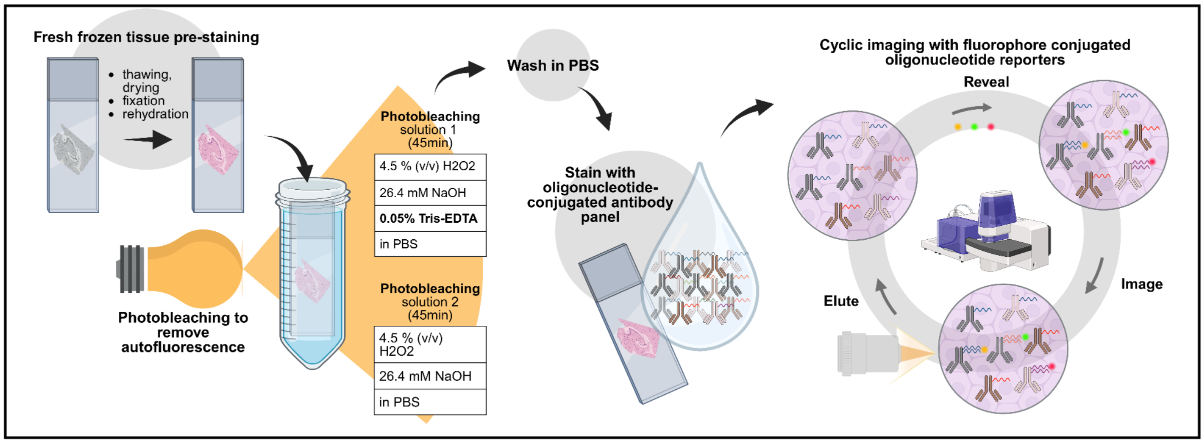


**Supplementary Figure 10. Schematic overview of the optimized photobleaching workflow developed for fresh-frozen brain tissue and its integration into the multiplexed immunofluorescence protocol.** Following tissue preparation and fixation, sections underwent two consecutive 45-min photobleaching steps under high-intensity LED illumination. The first photobleaching solution contained 0.05% Tris–EDTA to preserve tissue integrity during bleaching. After photobleaching, sections were stained with the DNA-barcoded antibody panel, and processed through iterative cycles of reporter hybridization, fluorescence imaging, and reporter removal to generate multiplexed spatial proteomic data. (Figure created in BioRender. Ayoglu, B. (2026) BioRender.com)

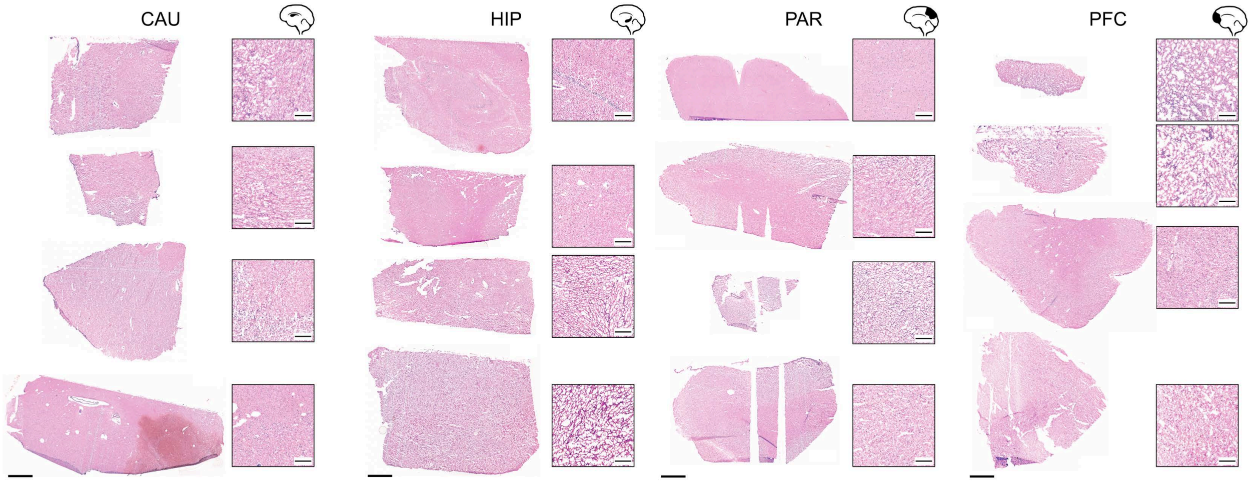


**Supplementary Figure 11. Histological assessment of fresh-frozen human brain tissue integrity following the optimized photobleaching workflow and multiplexed immunofluorescence staining.** Hematoxylin and eosin (H&E)-stained fresh-frozen human brain sections from caudate-putamen (CAU), hippocampus (HIP), parietal cortex (PAR), and prefrontal cortex (PFC), demonstrating preservation of tissue morphology after completion of the optimized photobleaching workflow and subsequent multiplexed immunofluorescence staining and imaging. Whole-section images and corresponding magnified regions are shown. Scale bars: 2 mm (whole section images), 300 μm (magnified regions).

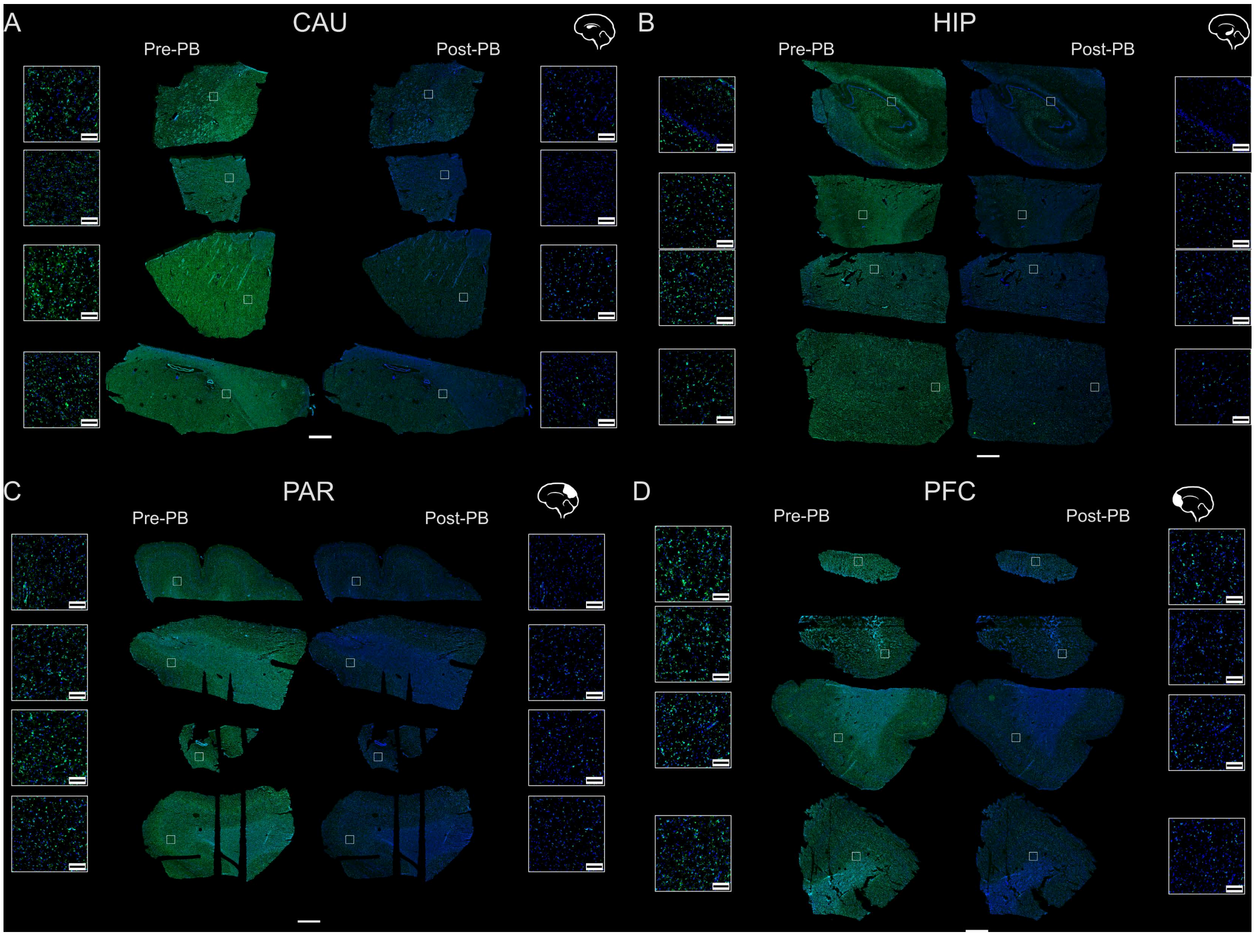


**Supplementary Figure 12. The optimized photobleaching protocol with 0.05% Tris-EDTA substantially reduces autofluorescence across brain regions.** Representative whole-section autofluorescence images acquired before (Pre-PB) and after photobleaching (Post-PB) in the 488 nm channel from caudate-putamen **(A)**, hippocampus **(B)**, parietal cortex **(C)** and prefrontal cortex **(D)**, demonstrating robust autofluorescence suppression across all analyzed brain regions. Whole-section images and corresponding magnified regions are shown. Scale bars: 2 mm for (whole-section images), and 150 µm (for magnified regions).

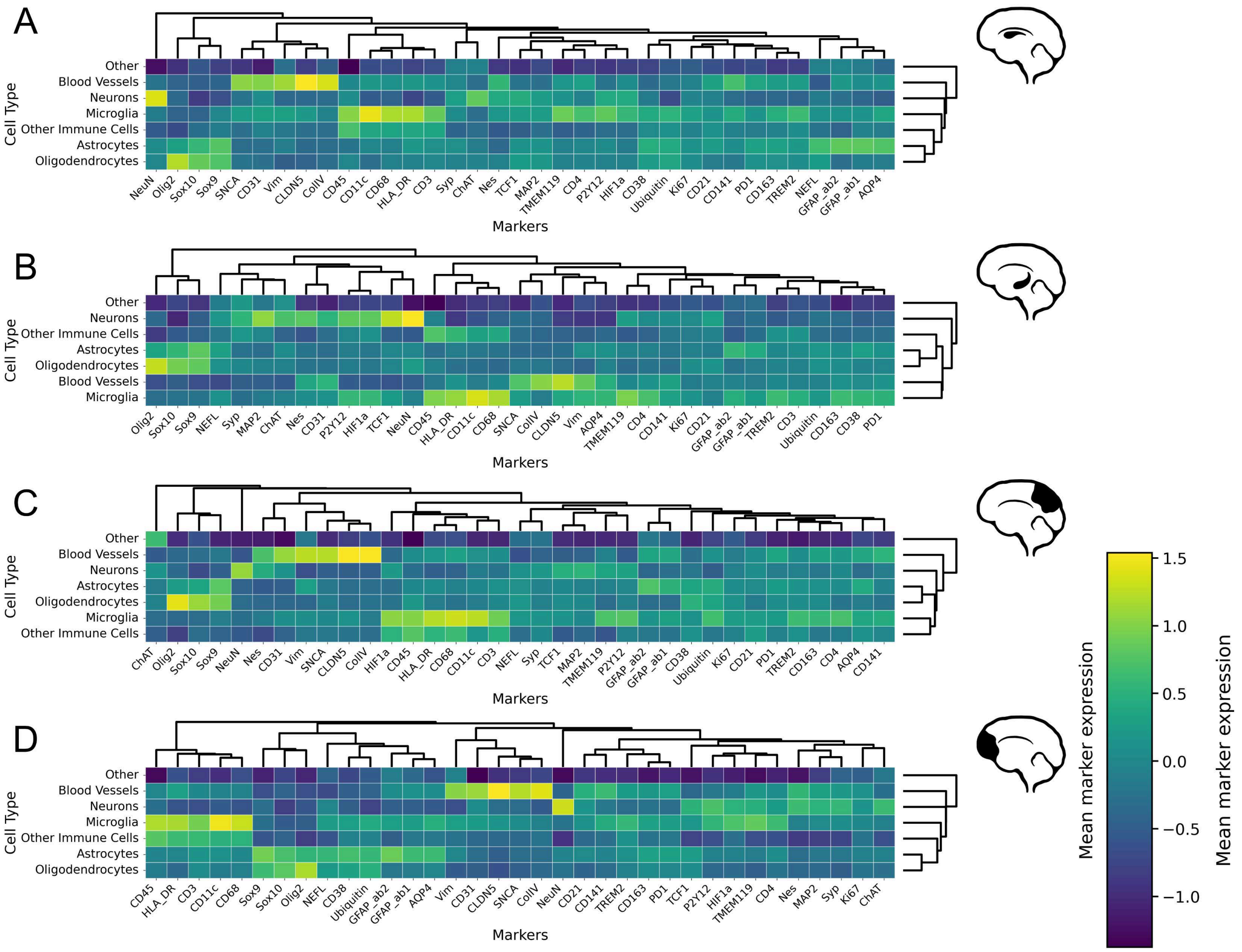


**Supplementary Figure 13. Clustered heatmaps of mean marker expression across assigned cell types in the analyzed brain regions.** Columns represent the 28 markers included in the antibody panel, whereas rows represent the assigned cell types. Color intensity represents relative mean marker expression after $\log_{10}$ transformation and z-score normalization. Heatmaps are shown separately for caudate-putamen **(A)**, hippocampus **(B)**, parietal cortex **(C)**, and prefrontal cortex **(D)**. Sample size: *n* (CAU) = 1, *n* (HIP) = 1, *n* (PAR) = 1, *n* (PFC) = 1.

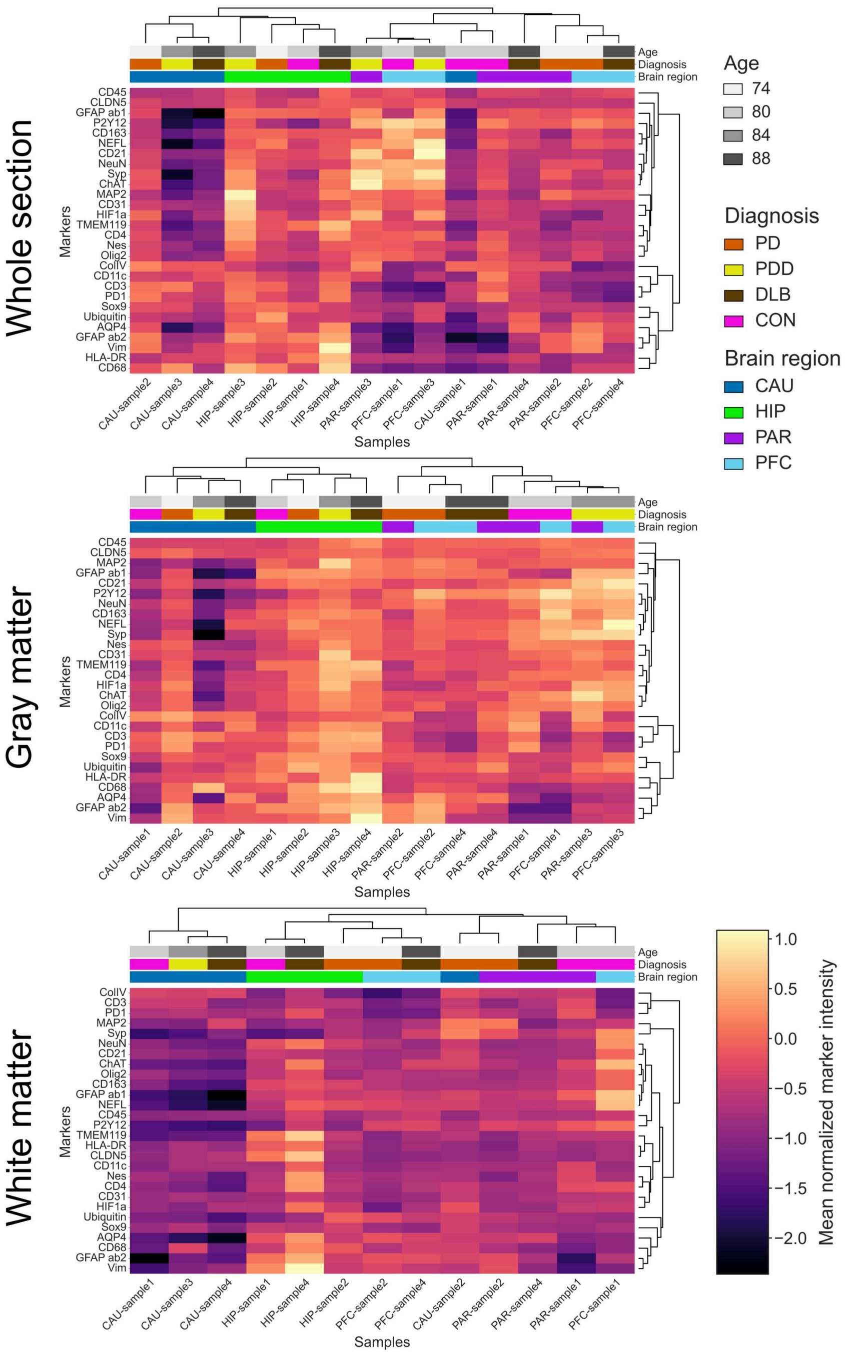


**Supplementary Figure 14. Mean protein marker intensities within autofluorescent (AF) particle masks across brain regions and tissue compartments.** Clustered heatmaps showing mean protein-marker intensities measured within segmented AF particle masks in whole tissue sections (top), gray matter (middle), and white matter (bottom). Columns represent individual donor samples, whereas rows represent the 28 markers included in the antibody panel. Sample annotations indicate tissue compartment, donor age, diagnosis, and brain region. Color intensity represents $\log_{10}$-transformed, z-score-normalized mean protein marker intensity within AF particle masks. Sample size: *n* (CAU) = 4, *n* (HIP) = 4, *n* (PAR) = 4, *n* (PFC) = 4.

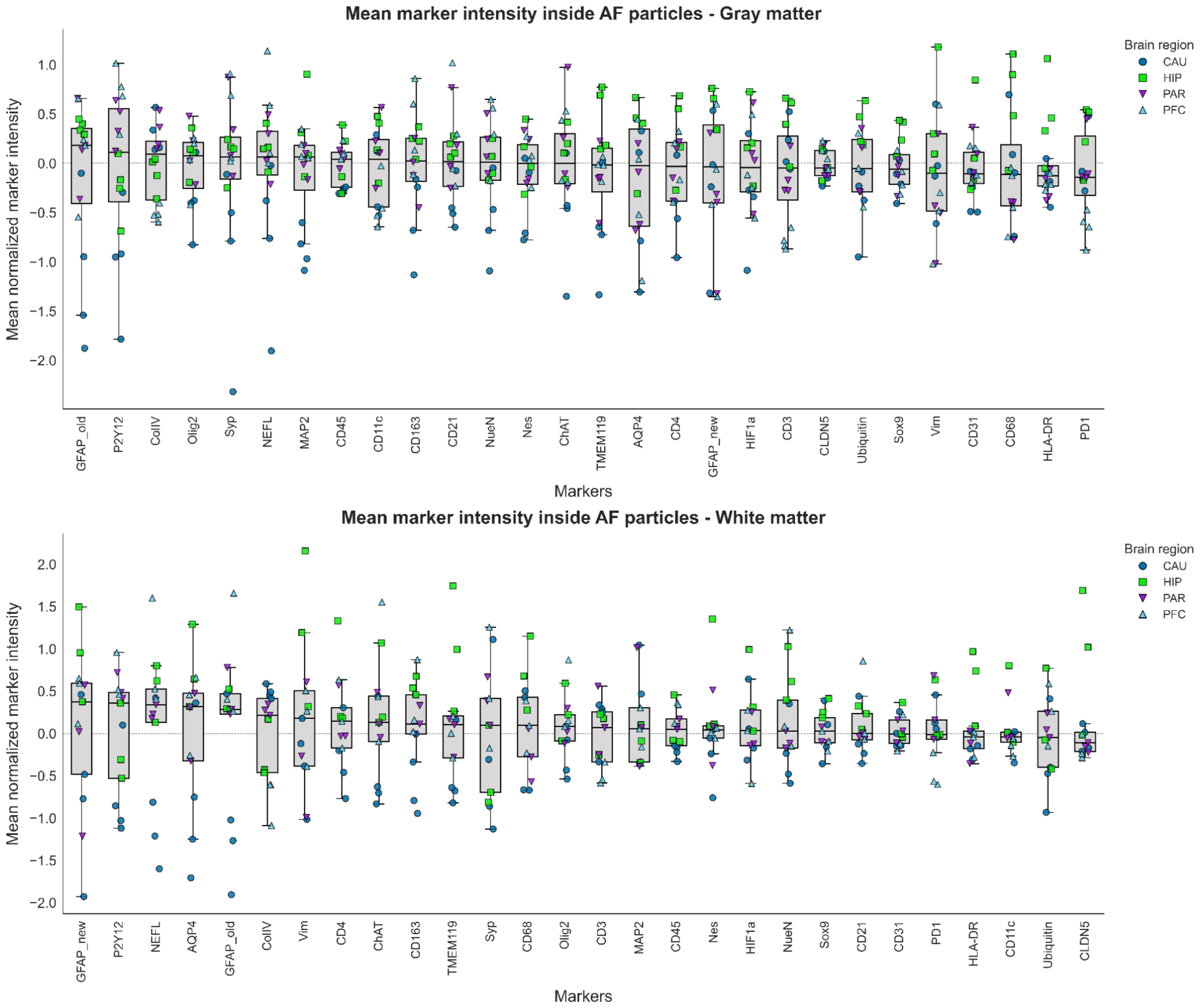


**Supplementary Figure 15. Mean protein marker intensities within autofluorescent (AF) particle masks across gray and white matter.** Box-and-whisker plots showing mean protein marker intensities measured within segmented AF particle masks in gray matter (top), and white matter (bottom). Protein markers on the x-axis are ordered from highest to lowest mean intensity (left to right). Each point represents one donor sample, with point colour and shape indicating the brain region. Sample size: *n* (CAU) = 4, *n* (HIP) = 4, *n* (PAR) = 4, *n* (PFC) = 4.

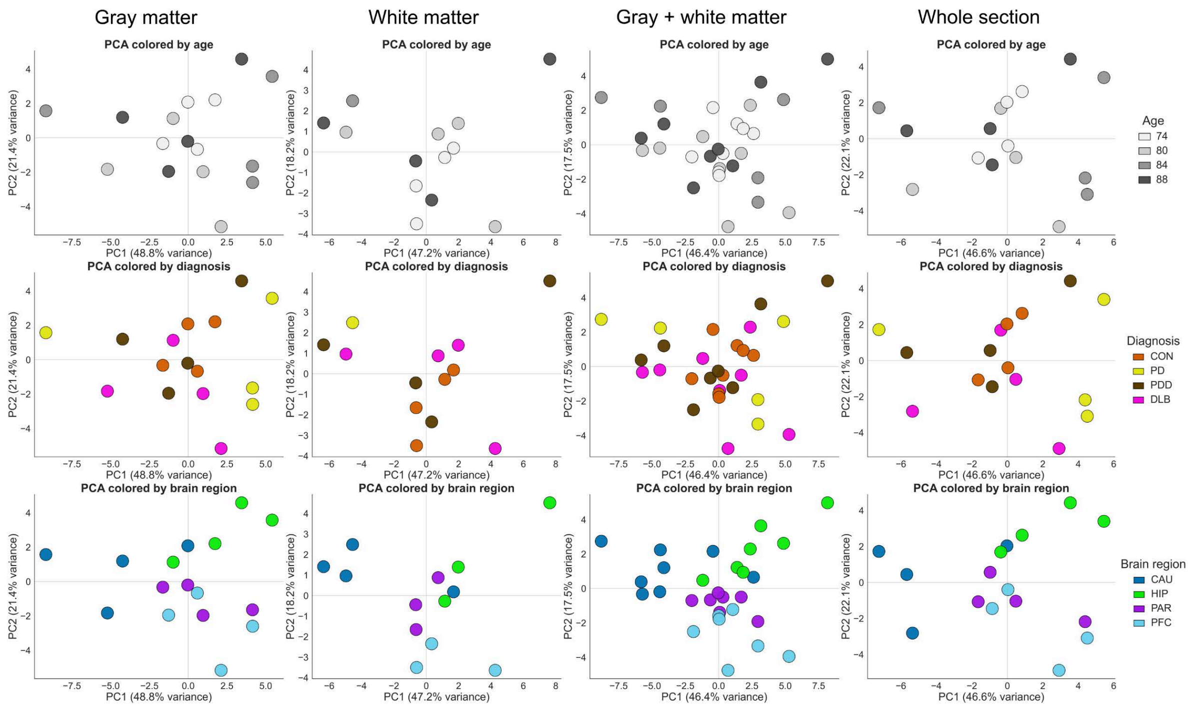


**Supplementary Figure 16. Principal component analysis (PCA) of mean protein-marker intensities within autofluorescent particle masks.** PCA was performed using the mean protein marker intensities measured within segmented autofluorescent particles. PCA plots are colored by age, diagnosis, and brain region (top to bottom) and shown separately for gray matter, white matter, combined gray and white matter, and whole tissue sections (left to right). Sample size: *n* (CAU) = 4, *n* (HIP) = 4, *n* (PAR) = 4, *n* (PFC) = 4.

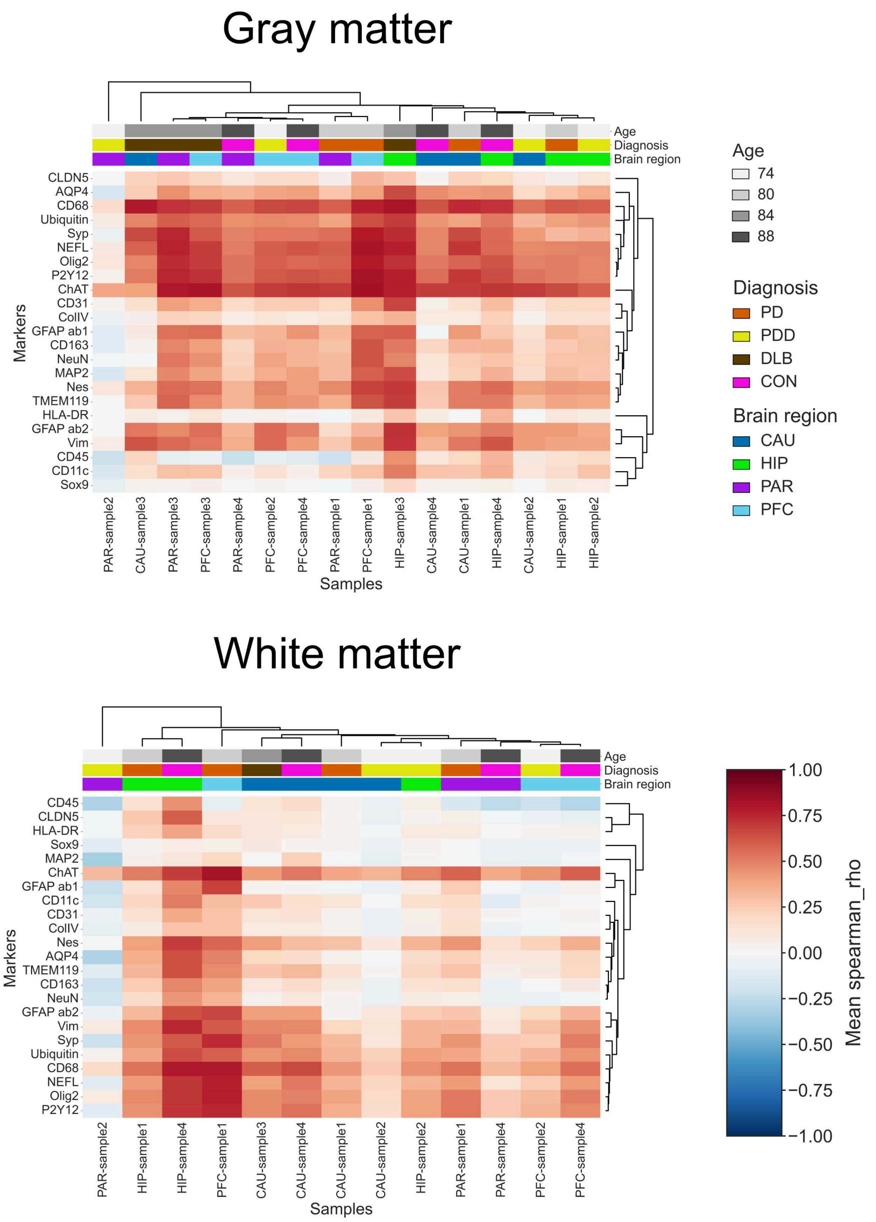


**Supplementary Figure 17. Pixel-wise correlations between 488 nm autofluorescence (AF) intensity and protein marker intensities**. Clustered heatmaps showing mean Spearman correlation coefficients between AF intensity in the 488 nm channel and protein marker intensities across selected ROIs in gray matter (top) and white matter (bottom). For each sample, pixel-wise correlations were calculated between the pre-photobleaching AF image and the corresponding multiplexed immunofluorescence images following image alignment. Columns represent individual donor samples, whereas rows represent markers included in the 28-plex antibody panel. Sample annotations include donor age, diagnosis, and brain region. Color intensity represents the mean Spearman correlation coefficient, where positive values indicate spatial co-localization and negative values indicate spatial segregation between AF and protein marker signals. Sample size: *n* (CAU) = 4, *n* (HIP) = 4, *n* (PAR) = 4, *n* (PFC) = 4.

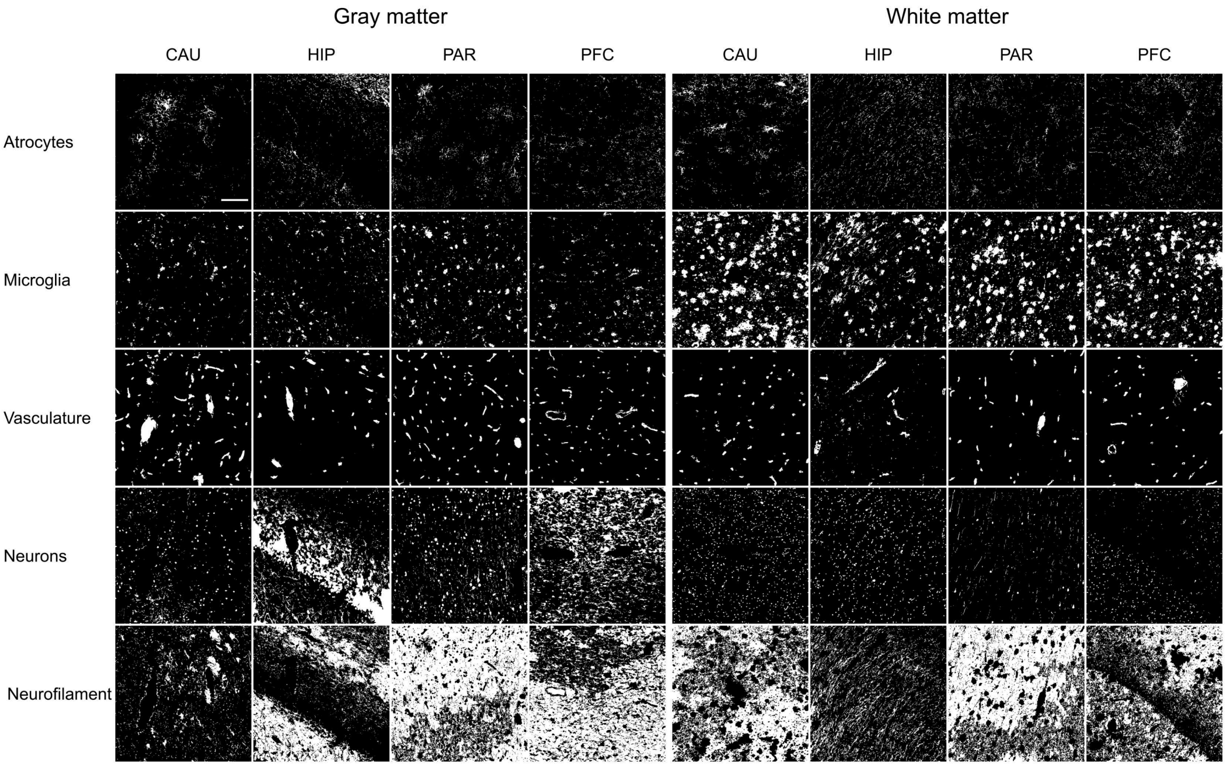


**Supplementary Figure 18. Representative binary masks for major cell types/tissue structures in gray and white matter of each analyzed brain region.** Binary masks are shown for gray matter and white matter from caudate–putamen, hippocampus, parietal cortex, and prefrontal cortex. Masks were generated by intensity thresholding followed by marker-based hierarchical exclusion to define astrocytes, microglia, neurons, neurofilaments, vasculature, and extracellular regions. Scale bar: 150 µm

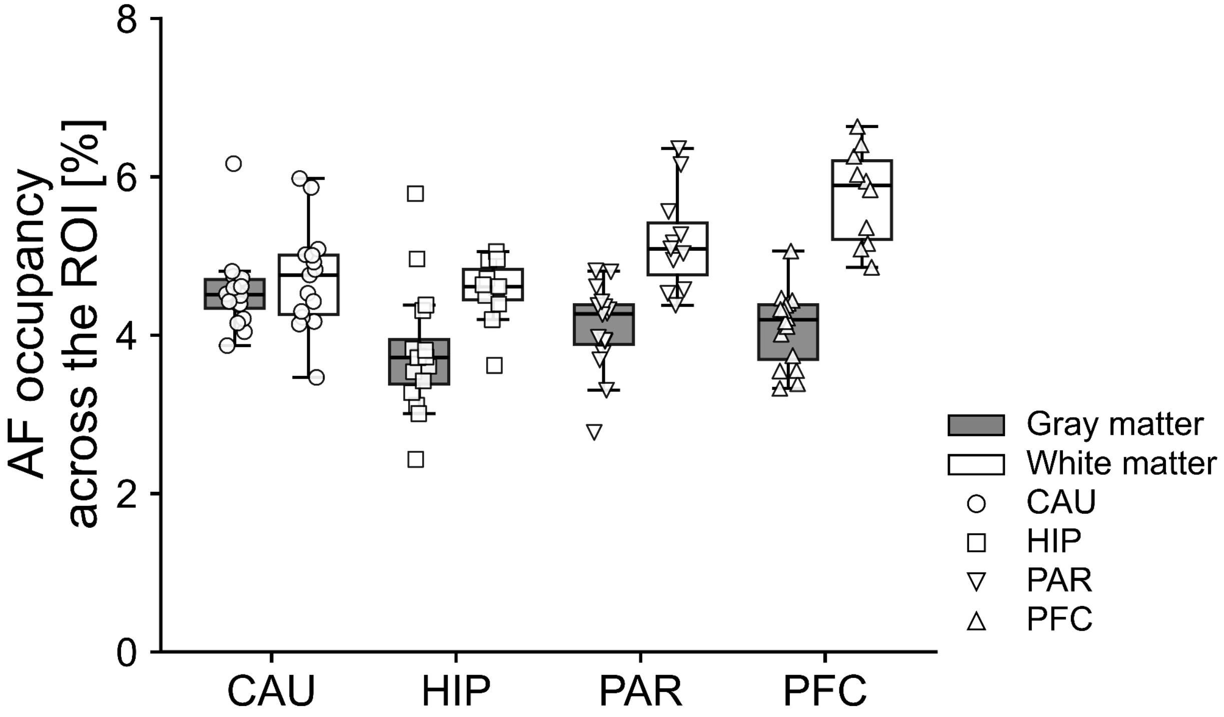


**Supplementary Figure 19. Autofluorescent (AF) particle coverage across analyzed regions of interest (ROIs) in gray and white matter.** Boxplots showing the average percentage of each ROI occupied by AF particle masks in gray matter and white matter across caudate-putamen (CAU), hippocampus (HIP), parietal cortex (PAR), and prefrontal cortex (PFC). Each point represents an individual ROI, with up to four ROIs per tissue compartment and sample. Boxplots indicate the median and interquartile range with whiskers representing the range. Gray matter is shown in gray and white matter in white. Sample size: *n* (CAU) = 4, *n* (HIP) = 4, *n* (PAR) = 4, *n* (PFC) = 4.

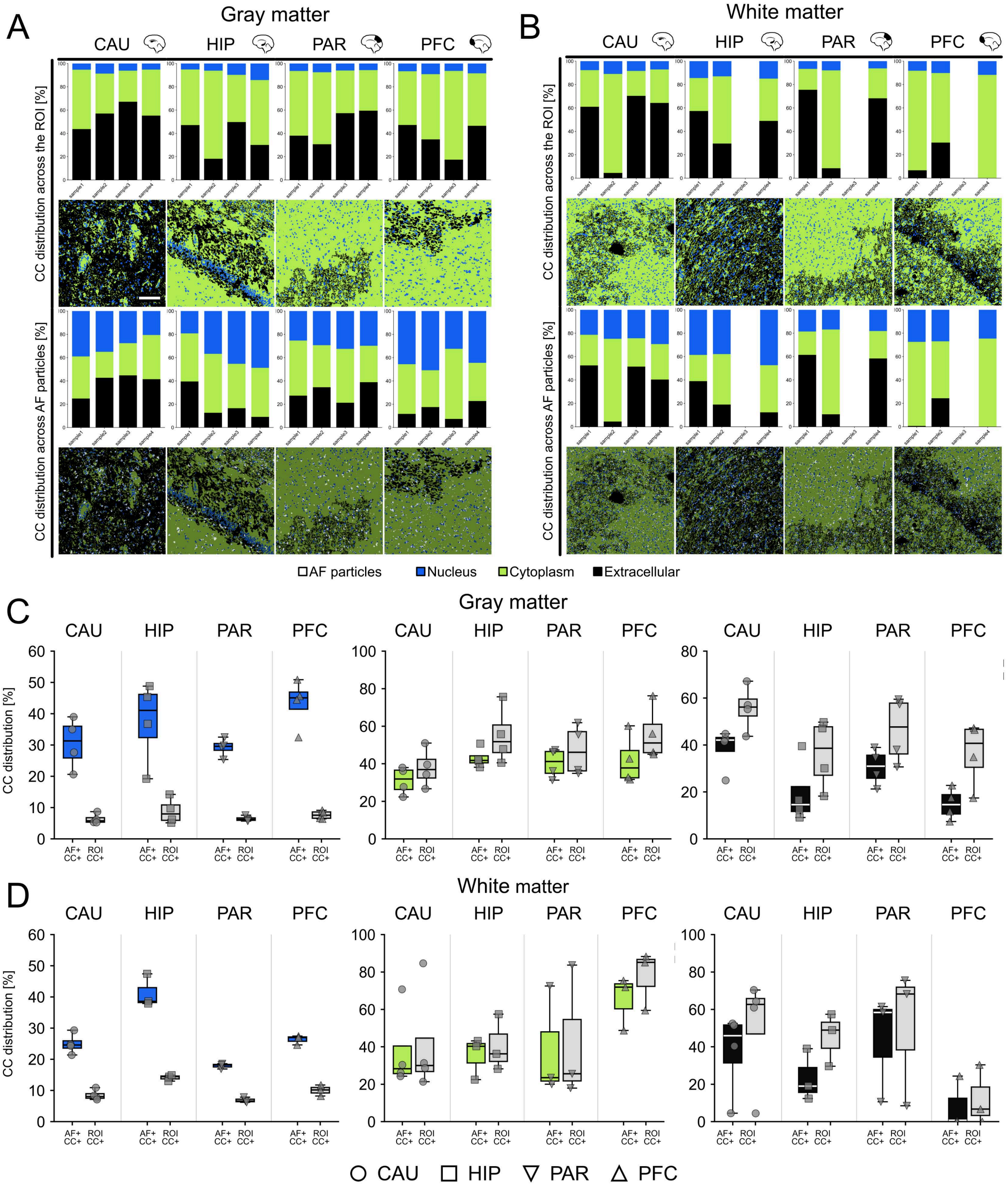


**Supplementary Figure 20. Distribution of autofluorescence (AF)-positive pixels across cellular compartments relative to compartment abundance within the analyzed ROIs.** The fraction of the AF mask overlapping each cellular compartment (AF+ CC+) was compared with the fraction of the total ROI occupied by the same cellular compartment (ROI CC+). Results are shown for gray matter (**A, C**) and white matter (**B, D**). Nuclei are shown in blue, cytoplasm in green, and extracellular space in black.

**A–B.** Stacked bar charts showing, for each sample, the relative abundance of each cellular compartment within the ROI (ROI CC+, top) and the the distribution of AF-positive pixels across these compartments (AF+ CC+, bottom), together with representative binary mask images. Samples are grouped by brain region and shown separately for gray matter (**A**) and white matter (**B**).
**C–D.** Boxplots comparing AF distribution across cellular compartments (AF+ CC+) with the relative abundance of the corresponding compartments within the ROI (ROI CC+, in gray), grouped by brain region, in gray matter (**C**) and white matter (**D**). Each point represents an individual sample, with point shape indicating brain region.
Scale bar: 150 µm. Sample size: *n* (CAU) = 4, *n* (HIP) = 4, *n* (PAR) = 4, *n* (PFC) = 4.

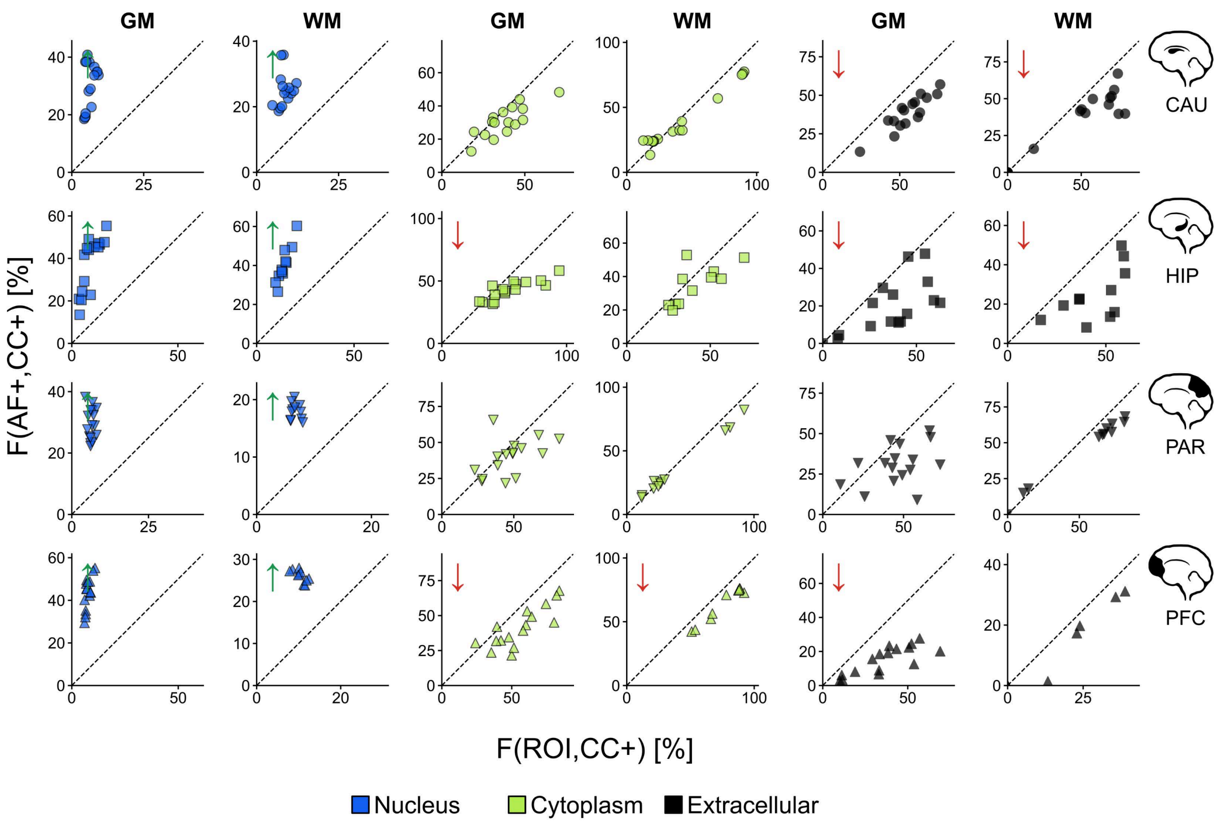


**Supplementary Figure 21. Relationship between autofluorescence distribution across cellular compartments and compartment abundance.** Scatterplots showing the relationship between the fraction of AF particle mask overlapping each cellular compartment and the relative abundance of the corresponding compartment within the region of interest (ROI). Data are shown separately for gray matter (GM) and white matter (WM) and stratified by brain region, as indicated on the right. Green upward arrows indicate significant enrichment, while red downward arrows indicate significant depletion, based on two-sided linear mixed-effect models ($p \leq 0.05$).

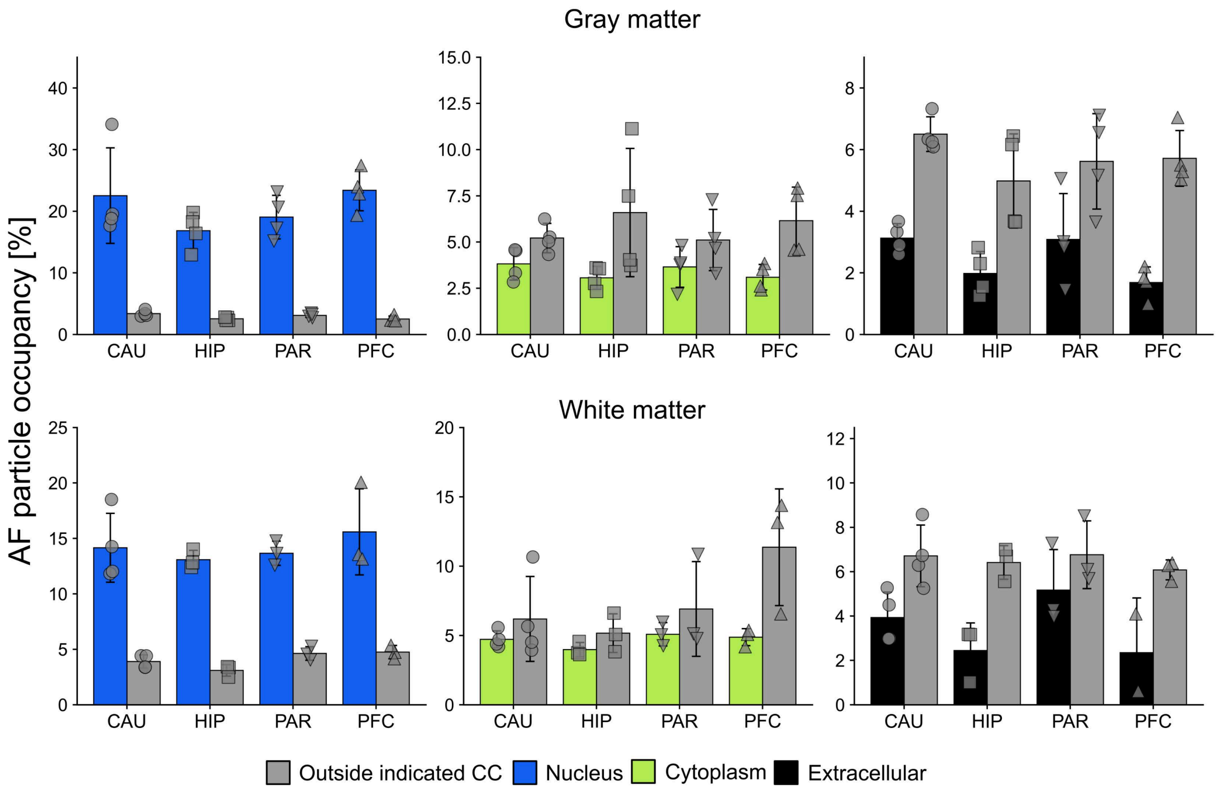


**Supplementary Figure 22. Autofluorescent (AF) particle occupancy within cellular compartments compared with AF occupancy outside each CC in the ROI.** Bar charts comparing the fraction of each cellular compartment occupied by AF-positive pixels (AF+ CC+) with the fraction of the ROI outside that CC occupied by AF-positive pixels (AF+ CC−, in gray). Data are grouped by brain region and shown separately for gray matter (top) and white matter (bottom). Each point represents an individual sample, and point shape indicates the brain region. Sample size: *n* (CAU) = 4, *n* (HIP) = 4, *n* (PAR) = 4, *n* (PFC) = 4.

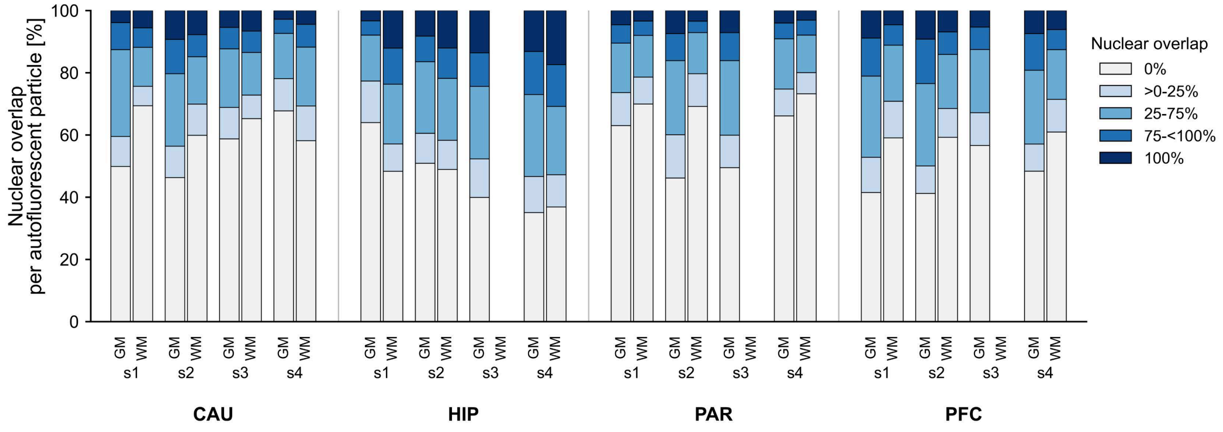


**Supplementary Figure 23. Fraction of individual AF particles overlapping with the nuclear mask.** Nuclear overlap was quantified separately for each individual AF particle to assess its spatial relationship with the nuclear compartment. The stacked bar chart shows the proportion of AF particles assigned to five nuclear-overlap categories: 0%, >0–25%, 25–75%, 75–<100%, and 100% overlap with the nuclear mask. Values represent averages across all analyzed ROIs within each sample.

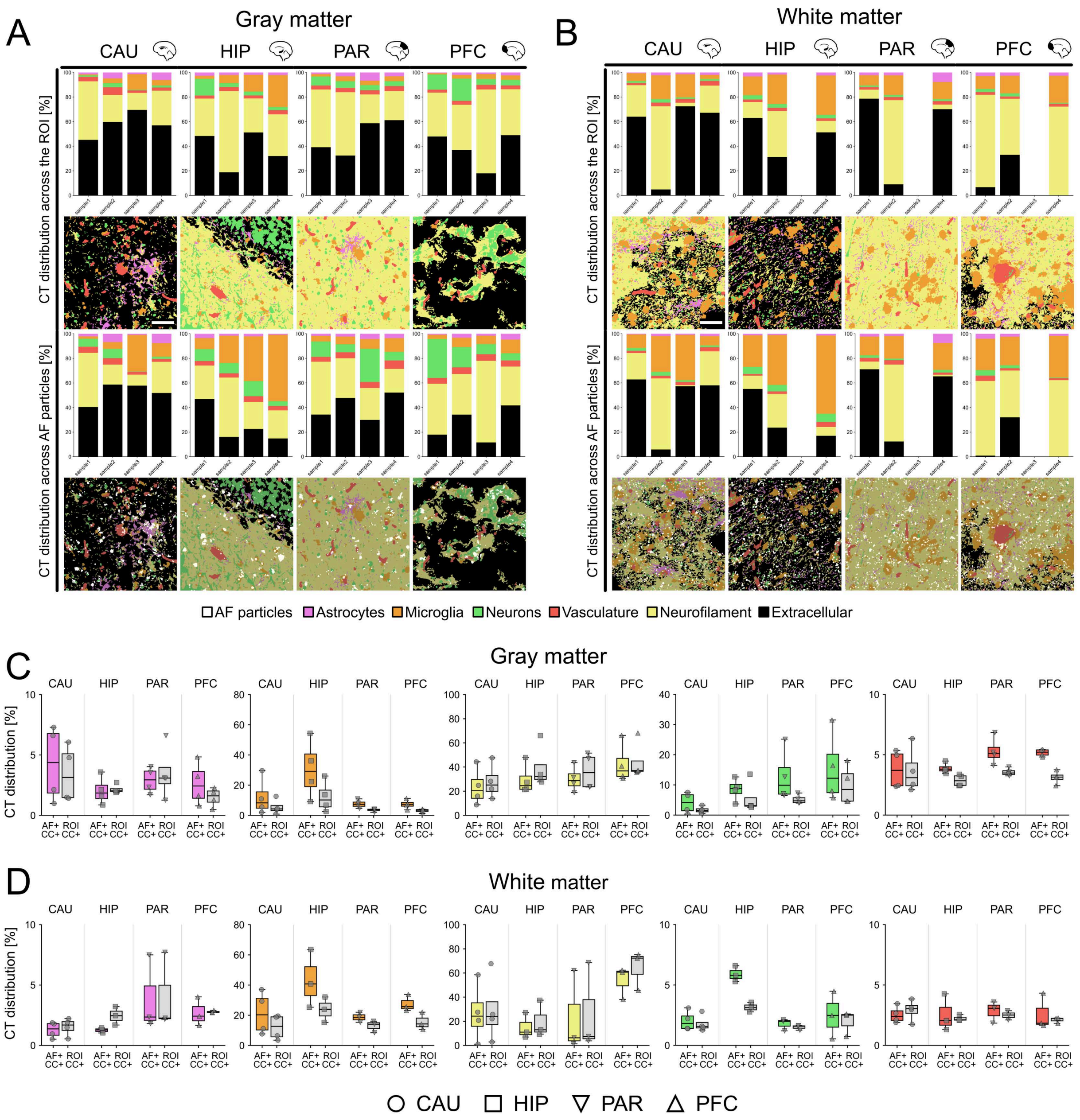


**Supplementary Figure 24. Distribution of autofluorescent (AF) pixels across cell types/tissue structures relative to cell type abundance.** The fraction of the AF particle mask overlapping each cell type (AF+ CT+) was compared with the fraction of the total ROI occupied by that cell type (ROI CT+). Results are shown for gray matter (**A, C**) and white matter (**B, D**). Astrocytes are shown in magenta, microglia in orange, neurons in green, neurofilaments in yellow, vasculature in red and extracellular space in black.

**A–B.** Stacked bar charts showing, for each sample, the relative abundance of each cell type within the ROI (ROI CT+, top) and the distribution of AF-positive pixels across these cell type masks (AF+ CT+, bottom), together with representative binary mask images. Samples are grouped by brain region and shown separately for gray matter (**A**) and white matter (**B**).

**C–D.** Boxplots comparing AF distribution across cell types (AF+ CT+) with the relative abundance of the corresponding cell types within the ROI (ROI CT+, gray), grouped by brain

region, in gray matter (**C**) and white matter (**D**). Each point represents an individual sample, with point shape indicating brain region.
Sample size: *n* (CAU) = 4, *n* (HIP) = 4, *n* (PAR) = 4, *n* (PFC) = 4.

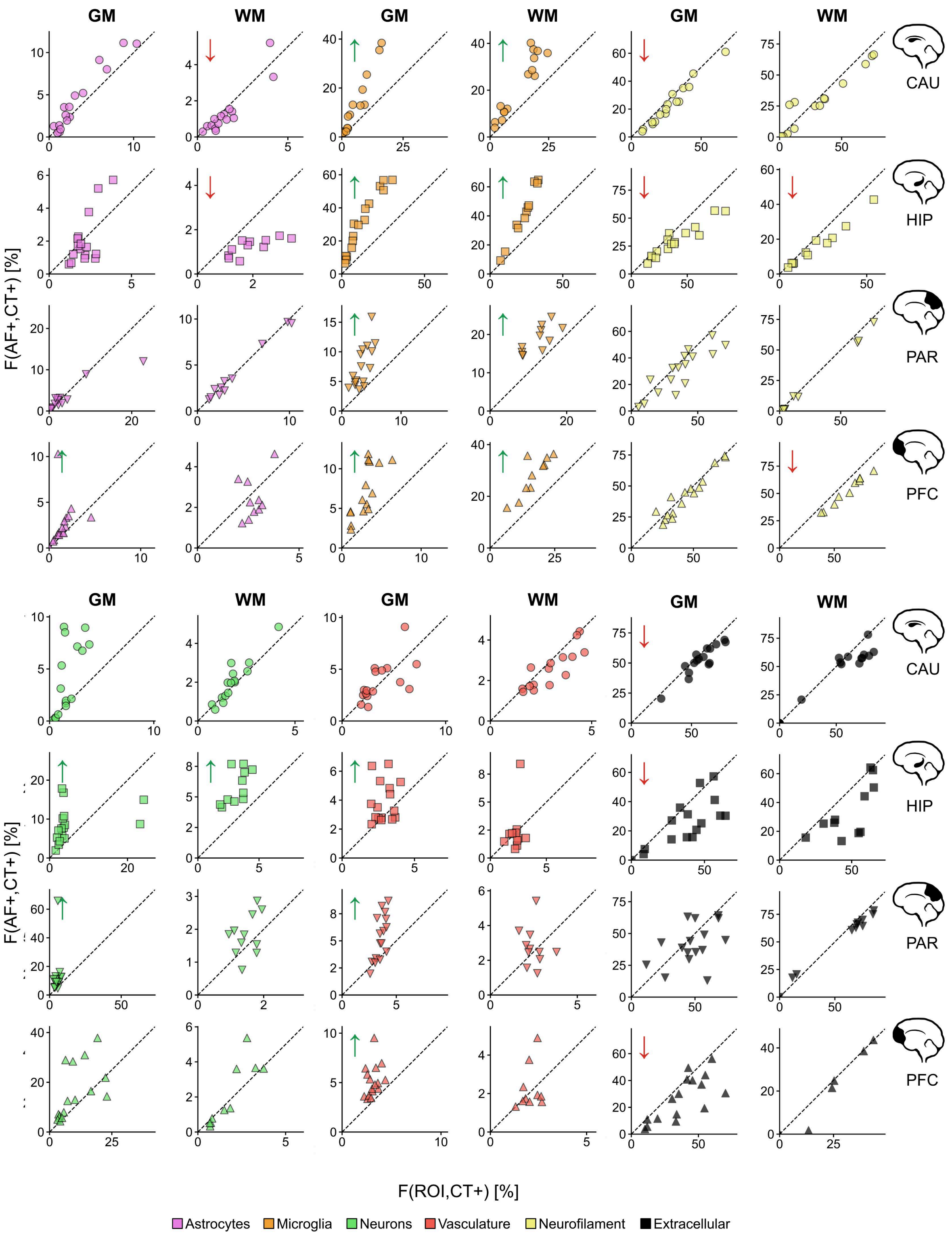


**Supplementary Figure 25. Relationship between autofluorescence distribution across cell types/tissue structures and their relative abundance.** Scatterplots showing the relationship between the fraction of AF particle mask overlapping each cell type/tissue structure and the relative abundance of the corresponding cell type/tissue structure within the ROI. Data are shown separately for gray matter (GM) and white matter (WM) and stratified by brain region, as indicated on the right. Green upward arrows indicate significant enrichment, while red downward arrows indicate significant depletion, based on two-sided linear mixed-effect models ($p \leq 0.05$).

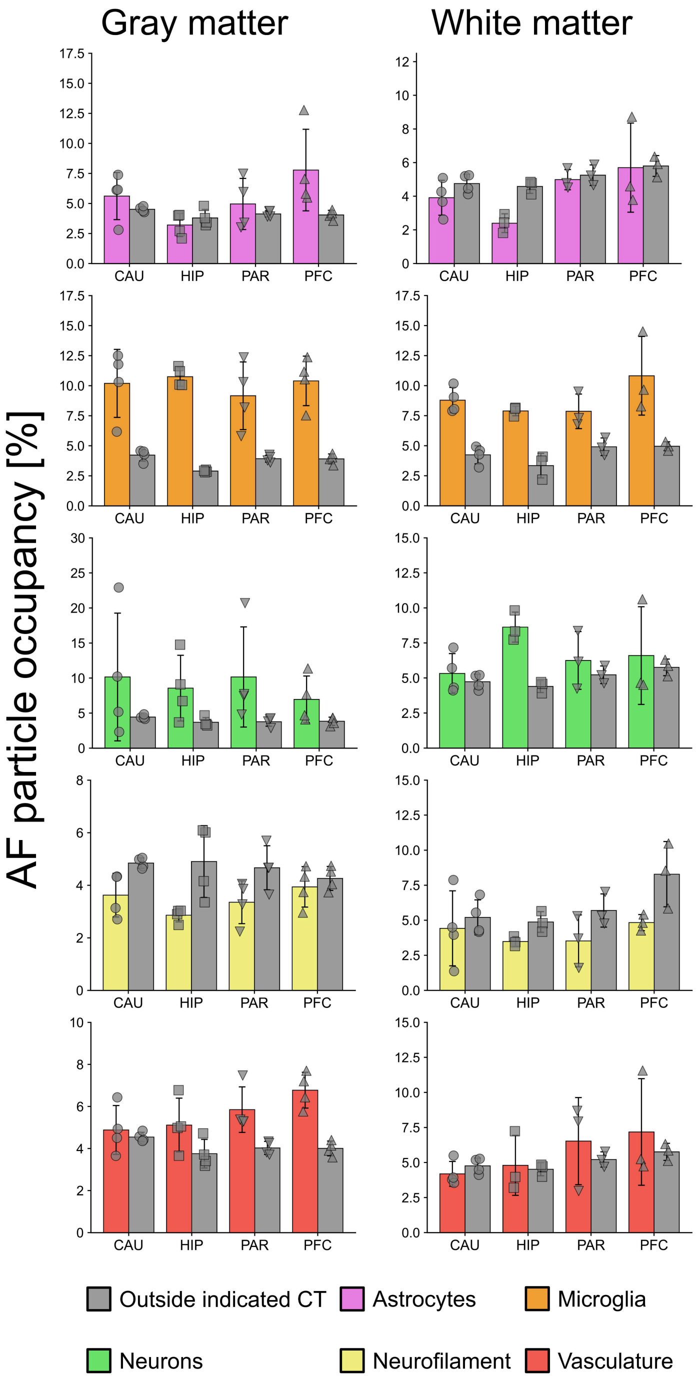


**Supplementary Figure 26. Autofluorescent (AF) particle occupancy within cell types/tissue structures (CT) compared with AF occupancy outside each CT in the ROI.** Bar charts comparing the fraction of each cell type/tissue structure (CT) occupied by AF-positive pixels (AF+ CT+) with the fraction of the ROI outside that CT occupied by AF-positive pixels (AF+ CT−). Data are grouped by brain region and shown for gray matter (left) and white matter (right). Each point represents an individual sample, with point shape indicating brain region. Sample size: *n* (CAU) = 4, *n* (HIP) = 4, *n* (PAR) = 4, *n* (PFC) = 4.

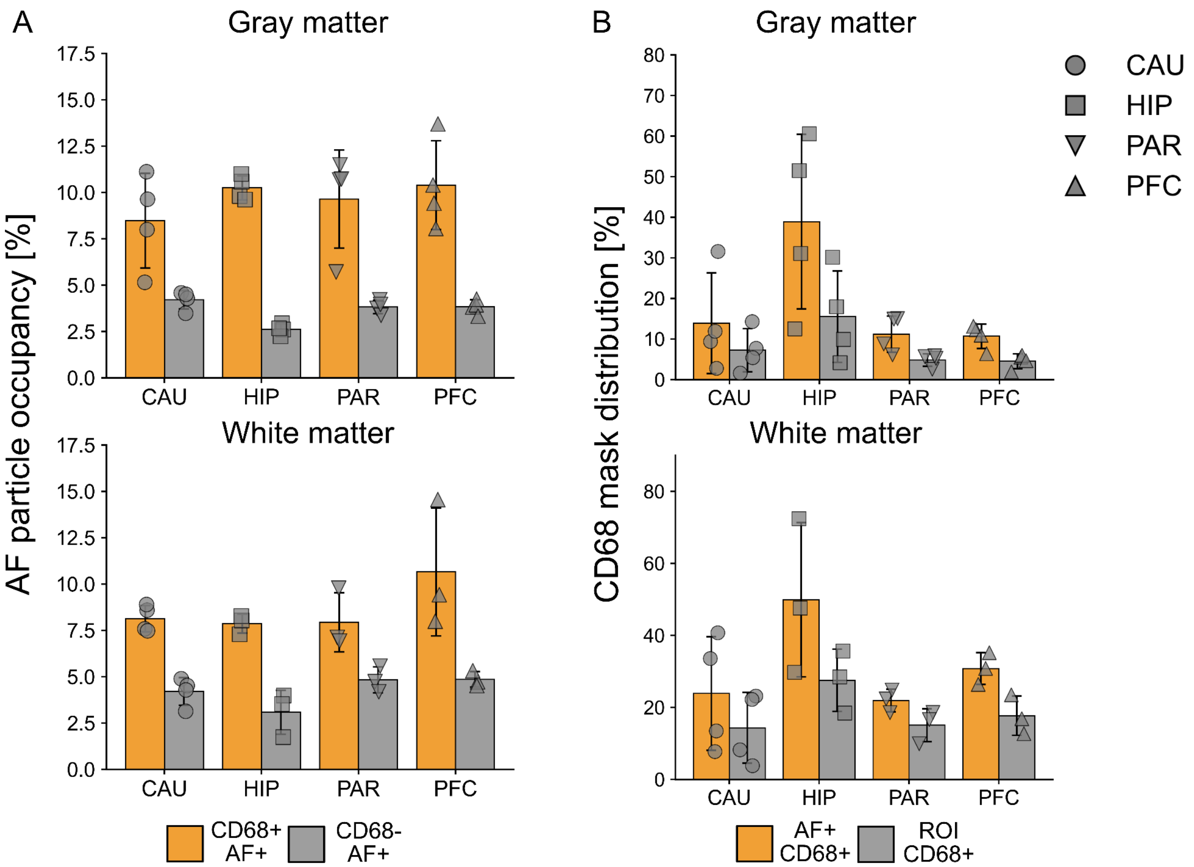


**Supplementary Figure 27. Spatial association between CD68-positive regions and autofluorescent particles.**
**A.** Bar charts comparing the fraction of AF particle mask overlapping the CD68 mask with the fraction of the ROI occupied by CD68-positive pixels across the four analyzed brain regions.
**B.** Bar charts comparing the fraction of the CD68 mask occupied by AF-positive pixels compared with the fraction of the surrounding ROI outside the CD68 mask occupied by AF-positive pixels. Each point represents an individual sample, with point shape indicating the brain region. Sample size: *n* (CAU) = 4, *n* (HIP) = 4, *n* (PAR) = 4, *n* (PFC) = 4.
